# Illuminating protein space with a programmable generative model

**DOI:** 10.1101/2022.12.01.518682

**Authors:** John Ingraham, Max Baranov, Zak Costello, Vincent Frappier, Ahmed Ismail, Shan Tie, Wujie Wang, Vincent Xue, Fritz Obermeyer, Andrew Beam, Gevorg Grigoryan

## Abstract

Three billion years of evolution have produced a tremendous diversity of protein molecules, and yet the full potential of this molecular class is likely far greater. Accessing this potential has been challenging for computation and experiments because the space of possible protein molecules is much larger than the space of those likely to host function. Here we introduce Chroma, a generative model for proteins and protein complexes that can directly sample novel protein structures and sequences and that can be conditioned to steer the generative process towards desired properties and functions. To enable this, we introduce a diffusion process that respects the conformational statistics of polymer ensembles, an efficient neural architecture for molecular systems based on random graph neural networks that enables long-range reasoning with sub-quadratic scaling, equivariant layers for efficiently synthesizing 3D structures of proteins from predicted inter-residue geometries, and a general low-temperature sampling algorithm for diffusion models. We suggest that Chroma can effectively realize protein design as Bayesian inference under external constraints, which can involve symmetries, substructure, shape, semantics, and even natural language prompts. With this unified approach, we hope to accelerate the prospect of programming protein matter for human health, materials science, and synthetic biology.

## Introduction

Protein molecules carry out most of the biological functions necessary for life, but inventing them is a complicated task that has taken evolution millions to billions of years. The field of computational protein design aims to shortcut this by automating the design of proteins for desired functions in a manner that is *programmable.* While there has been significant progress towards this goal over the past three decades [Kuhlman and Bradley, 2019, Huang et al., 2016], including the design of novel topologies, assemblies, binders, catalysts, and materials [Koga et al., 2012, Cao et al., 2022, Kries et al., 2013, Joh et al., 2014], most *de novo* designs have yet to approach the complexity and variety of macromolecules that are found in nature, which possess complex and asymmetric layered architectures built from many distinct sub-domains. Reasons for this include that (i) modeling the relationship between sequence, structure, and function is difficult, and (ii) most computational design methods rely on iterative search and sampling processes which, just like evolution, must navigate a rugged fitness landscape incrementally Maynard Smith [1970]. While many computational techniques have been developed to accelerate this search [Huang et al., 2016] and to improve the prediction of natural protein structures [Jumper et al., 2021], the space of *possible* proteins remains combinatorially large and only partially accessible by traditional computational methods. Determining how to efficiently explore the space of designable protein structures while also biasing towards specific functions remains an open challenge.

An alternative and potentially appealing approach to protein design would be to directly sample from the space of proteins that are compatible with a set of desired functions. While this could address the fundamental limitation of iterative search methods, it would require an effective parameterization of a-priori “plausible” protein space, a way to draw samples from this space, and a way to bias this sampling towards desired properties and functions. Deep generative models have proven successful in solving these kinds of high-dimensional modeling and inference problems in other domains, for example, in the text-conditioned generation of photorealistic images [Ramesh et al., 2021, 2022, Saharia et al., 2022]. For this reason, there has been considerable work developing generative models of protein space, applied to both protein sequences [Riesselman et al., 2018, Greener et al., 2018, Ingraham et al., 2019, Anand et al., 2022, Madani et al., 2020, Rives et al., 2021, Notin et al., 2022] and structures [Anand and Huang, 2018, Lin et al., 2021, Eguchi et al., 2022, Anand and Achim, 2022, Trippe et al., 2022, Wu et al., 2022a].

Despite these recent advances in generative models for proteins, we argue that there are three desiderata that have yet to be realized simultaneously in one system. These are (i) to jointly model the 3D structures and sequences of *full protein complexes*, (ii) to do so with computation that scales *sub-quadratically* with system size, and (iii) to enable *conditional sampling* under diverse cues without re-training. The first, generating full complexes, is important because protein function is often interpretable only in the context of a bound complex. The second, sub-quadratic scaling of computation, is important because it has been an essential ingredient for managing complexity in other modeling disciplines, such as in computer vision, where convolutional neural networks scale linearly with the number of pixels in an image, and in computational physics, where fast *N*-body methods are used for efficient simulation of everything from stellar to molecular systems Barnes and Hut [1986]. And lastly, the requirement to sample *conditionally* from a model without having to retrain it on new target functions is of significant interest because protein design projects often involve many complex and composite requirements which may vary over time.

Here we introduce Chroma, a generative model for proteins that achieves all three of these redifferent properties suchquirements by modeling full complexes with quasi-linear computational scaling and by admitting arbitrary conditional sampling at generation time. It builds on the framework of diffusion models Sohl-Dickstein et al. [2015], Song et al. [2021], which model high-dimensional distributions by gradually transforming them into simple distributions and learning to reverse this process, and of graph neural networks Gilmer et al. [2017], Battaglia et al. [2018], which can efficiently reason over complex molecular systems. We show that it produces high-quality, diverse, novel, and designable structures, and that it enables *programmable* generation of proteins conditioned on several different properties such as symmetry, shape, protein class, and even textual input. We anticipate that scalable generative models like Chroma will enable a widespread and rapid increase in our ability to design and build protein systems fit for function.

## Results

### A scalable generative model for protein systems

Chroma achieves high-fidelity and efficient generation of proteins by introducing a new diffusion process, neural network architecture, and sampling algorithm based on principles from contemporary generative modeling and biophysical knowledge. Diffusion models generate data by learning to reverse a noising process, which for previous image modeling applications has typically been uncorrelated Gaussian noise. In contrast, our model learns to reverse a *correlated* noise process to match the empirical covariance structure in real proteins which is dominated by the constraints of a collapsed polymer to be chain structured with a particular radius of gyration (Fig. 1a, Appendix C). Prior models for protein structure have typically leveraged computation that scales as 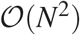 [Trippe et al., 2022, Wu et al., 2022a] or 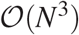 [Jumper et al., 2021, Anand and Achim, 2022] in the number of residues *N*, which has limited their application to small systems or required large amounts of computation for modestly sized systems. To overcome this, Chroma introduces a novel neural network architecture (Fig. 1b) for processing and updating molecular coordinates that uses random long range graph connections with connectivity statistics inspired by fast *N*-body methods [Barnes and Hut, 1986] and that scales sub-quadratically 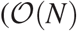 or 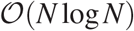, Appendix D). Finally, we also introduce methods for principled *low-temperature sampling* from diffusion models with a modified diffusion process that allows us to increase quality of sampled backbones (increasing likelihood) while reducing conformational diversity (reducing entropy). A *design network* then generates sequence and side-chain conformations conditioned on the sampled backbone, yielding a joint generative process for the full sequence and the 3D positions of all heavy atoms in a protein complex. The design network is based on the same graph neural network as the backbone network, but with conditional sequence decoding layers and side-chain decoding layers that are similar to prior works [Ingraham et al., 2019, Anand et al., 2022] and have recently seen further refinement and experimental validation [Jing et al., 2020, Hsu et al., 2022, Dauparas et al., 2022]. While it is also possible to model sequence and side chain degrees of freedom as part of a joint diffusion [Hoogeboom et al., 2021], we found that a sequential factorization is effective while being considerably more efficient.

**Figure 1:**
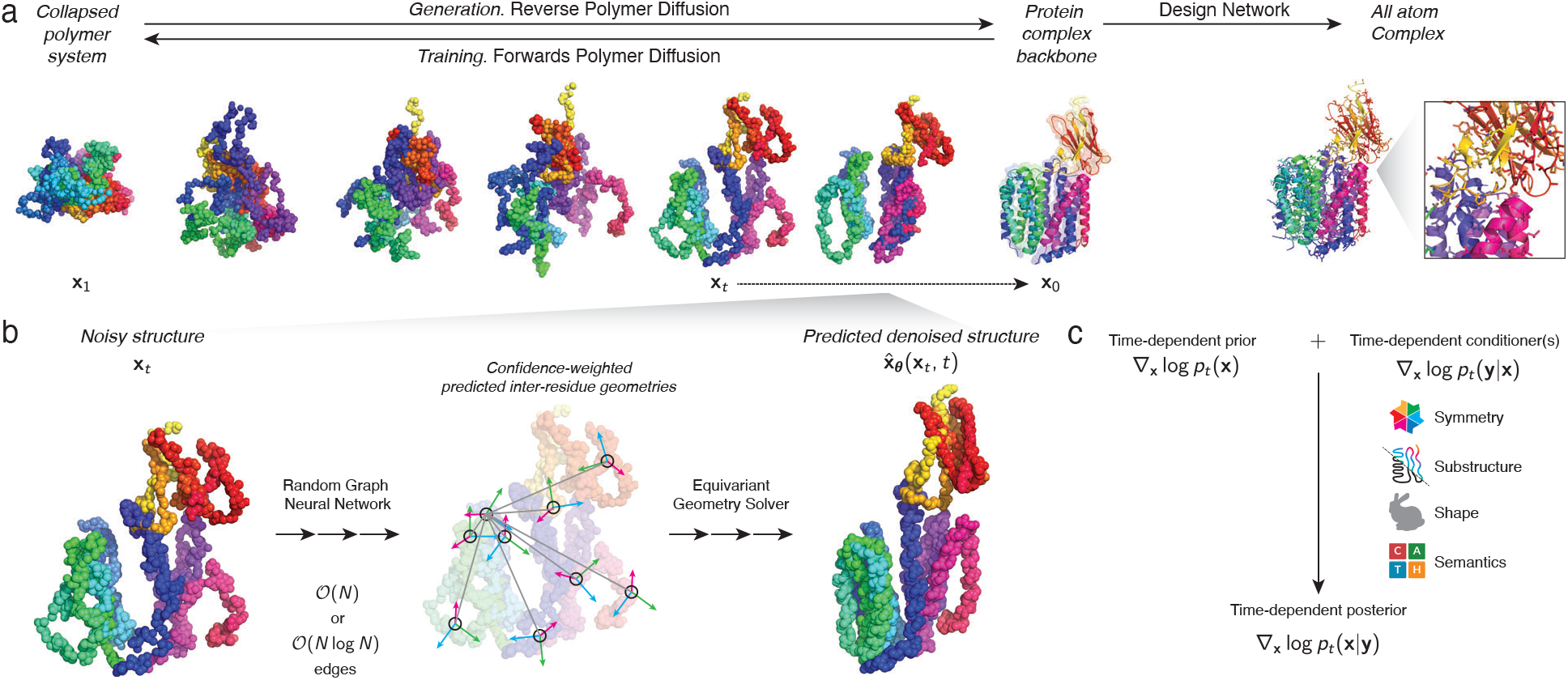
*Chroma* is a generative model for proteins and protein complexes that combines a structured diffusion model for protein backbones with scalable molecular neural networks for backbone synthesis and all-atom design. **a**, A correlated diffusion process with chain and radius of gyration constraints gradually transforms protein structures into random collapsed polymers (right to left). The reverse process (left to right) can be expressed in terms of a time-dependent optimal denoiser 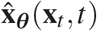 (**b**), which we parameterize in terms of a random graph neural network with long-range connectivity inspired by efficient *N*-body algorithms (**b:**, middle) and a fast method for solving for a global consensus structure given predicted inter-residue geometries (**b:**, right). (**a**, top right) Another graph-based *design network* generates protein sequences and side-chain conformations conditionally based on the sampled backbone. **c**, The time-dependent protein prior learned inside of the diffusion model can be combined with auxiliary conditioning information for programmable generation of protein systems.

An important aspect of our diffusion-based framework is that it enables *conditional* sampling under combinations of user-specified constraints. This is made possible by a key property of diffusion models: they can recast the diffusion process for a target conditional distribution *p*(**x**|**y**) of proteins **x** given constraints **y**, as a combination of the learned gradient field from the diffusion model ∇_**x**_ log*p_t_* (**x**) and gradients from external classifiers that have been trained to predict labels **y** from noisy examples of **x**, i.e. ∇_**x**_log*p_t_*(**y**|**x**) [Song et al., 2021] (Fig. 1c, Appendix B). This means that any classifier that predicts a protein property (e.g. *p_t_* (**y**|**x**)) from structure can be repurposed to guide the diffusion process towards proteins with those properties. To demonstrate and explore the extent of this programmatic conditioning, we introduce a variety of analytic and learned conditioners *p_t_*(**y**|**x**) (Fig. 1c, Appendix I). This includes geometrical constraints, which are typically analytic and include constraints on distance (Appendix J), substructure root mean-squared deviation (RMSD) from a target substructure (Appendix K), symmetric complexes under arbitrary symmetry groups (Appendix L), and shape matching to arbitrary point clouds (Appendix M). We also explore the possibilities of *semantic* prompting by training graph neural networks to predict multi-scale classifications (Appendix N) and natural language annotations (Appendix O) directly from protein structures. Any subset of all of these constraints may then be combined for bespoke, on-demand protein generation subject to problem-specific desiderata.

### Analysis of unconditional samples

We first sought to characterize the diffusion model for protein backbone structures by analyzing a large set of unconditional samples of proteins and protein complexes. When initially exploring unconditional samples using the diffusion model, we observed an interesting phenomenon where the model assigned high likelihoods to natural structures but still produced samples that usually were mostly unstructured with little backbone hydrogen bonding and secondary structure content (Appendix B). We reasoned that this phenomenon was analogous to a common issue with likelihood-based generative models such as language models [Holtzman et al., 2020] and diffusion models [Dhariwal and Nichol, 2021], where it is typical for models to take longer to eliminate the probability mass for poorly structured states (of which there are usually exponentially many more) than it does for models to assign high probabilities to well-structured states (e.g. well-formed sentences and images). The standard solution to this issue of *overdispersion* of likelihood-based generative models is to leverage modified sampling procedures which bias towards higher probability states, such as beam-search or greedy decoding in lanaguage models [Holtzman et al., 2020] or *classifier guidance* and *classifier-free guidance* for conditional diffusion models [Ho and Salimans, 2021]. The latter methods of classifier guidance heavily rely on strong classifiers to be effective, and we found them insufficient to improve sample quality of our backbones. Instead, we developed a novel and general sampling algorithm for diffusion models that enables sampling from the temperature-perturbed distribution 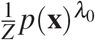 (Appendix B). Increasing the inverse temperature parameter *λ*_0_ redistributes probability mass towards higher likelihood states. We emphasize that these low-temperature sampling methods are of critical importance to our framework, and throughout the paper we sample backbones at inverse temperatures ranging from *λ*_0_ = 16 to *λ*_0_ = 1 (Appendix H).

Equipped with our low-temperature sampling method, we characterized a large number of samples from the prior and to compare their structural statistics to data from the Protein Data Bank (PDB). We sampled 50,000 single-chain backbones and 10,000 complex backbones at inverse temperature *λ*_0_ = 10. As can be seen in Fig. 2a, the unconditional samples display many properties shared by natural proteins, such as complex layering of bundled alpha helices and beta sheets in cooperative unknotted folds. We provide grids of randomly picked subsets of these samples in Appendix H (Supplementary Figs. 6 and 7 for single-chain and complex structures, respectively). To quantitatively measure the agreement with natural folds, we sampled another set 10,000 single chain samples and computed several key structural properties, including secondary structure utilization, contact order [Plaxco et al., 1998], length-dependent radius of gyration [Tanner, 2016], length-dependent long-range contact frequency and density of inter-residue contacts (Appendix H). We generally observe agreement between the distribution of these statistics for Chroma and samples from the PDB, indicateing that these backbone structures appear to be similar to native proteins (Fig. 2b and c). We do see a slight over-utilization of *α*-helices (by ~ 0.5-1.0 standard deviations), which we suspect may be a consequence of low-temperature sampling (i.e., helices are used more frequently than strands in natural proteins, but the ratio is somewhat accentuated in low-temperature samples).

**Figure 2:**
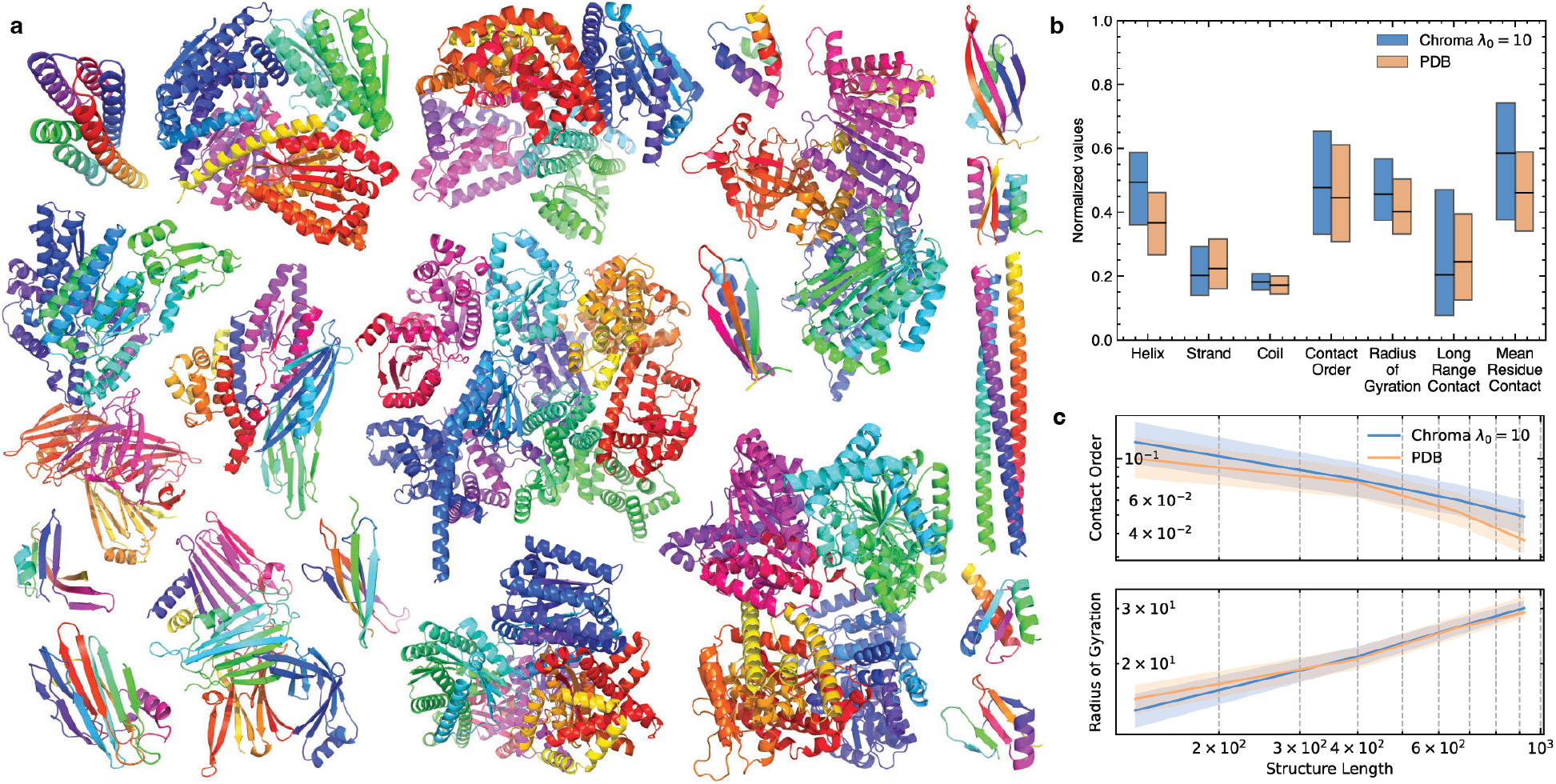
Analysis of unconditional samples reveals diverse geometries that reproduce low-level structural statistics but exhibit novel higher-order structure. **a**, A representative set of proteins and protein complexes sampled from the unconditioned backbone model at inverse temperature *λ*_0_ = 10 exhibits complex and diverse architectures with high secondary structure content. **b**, Across a set a set of 10,000 single chains, samples from Chroma have structural properties that are similar to natural protein structures from the PDB, including secondary structure utilization and length-normalized contacBt order, radius of gyration, and contact density statistics. Low-temperature samples from Chroma tend to slightly favor helices over strands and are more compact than those found in the PDB. **c**, Chroma samples reproduce length-dependent scaling of contact order [Plaxco et al., 1998] and radius of gyration.

### Evaluation of generated protein structures

An important question is whether the backbone structures generated by Chroma can be realized with sequences of natural amino acids—i.e., whether they are “designable”. While the only way to answer this question definitively is through experimental characterization, we performed two types of analyses to provide *in-silico* support for the designability of our generated structures. In the first, we created sequences for our generated backbones and assessed whether open-source structure prediction models representative of the current state of art [Wu et al., 2022b] would predict that the sequences would correctly fold into the original, generated structure (Fig. 3a). We note that this type of sequence-structure mutual consistency test rests on the ability of the structure prediction model to generalize to novel folds and topologies, which has yet to be conclusively demonstrated. Nevertheless, this evaluation is able to provide partial supporting evidence for designability in the instances where the predicted and generated structures have strong agreement, and there have been some successful applications of structure prediction for de novo design [Anishchenko et al., 2021]. Figs. 3b-d show the results of this analysis for a set of 100 backbones generated at random using Chroma with length uniformly randomly sampled in the range [100,500] (Appendix H.4). Fig. 3b shows the TM-score [Zhang and Skolnick, 2005] achieved between generated and predicted structures in this test as a function of protein length. While it is not surprising that this task is more challenging for longer proteins, as the difficulty of both generation and prediction will generally increase with chain length, it is remarkable that TM > 0.5 (a broadly utilized cutoff to indicate “the same fold”) is achieved even for proteins as long as 480 amino acids. The overall distribution of TM-scores in Fig. 3c shows the cutoff of 0.5 is achieved in 55% of cases overall.

**Figure 3:**
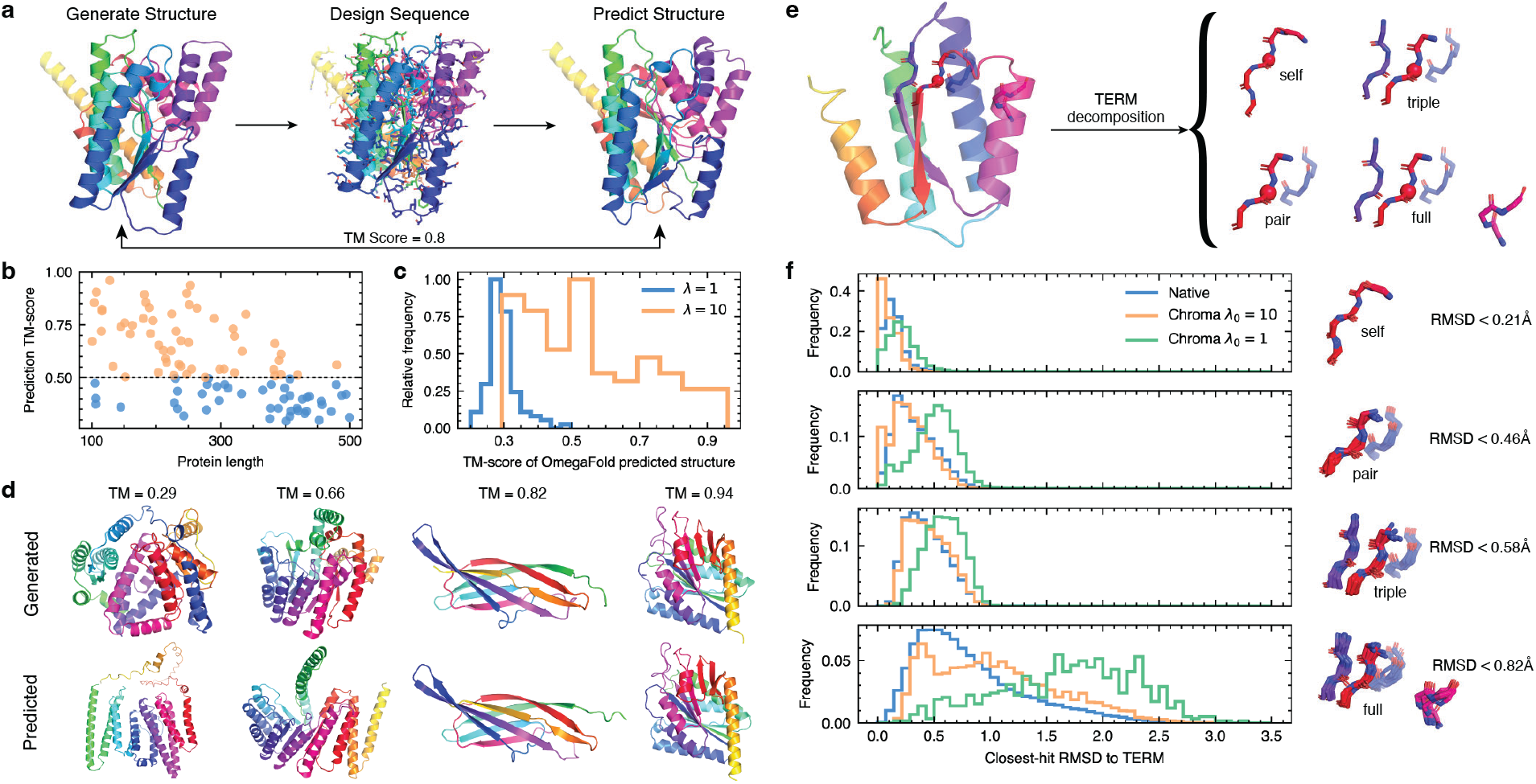
*Chroma*-generated backbones are designable by a variety of computational metrics. **a**, Workflow of the test involving generation of a backbone using Chroma, design of sequences for this backbone using our sequence design model, structure prediction for these sequences with OmegaFold [Wu et al., 2022b], and comparison of predicted versus generated structures. **b:** Resulting TM-scores (best out of 100 design attempts for each structure) vs. to protein length. **c**, Distribution of resulting TM-scores (orange). Shown in blue is the same distribution from a control calculation involving the same workflow but with Chroma sampling at inverse temperature *λ*_0_ = 1 (versus *λ*_0_ = 10 in the original test). **d**, Comparisons between generated and predicted structures for several cases spanning a range of TM-scores from the worst (left) to the best (right) observed in this test. **e**, A visual depiction of a TERMs decomposition. **f**, The distribution of closest-match RMSD for TERMs of increasing order originating from native or Chroma-generated backbones (with inverse temperature *λ*_0_ being 1 or 10).

To isolate the possible confounding effect of structure prediction, we also directly analyzed the higher-order, local backbone geometries in Chroma-generated backbones versus natural backbones. Natural protein structures exhibit considerable degeneracy in their use of local tertiary backbone geometries, such that completely unrelated proteins tend to utilize very similar structural motifs and relatively few local tertiary geometries account for the majority of the observed structures [Mackenzie et al., 2016]. These tertiary motifs, or *TERMs*, consist of a central residue, its backbone-contiguous neighbors, neighboring residues capable of contacting the central residue, and their backbone-contiguous neighbors [Mackenzie et al., 2016, Zheng et al., 2015]. Depending on how many contacting residues are combined into the motif, TERMs can be distinguished as self, pair, triple, or higher-order, corresponding to having zero, one, two, or more contacting neighbors (Fig. 3e). To compare the local geometry of Chroma-generated backbones with that of native structures, we isolated all possible self, pair, triple, and full TERMs (i.e., TERMs containing all contacting residues for a given central residue) and identified the closest neighbor (by backbone RMSD) to each within a redundancy-pruned subset of the PDB, the search database (Appendix H.5). We performed a similar analysis on a set of native proteins not contained within the search database, additionally taking care to remove any homologous matches (Appendix H.5). Figure 3f shows the distribution of closest-neighbor RMSDs for TERMs derived from both natural (native) and Chroma-sampled backbones that were generated at inverse temperatures *λ*_0_ = 10 and *λ*_0_ = 1. The distributions of nearest neighbor RMSD were very close for low-temperature samples from Chroma and native proteins, indicating that Chroma geometries are valid and likely to be as designable as native proteins, including complex motifs with four or five disjoint fragments (Fig. 3f, bottom panel). Because native amino-acid choices are driven by these local geometries [Zhou et al., 2019], and adherence to TERM statistics has been previously shown to correlate with structural model accuracy and success in *de-novo* design Zheng et al. [2015], Zhou et al. [2019], this argues for the general designability of Chroma-generated backbones in a model-independent manner. Notably, the samples from Chroma without temperature adjustment (i.e. *λ*_0_ = 1) exhibit unfolded and unstructured geometries (Fig. 3f), which underlines the importance of having a method for low-temperature sampling.

### Programmability

Being able to efficiently sample realistic proteins is necessary but not sufficient for downstream applications such as therapeutic development, since unconditional samples are unlikely to possess desired functional properties. An important aspect of Chroma is its *programmability*, which means that it is straightforward to directly bias sampling towards desired properties (Section “A scalable generative model for protein systems” and Fig. 1). This leverages a remarkable property of diffusion models, which is that they can make traditionally difficult Bayesian inversion problems tractable [Sohl-Dickstein et al., 2015, Song et al., 2021]. Specifically, to condition on a property or set of properties (collection of events) **y**, it is sufficient to train a time-dependent classifier model *p_t_* (**y**|**x**) on the noised structures **x**_*t*_ ~ *p_t_* (**x**|**x_0_**) and to adjust the sampling process by ∇_*x*_ log *p_t_* (**y**|**x**), the gradient of the log likelihood that structure **x_0_** will have the desired property at time *t* = 0 (the end of the reverse diffusion). Depending on the property in question, this probability can be expressed analytically in a closed form (e.g., see Appendix J), as an empirical, analytic approximation (e.g., see Appendix K), or as the prediction of a neural network (e.g., see Appendix N and O). This conditioning formalism is very natural for protein design as it decouples the problem of parameterizing the space of likely structured proteins, the cost of which is amortized by training a strong prior model once, from the problem of expressing the correct determinants of the desired function, which can be isolated to the classifier model. Thus, by focusing effort on building the right classifiers, designers can spend most of their time on solving specific functional objectives.

To demonstrate what may be achievable with conditional generation, we built several *p_t_* (**y**|**x**) classifier models (Appendix I), including those where **y** encoded: (i) a distance-based constraint (e.g., a “contact” between residues), (ii) the presence of a disjoint sub-structure (based on backbone RMSD), either anywhere in the generated structure or in a pre-specified alignment, (iii) various residue-, domain-, and complex-level classes (e.g., CATH or PFAM domains, secondary structure labels, interfacial residues), and (iv) natural language annotations trained on protein captions from the PDB and Uniprot [Consortium, 2020]. While we believe that each of these classifiers represents only a preliminary realization of these conditioning modes, we already see that they suggest tremendous possibilities for *programmable* design.

We begin by considering analytic conditioners that can control protein backbone geometry. We found that conditioning on the symmetry of protein complexes, which can also be cast as sampling under a symmetry constraint, can very flexibly generate samples under arbitrary symmetry groups (Fig. 4a, Appendix L). Figure 4a illustrates symmetry-conditioned generation across many groups, from simple 4-subunit cyclic symmetries up to a capsid-sized icosahedral complex with 96,000 total residues and over 380,000 atoms. This also demonstrates why favorable computational scaling properties, such as quasilinear computation time (Appendix D), are important, as efficient computation facilitates scaling to larger systems. Symmetric assemblies are common in nature and there have been some successes with *de novo* symmetric designs [Wicky et al., 2022, King et al., 2014], but it has been generally challenging to simultaneously optimize for both the molecular interaction details between protomers and the desired overall symmetry in design. By enabling simple symmetry conditioning within the generation process, joint Chroma should make it simpler to more easily sample structures that simultaneously meet both requirements.

**Figure 4:**
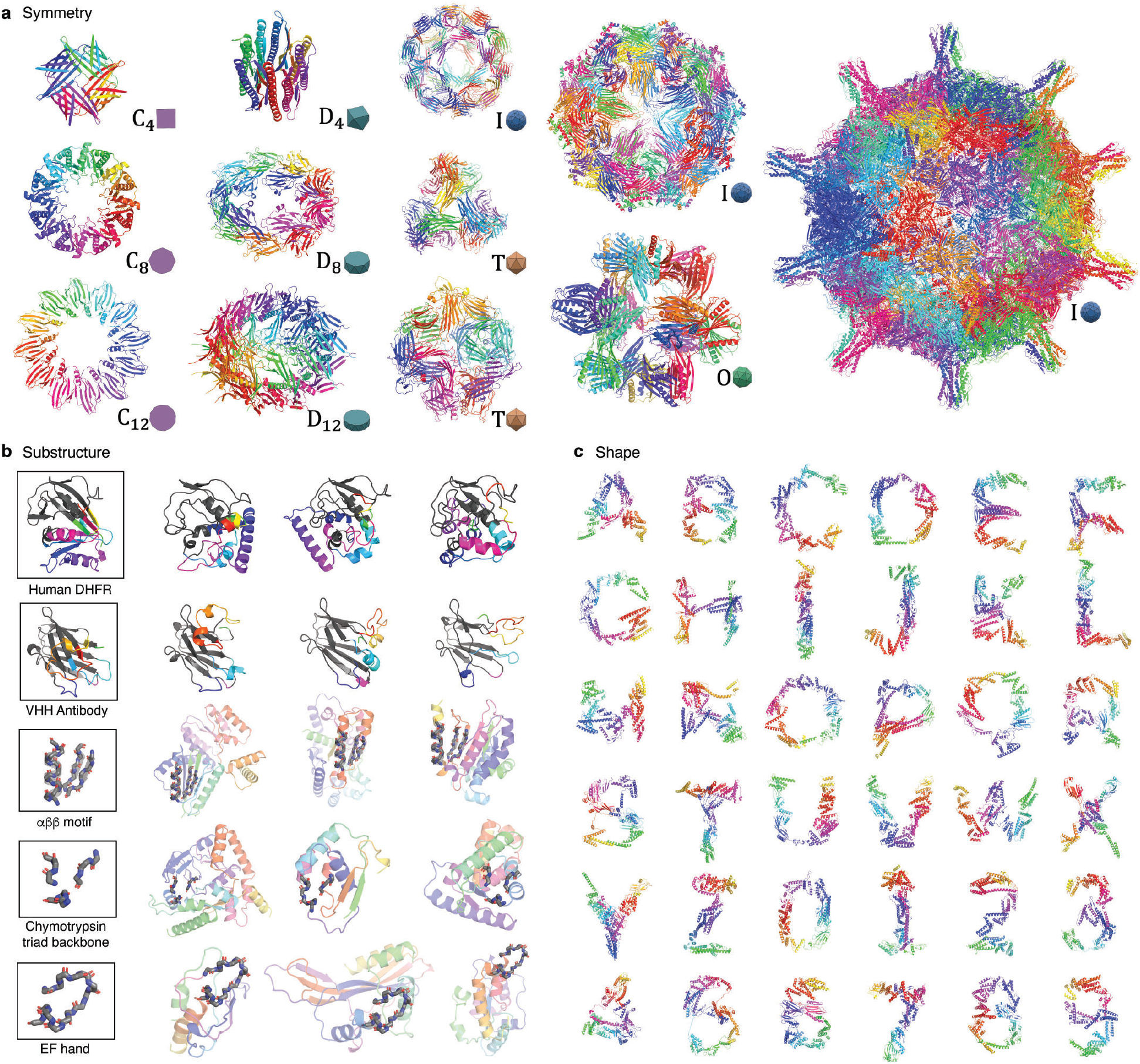
Symmetry, substructure, and shape conditioning enable geometric molecular programming. **a**, Conditioning on arbitrary symmetry groups is possible by symmetrizing gradient, noise, and initialization through the sampling process (Appendix L). We show how cyclic *C_n_*, dihedral *D_n_*, tetrahedral *T*, octahedral *O*, and icosahedral *I* symmetries can produce a wide variety of possible homomeric complexes. The righmost protein complex contains 60 subunits and 96,000 total residues. **b**, Conditioning on partial substructure (monochrome) enables protein “infilling” or “outfilling”. Top two rows illustrate regeneration (color) of half of a protein (enzyme DHFR, first row) or CDR loops of an antibody (second row); Appendix K. Next three rows show conditioning on a pre-defined motif; order and matching location of motif segments is not pre-specified here. **c**, Lastly, it is possible to condition on arbitrary volumetric shapes by using gradients derived from Optimal Transport (Appendix L). We test the ability of Chroma to solve for backbone configurations subject to the complex geometries of the Latin alphabet and numerals.

Next, we explore substructure conditioning in Fig. 4b, which is a central problem for protein design as it can facilitate preserving one part of a protein’s structure (and function) while modifying another part of the structure (and potentially function). In the top row, we “cut” the structure of human dihydrofolate reductase (PDB code 1DRF) into two halves with a plane, remove one of the halves, and regenerate the other half anew. The cut plane introduces several discontinuities in the chain simultaneously, and the generative process needs to sample a solution that satisfies these boundary conditions while being realistic. Nevertheless, the samples achieve both goals and, interestingly, do so in a manner very different from both each other and from natural DHFR. In the second row of Fig. 4b, we cut out the complementarity-determining regions of a VHH antibody and rebuild them conditioned on the remaining framework structure. The generated structures are once again plausible despite similar difficulties to the DHFR example. Lastly, in the bottom three rows of Fig. 4b, condition on sub-structure in a *unregistered* manner, meaning that the exact alignment of the substructure (motif) within the chain is not specified *a priori* as it was in the prior examples. We ”outfill” protein structure around several structural and functional motifs, including an *αββ* packing motif, backbone fragments encoding the catalytic triad active site of chymotrypsin, and the EF-hand Ca-binding motif. Again, these motifs are accommodated in a realistic manner using diverse and structured solutions.

In Fig. 4c we provide an early demonstration of a more exotic kind of conditioning in which we attempt to solve for backbone configurations subject to arbitrary volumetric shape specifications. We accomplish this by adding heuristic classifier gradients based on optimal transport distances [Peyré et al., 2019] between atoms in the structures and user-provided point clouds (Appendix M). As a stress test of this capability, we conditioned the generation of 1,000-residue single protein chains on the shapes of the Latin alphabet and Arabic numerals. While it remains unclear if any of these backbones would be sufficiently realistic to autonomously fold into their intended shapes, we see the model routinely implementing several core phenomena of protein backbones such as high secondary structure content, close packing with room for designed sidechains, and volume-spanning alpha-helical bundle and beta sheet elements. Although these shapes represent purely a challenging set of test geometries, more generally, shape is intimately related to functions in biology, for example, with membrane transporters, receptors, and structured assemblies that organize molecular events in space. Being able to control shape would be a useful subroutine for generalized protein engineering and design.

Finally, we demonstrate in Fig. 5 that it is possible to condition on protein *semantics* such as secondary structure, fold class (Fig. 5a) and natural language (Fig. 5b). Unlike for the geometric conditioning where the classifier is correct by construction (e.g., the presence of a motif under a certain RMSD is unambiguous), here the classifiers are neural networks trained on structure data (and structure data are considerably more sparse than image data), so there can be a discrepancy between the label assigned by the classifier and the ground truth class. Thus, looking at the fold-conditioned generation (Fig. 5a), we see that conditional samples always improve classifier probabilities over unconditioned samples taken from the same random seed, but the classification is not always perfect. For example, for the cases of “beta barrel” and “Ig fold” classes, the generated samples look like believable representatives of the respective class. On the other hand, in the “Rossman fold” example, the structure has some of the features characteristic of the class (i.e., two helices packed against a sheet on one side), but does not contain all such features (e.g., the opposing side of the sheet is not fully packed with helices like in a classical Rossman fold). In Fig. 5b we demonstrate semantic conditioning on natural language captions, which similarly improves probabilities while not generically being valid. It is exciting to imagine the potential of such a capability—i.e., being able to request desired protein features and properties directly via natural language prompts. Generative models such as Chroma can reduce the challenge of function-conditioned generation to the problem of building accurate classifiers for functions given structures. While there is clearly much more work to be done to make this useful in practice, high-throughput experiments and evolutionary data can likely make this possible in the near term.

**Figure 5:**
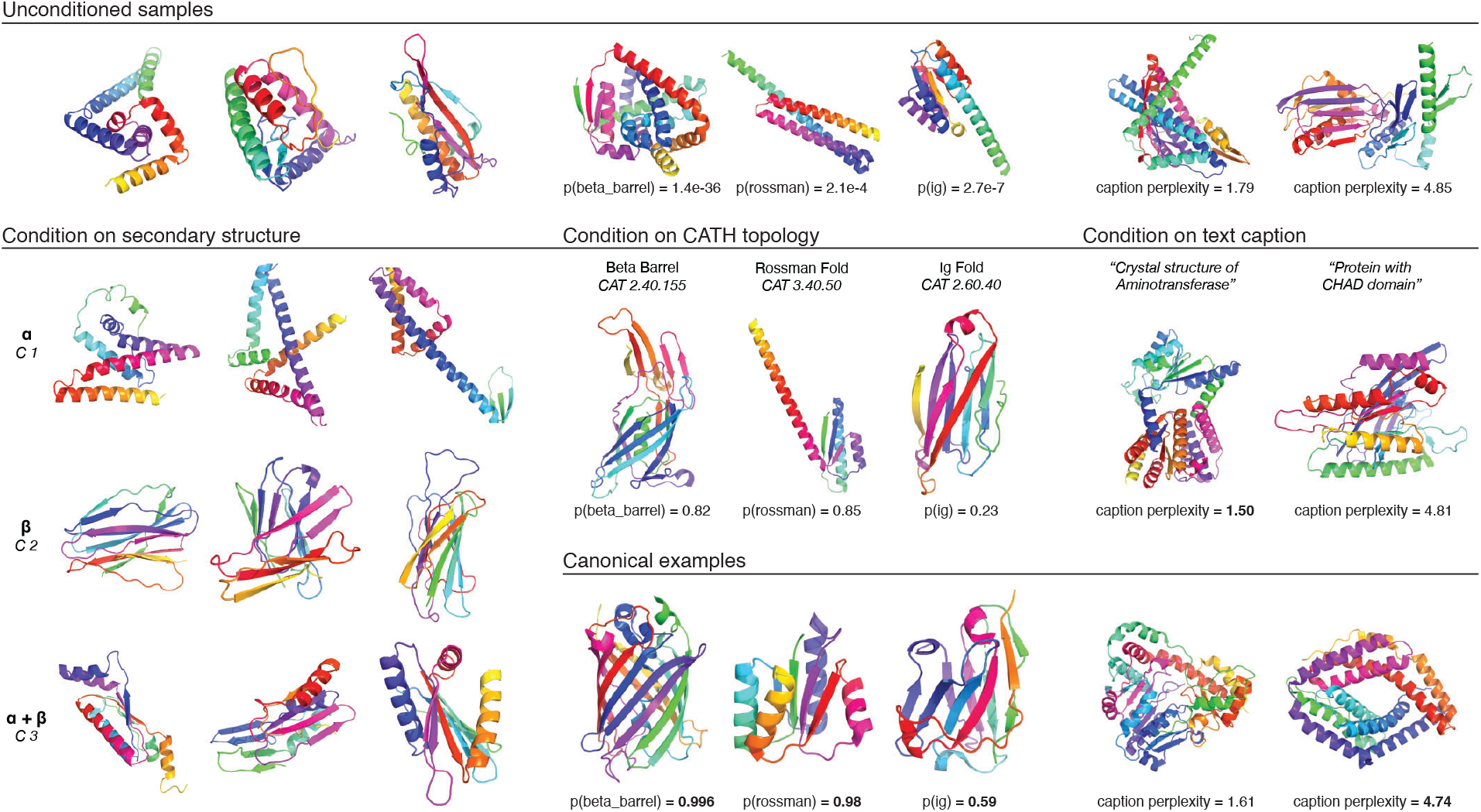
Protein structure classifiers and caption models can bias the sampling process towards user-specified properties. The top row shows example structures drawn unconditionally from the *p* (structure) model. Below, models trained to predict protein semantics are used to conditionally sample structures with desired secondary structures, belonging to particular topologies, or corresponding to natural language captions. In each column, all conditional samples are drawn starting from the same random seed as the unconditional sample shown at the top of the column. The samples based on secondary structure conditioning show the impact of classifiers trained to predict mainly alpha, mainly beta, and mixed alpha-beta structures. In the columns with topology-conditioned samples, the classifier’s predicted probabilities for the intended topology are indicated. Similarly, in the columns with samples based on text conditioning, the caption model’s average perplexities are shown. For the topology and text caption columns, PDB structures are shown (“Canonical examples”) that exemplify the target condition.

**Figure 6:**
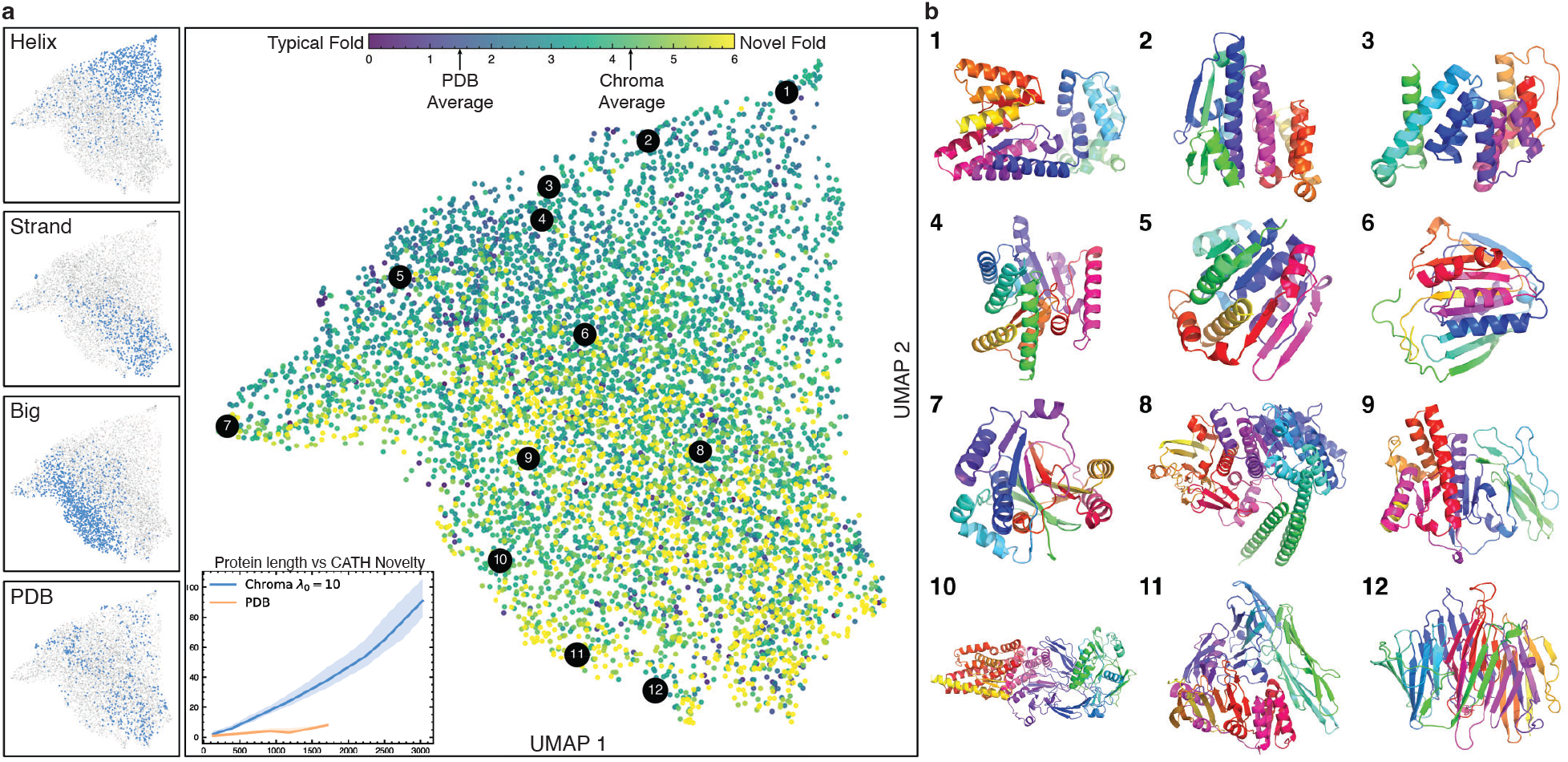
Chroma-generated structures span natural protein space while also frequently demonstrating high novelty. **a**, Proteins from the PDB and Chroma are featurized with 31 global fold descriptors derived from knot theory [Røgen and Fain, 2003, Harder et al., 2012] and are embedded into two dimensions using UMAP. The large figure is colored by the CATH coverage novelty measure normalized by protein length. Structural novelty was assessed by counting the number of CATH domains needed to achieve a greedy cover at least 80% of residues with TM > 0.5. On average Chroma needs 4.3 CATH domains per 200 amino acids to cover 80% of its residues while structures from the PDB need only 1.6. (**a**, inset) Chroma generated structures are more diverse and novel compared to structures from the PDB (regardless of protein length). The line represents the median value and is bounded by first (25%) and third (75%) quartile bands. The 4 smaller UMAP plots demonstrate the structure of the embedding by highlighting populations of structures that are mainly helices, strands, large (more than 500 residues), or natural proteins. The panel labeled *PBD* shows the distribution of natural proteins used to train the model. **b**, We render twelve proteins from across the embedding space with high novelty score (numbered in the embedding plot). The highlighted structures all have a novelty score of at least one standard deviation greater than the PDB.

### Expanding protein space

So far, we have shown how samples from Chroma are realistic, designable, and can be conditioned, but it remains unclear how much they simply reproduce structures from the training set or involve novel fold topologies. We sought to characterize the novelty of samples from Chroma by taking our large set of single-chain samples (Section “Analysis of unconditional samples”) and mapping their structural homology to proteins in the PDB. Classifying protein structures into distinct topologies is not simple, but initiatives such as the CATH database [Sillitoe et al., 2021] have identified a relatively compact set of structural domains, which can efficiently explain most of the structures in the PDB as sub-domain combinations. Therefore, we can invoke a notion of “Novelty as CATH-compressibility” where we ask how easy it is to reduce Chroma-sampled structures to the compositions of CATH domains. We define a novelty score as the number of CATH domains required to achieve a greedy cover of 80% of the residues in a protein at a TM score above 0.5. Note that most valid proteins will be covered by at least *some* finite number of CATH domains as we include even very small domains in the coverage test (including those comprising a single secondary-structural element). The results of this analysis are shown in the inset of Fig. 6a, where the novelty score is plotted as a function of protein length for both Chroma-generated and native proteins. The gap between the two is apparent. Most native backbones are described by a handful of CATH domains, with this number rising slowly as a function of length. On the other hand, Chroma-generated structures routinely require tens of CATH domains to reach 80% coverage, and this number rises sharply with length.

We further find that samples from Chroma are diverse and covering of all of natural protein space. In Fig. 6a, we jointly represent samples from Chroma and a set of structures from the PDB with global topology descriptors derived from knot theory [Røgen and Fain, 2003, Harder et al., 2012], and embed these into two dimensions with UMAP [McInnes et al., 2018]. The resulting embedding appears to be semantically meaningful as sub-sets of structures belonging to different categories by size and secondary structures appear to cluster in this projection (sub-panels on the left in Fig. 6a). Coloring individual points in the embedding by the degree of novelty (i.e., the length-normalized number of CATH domains required to achieve 80% coverage), we can see that novelty in the space being sampled is spread broadly and not biased to only certain types of structures (Fig. 6a, right). This is especially clear when looking at a representative selection of novel samples shown in Fig. 6b (novelty threshold for selecting being > 2.2 CATH domains needed for 80% per 200 amino acids of length). Taken together, these results show that Chroma samples are diverse and have not yet appeared to exhibit any obvious biases.

## Discussion

In this work, we present Chroma, a new generative model capable of generating novel and diverse proteins across a broad array of structures and properties. Chroma is *programmable* in the sense that it can be conditioned to sample proteins with a wide-array of user-specified properties, including: inter-residue distances and contacts, domains, sub-structures, and semantic specifications from classifiers. Chroma is able to generate proteins that have arbitrary and complex shapes, and we have shown the beginning of the ability to accept descriptions of desired properties as free text. Due to an efficient design with a new diffusion process, quasilinear scaling neural architecture, and low-temperature sampling method, Chroma is able to generate extremely large proteins and protein complexes (e.g. with ≥ 3000 residues) on a commodity GPU (e.g., an NVIDIA V100) in a few minutes.

These results are particularly striking in light of the historical difficulty of sampling realistic protein structures. The task of exploring the structure space in a way that can produce physically reasonable and designable conformations has been a long-standing challenge in protein design. In a few protein systems, it has been possible to parameterize the backbone conformation space mathematically–most notably the *α*-helical coiled coil [Grigoryan and DeGrado, 2011] and a few other cases with high symmetry [Woolfson et al., 2015]—and in these cases design efforts have benefited tremendously creating possibilities not available in other systems [Beesley and Woolfson, 2019, Woolfson et al., 2015]. For all other structure types, however, a great amount of computational time is being spent on the search for reasonable backbones, often leaving the focus on actual functional specifications out of reach. Chroma has the potential to address this problem, enabling a shift from focusing on generating feasible structures towards a focus on the specific task at hand— i.e., what the protein is intended to do. By leveraging proteins sampled over the first 3+ billion years of evolution on Earth and finding new ways to assemble stable protein matter, generative models such as Chroma are well poised to drive another expansion of biomolecular diversity for human health and bioengineering.

## Acknowledgements

We would like to thank William F. DeGrado and Generate employees Adam Root, Alan Leung, Alex Ramos, Brett Hannigan, Eugene Palovcak, Frank Poelwijk, James Lucas, James McFarland, Karl Barber, Kristen Hopson, Martin Jankowiak, Mike Nally, Molly Gibson, Ross Federman, Stephen DeCamp, Thomas Linsky, Yue Liu, and Zander Harteveld for reading of the manuscript draft and providing helpful comments.

## Supplementary Information

**Table 1:**
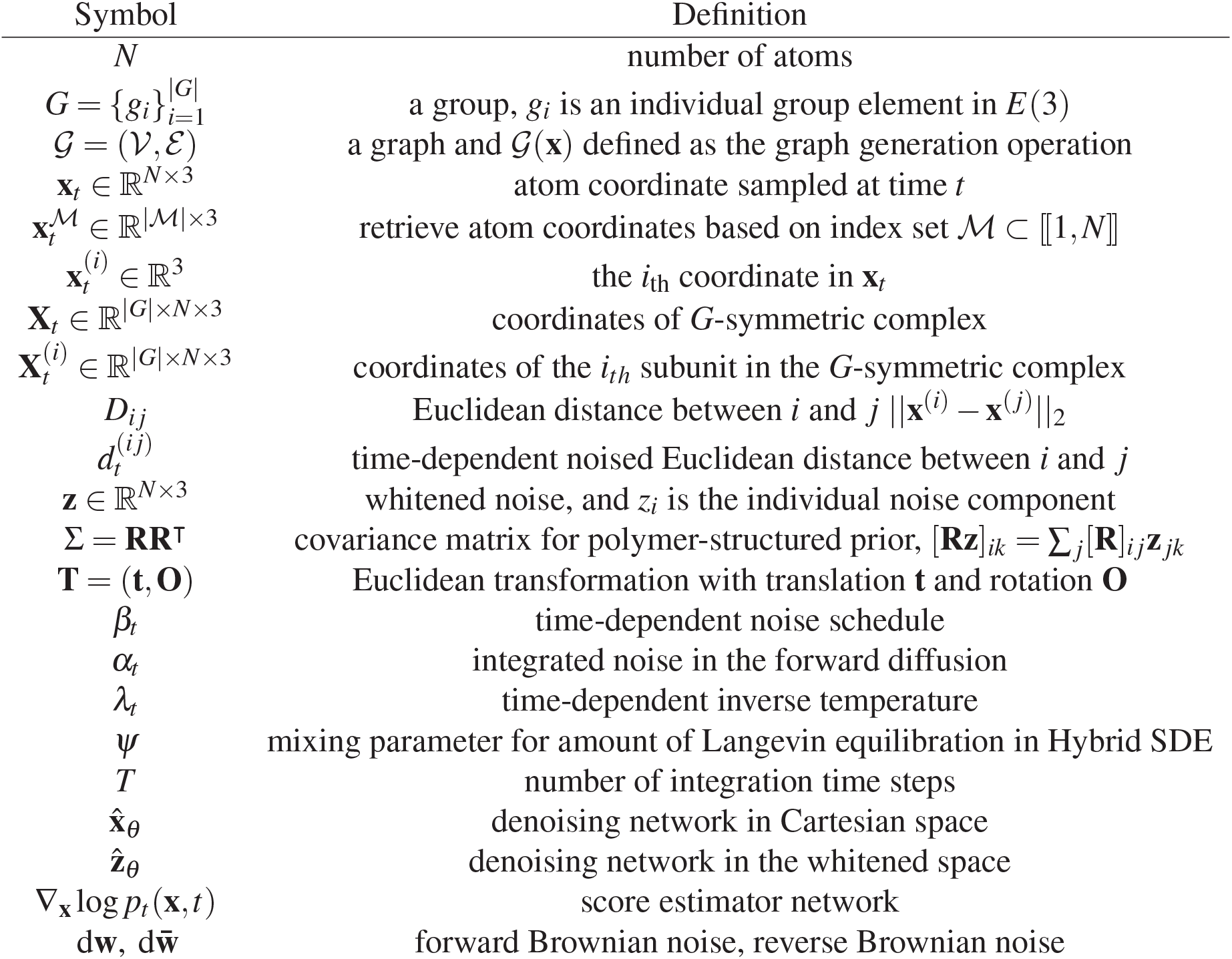
Table of notation.

| Symbol | Definition |
| --- | --- |
| $N$ | number of atoms |
| $G = \{g_i\}_{i=1}^{\vert G\vert }$ | a group, $g_i$ is an individual group element in $E(3)$ |
| $\mathcal{G} = (\mathcal{V}, \mathcal{E})$ | a graph and $\mathcal{G}(\mathbf{x})$ defined as the graph generation operation |
| $\mathbf{x}_t \in \mathbb{R}^{N \times 3}$ | atom coordinate sampled at time $t$ |
| $\mathbf{x}_t^{\mathcal{M}} \in \mathbb{R}^{\vert \mathcal{M}\vert \times 3}$ | retrieve atom coordinates based on index set $\mathcal{M} \subset \llbracket 1, N \rrbracket$ |
| $\mathbf{x}_t^{(i)} \in \mathbb{R}^3$ | the $i_{\text{th}}$ coordinate in $\mathbf{x}_t$ |
| $\mathbf{X}_t \in \mathbb{R}^{\vert G\vert \times N \times 3}$ | coordinates of $G$-symmetric complex |
| $\mathbf{X}_t^{(i)} \in \mathbb{R}^{\vert G\vert \times N \times 3}$ | coordinates of the $i_{th}$ subunit in the $G$-symmetric complex |
| $D_{ij}$ | Euclidean distance between $i$ and $j$ $\ \mathbf{x}^{(i)} - \mathbf{x}^{(j)}\ _2$ |
| $d_t^{(ij)}$ | time-dependent noised Euclidean distance between $i$ and $j$ |
| $\mathbf{z} \in \mathbb{R}^{N \times 3}$ | whitened noise, and $z_i$ is the individual noise component |
| $\Sigma = \mathbf{R}\mathbf{R}^\top$ | covariance matrix for polymer-structured prior, $[\mathbf{R}\mathbf{z}]_{ik} = \sum_j [\mathbf{R}]_{ij} \mathbf{z}_{jk}$ |
| $\mathbf{T} = (\mathbf{t}, \mathbf{O})$ | Euclidean transformation with translation $\mathbf{t}$ and rotation $\mathbf{O}$ |
| $\beta_t$ | time-dependent noise schedule |
| $\alpha_t$ | integrated noise in the forward diffusion |
| $\lambda_t$ | time-dependent inverse temperature |
| $\psi$ | mixing parameter for amount of Langevin equilibration in Hybrid SDE |
| $T$ | number of integration time steps |
| $\hat{\mathbf{x}}_\theta$ | denoising network in Cartesian space |
| $\hat{\mathbf{z}}_\theta$ | denoising network in the whitened space |
| $\nabla_{\mathbf{x}} \log p_t(\mathbf{x}, t)$ | score estimator network |
| $d\mathbf{w}, d\bar{\mathbf{w}}$ | forward Brownian noise, reverse Brownian noise |

## A Diffusion Models with Structured Correlations

### A.1 Correlated diffusion as uncorrelated diffusion in whitened space

#### Correlation and diffusion

Most natural data possess a hierarchy of correlation structures, some of which are very simple (e.g., most nearby pixels in natural images will tend to be a similar color) and some of which are very subtle (e.g., complex constraints govern the set of pixels forming an eye or a cat). With finite computing resources and modeling power, it can be advantageous to design learning systems that capture simple correlations as efficiently as possible such that most model capacity can be dedicated to nontrivial dependencies (see Appendix C).

Diffusion models capture complex constraints in the data by learning to reverse a diffusion process that transforms data into noise [Sohl-Dickstein et al., 2015, Song et al., 2021]. While most of these original diffusion frameworks considered the possibility of correlated noise, it is typical in contemporary models to use isotropic noise that is standard normally distributed. In this configuration, models must learn both simple correlations and complex correlations in data *from scratch*.

#### Whitening transformations and linear generative models

One classical approach for removing nuisance correlations in data is to apply a “whitening transformation”, i.e., an affine linear transformation 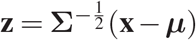 that decorrelates all factors of variation by subtracting the empirical mean ***μ*** and multiplying by a square root of the inverse covariance matrix 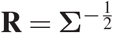.

Whitening data can also be related to fitting the data to a Gaussian model **x** = *F*(**z**) = **Rz** + **b** where the whitened factors **z** are standard normally distributed as 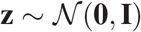 [Murphy, 2012]. The density in the whitened space can be related to the density in the transformed space by the change of variables formula as

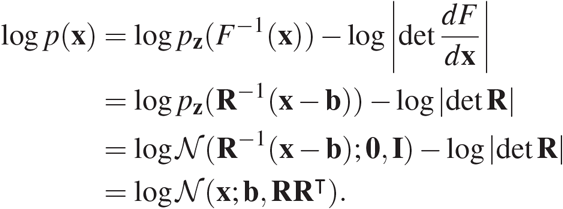

#### From whitened diffusion to dewhitened diffusion

If we possess a linear Gaussian prior for our data 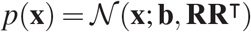 which can be sampled as **x** = **Rz** with 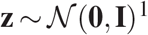^1^, then an *uncorrelated* diffusion process on the whitened coordinates **z**_*t*_ ~ *p_t_*(**z**|**z_0_**) will induce a *correlated* diffusion process on the original coordinates **x**_*t*_ ~ *p_t_*(**x**|**x_0_**). When the diffusion process is the so-called Variance-Preserving (VP) diffusion [Sohl-Dickstein et al., 2015, Ho et al., 2020], then the diffusion will transition from the data distribution at time *t* = 0 to the Gaussian prior distribution at time *t* = 1. Throughout this work we use the continuous time formulation of VP diffusion [Song et al., 2021] in whitened space. This process evolves in time *t* ∈ (0,1) according to the Stochastic Differential Equation (SDE)

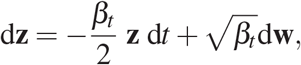

where **w** is a standard Wiener process and *β_t_* is the time-dependent schedule at which noise is injected into the process. We can also write the correlated SDE in terms of **x**_*t*_ if we substitute **x** = **Rz** as

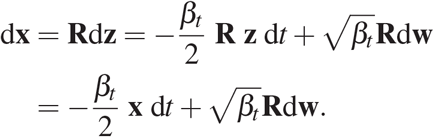

#### Sampling from the diffusion

This diffusion process is simple to integrate forward in time [Sohl-Dickstein et al., 2015, Song et al., 2021]. Given an initial data point **x**_0_, then **x**_*t*_ will be distributed as 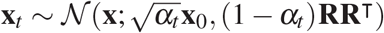 where 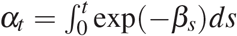 is the integrated noise. Samples at any time *t* can thus be generated from standard normally distributed noise as

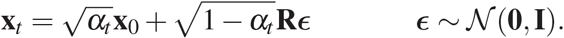

### A.2 Evidence Lower Bound (ELBO)

#### Denoising loss

Diffusion models can be parameterized in terms of a denoising neural network 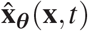 that is trained to predict **x_0_** given a noisy sample **x**_*t*_. Typically this is done by minimizing a *denoising loss*

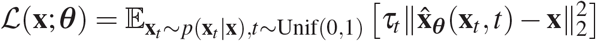

where *τ_t_* is a time-dependent weighting to emphasize the loss at particular points in time (noise levels) [Song et al., 2021]. Training with this loss can be directly related to *score matching* and *noise prediction* which can be cast as alternative parameterizations of the target output of the network [Kingma et al., 2021].

#### Approximate likelihood bound

We train the diffusion model by optimizing an approximate bound on the marginal likelihood of data together with a regularization loss. Following [Kingma et al., 2021], the negative Evidence Lower Bound (ELBO) in whitened space is

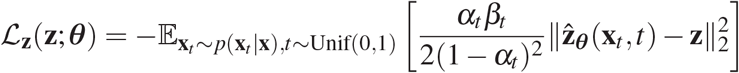

where in this case 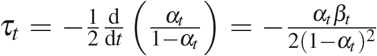 is the derivative of the *signal to noise ratio* 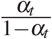. To express this loss in terms of **x** = **Rz** we may again apply change of variables formula to obtain

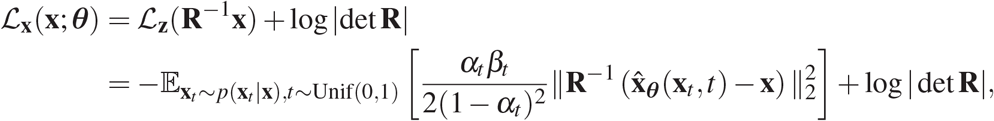

where the term log | det **R**| is a constant (we fit the parameters of **R** offline from training) and can be ignored during optimization of the denoiser parameters ***θ***.

#### Regularized loss

In practice we optimize a regularized variant of the ELBO loss which is the sum of whitened and unwhitened reconstruction errors as

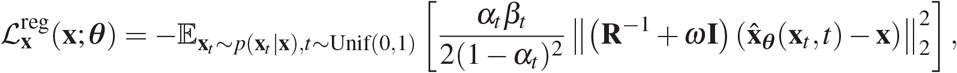

where **I** is the identity matrix and we set the scale factor *ω* to give **x** units of nanometers. We found this regularization to be important because in practice we care about absolute errors in **x** space, i.e. absolute spatial errors, at least as much as we care about errors in **z** space, which will correspond under our covariance models (Appendix C) to relative local chain geometries.

### A.3 Reverse-time SDE

In whitened space, we can express the *reverse-time* dynamics for the forwards-time SDE in terms of another SDE [Anderson, 1982, Song et al., 2021] that depends on the *score function* of the time-dependent marginals ∇_z_ log *p_t_* (**z**) as

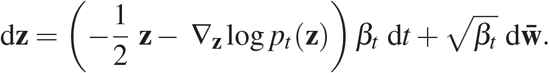

We can similarly express this in the score function of the transformed coordinate system as

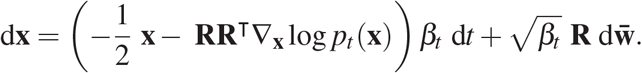

To sample from the diffusion model by taking a sample from the “prior” (time 1 distribution) and integrate the SDE above backward in time from *t* = 1 to *t* = 0. We can rewrite the above SDE in terms of our optimal denoising network 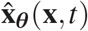 (trained as described above) by leveraging the relationship [Song et al., 2021, Kingma et al., 2021] that

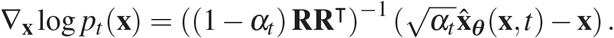

Therefore we can express the reverse-time SDE in terms of the optimal denoising network 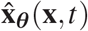 as

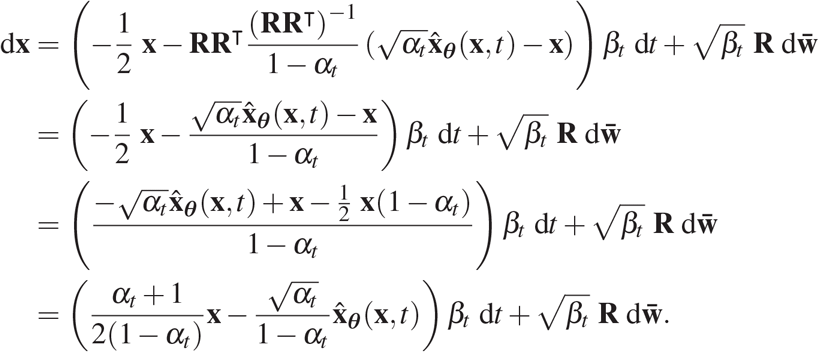

### A.4 Probability Flow ODE

#### Probability Flow ODE for deterministic encoding and sampling

Remarkably, it is also possible to derive a set of deterministic *ordinary differential equations* (ODEs) whose marginal evolution from the prior is identical to above SDEs [Song et al., 2021, Maoutsa et al., 2020]. In the context of our covariance model this can be expressed either in terms of the score function ∇_**x**_ log *p_t_* (**x**) as

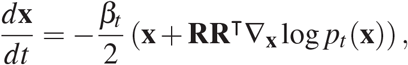

or in terms of the optimal denoiser network 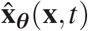 as

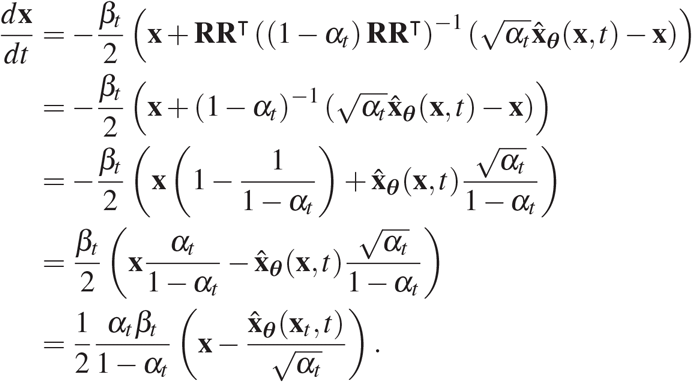

The ODE formulation of sampling is especially important because it enables reformulating the model as a Continuous Normalizing Flow [Chen et al., 2018], which can admit efficient and exact likelihood calculations using the adjoint method [Grathwohl et al., 2018].

### A.5 Conditional sampling from the posterior under auxiliary constraints

#### Bayesian posterior SDE for conditional sampling

An extremely powerful aspect of the reverse diffusion formulation is that it can also be extended to enable conditional sampling from a Bayesian posterior *p*(**x**|**y**) by combining with auxilliary classifiers log *p_t_*(**y**|**x**) and without re-training the base diffusion model [Song et al., 2021]. When extended to the correlated diffusion case, this gives the SDE

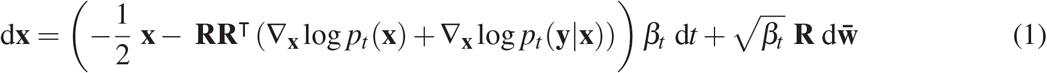

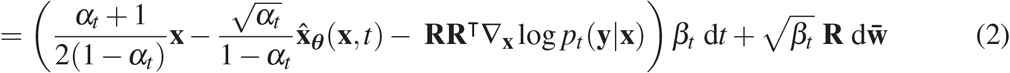

#### Bayesian posterior ODE for conditional sampling

In the context of our covariance model and conditional constraints, the Probability Flow ODE for sampling from the posterior is

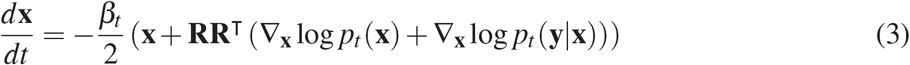

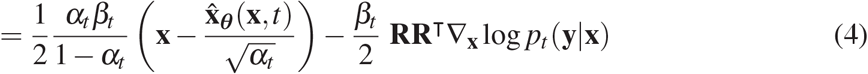

### A.6 Related work

Subspace diffusion models [Jing et al., 2022] also consider correlated diffusion, with a particular emphasis on focusing the diffusion to most relevant factors of variation for statistical and computational efficiency. Additionally, latent-space diffusion models [Rombach et al., 2022] might be viewed as learning a transformed coordinate system in which the diffusion process can more efficiently model the targer distribution. Our work provides further evidence for how correlated diffusion may be an underutilized approach to distributional modeling and shows how domain knowledge can be incorporated in the form of simple constraints on the covariance structure of the noise process.

## B Low-Temperature Sampling for Diffusion Models

Maximum likelihood training of generative models enforces a tolerable probability of *all* data-points and, as a result, misspecified or low-capacity models fit by maximum likelihood will typically be overdispersed. This can be understood through the perspective that maximizing likelihood is equivalent to minimizing the KL divergence from the model to the data distribution, which is the mean-seeking and mode-covering direction of KL divergence.

To mitigate overdispersion in generative models, it is common practice to introduce modified sampling procedures that increase sampling of high-likelihood states (mode emphasis, precision) at the expense of reduced sample diversity (mode coverage, recall). This includes approaches such as shrunken encodings in normalizing flows [Kingma and Dhariwal, 2018], low-temperature greedy decoding algorithms for language models [Holtzman et al., 2020], and stochastic beam search [Kool et al., 2019].

A powerful but often intractable way to trade diversity for quality in generative models is low-temperature sampling. This involves perturbing a base distribution *p*(**x**) by exponentiating with an inverse temperature rescaling factor *λ* and renormalizing as 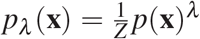. As the inverse temperature becomes large *λ* ≪ 1, this perturbed distribution will trade diversity (entropy) for sample quality (likelihood) and ultimately will collapse into the global optimum as *λ* → ∞. Unfortunately, low temperature sampling in the general case will require expensive iterative sampling methods such as Markov Chain Monte Carlo which typically offer no guarantee of convergence in a practical amount of time [MacKay, 2003].

### Low temperature and diffusion models

The issue of trading diversity for sample quality in diffusion models has been discussed previously, with some authors reporting that simple modifications like upscaling the score function and/or downscaling the noise were ineffective [Dhariwal and Nichol, 2021]. Instead, classifier guidance and classifier-free guidance have been widely adopted as critical components of contemporary text-to-image diffusion models such as Imagen and DALL-E 2 [Ho and Salimans, 2021, Saharia et al., 2022, Ramesh et al., 2022].

### Equilibrium versus Non-Equilibrium Sampling

Here we offer an explanation for why these previous attempts at low temperature sampling did not work and produce a novel algorithm for low-temperature sampling from diffusion models. We make two key observations, explained in the next two sections

1. **Upscaling the score function of the reverse SDE is insufficient** to properly re-weight populations in a temperature perturbed distribution.
2. **Annealed Langevin dynamics *can* sample from low temperature distributions** if given sufficient equilibration time.

### B.1 Reverse-time SDE with temperature annealing

#### The isotropic Gaussian case

To determine how the Reverse SDE can be modified to enable (approximate) low temperature sampling, it is helpful to first consider a case that can be treated exactly: transforming a Gaussian data distribution 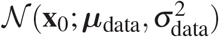 to a Gaussian prior 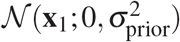. Under the Variance-Preserving diffusion, the time-dependent marginal density will be given by

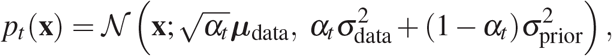

which means that the score function **s**_*t*_ will be

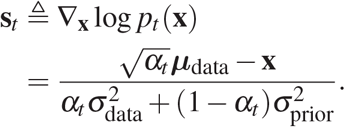

Now, suppose we wish to *modify* the definition of the time-dependent score function so that, instead of transitioning to the original data distribution, it transforms to the perturbed data distribution, i.e. so the it transitions to 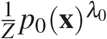. For a Gaussian, this operation will simply multiply the precision (or equivalently, divide the covariance) by the factor *λ*_0_. The perturbed score function will therefore be

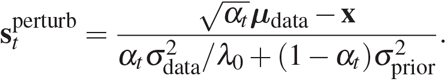

Based on this, we can express the perturbed score function as a *time-dependent rescaling* of the original score function with scaling based on the ratios of the time-dependent inverse variances as

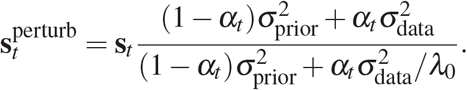

**Supplementary Figure 1:**
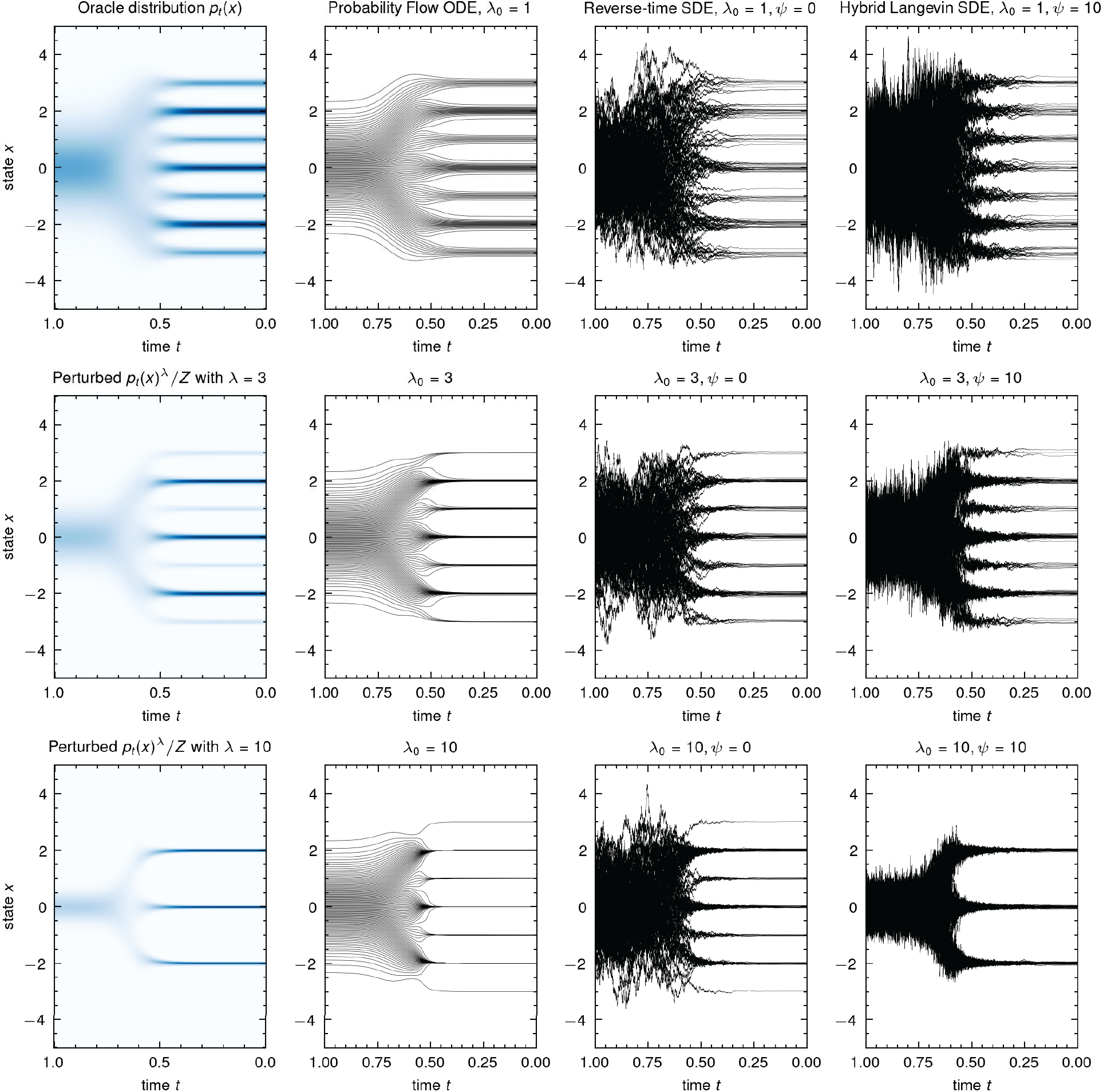
The Hybrid Langevin SDE can sample from temperature-perturbed distributions. The marginal densities of the diffusion process *p_t_* (*x*) (top left) gradually transform between a toy 1D data distribution at time *t* = 0 and a standard normal distribution at time *t* = 1. Reweighting the distribution by inverse temperature *λ*_0_ as 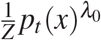 (left column, bottom two rows) will both concentrate and reweight the population distributions. The annealed versions of the reverse-time SDE and Probability Flow ODEs (middle columns) can concentrate towards local optima but do not correctly reweight the relative population occupancies. Adding in Langevin dynamics with the Hybrid Langevin xSDE (right column) increases the rate of equilibration to the time-dependent marginals and, when combined with low temperature rescaling, successfully reweights the populations (bottom right).

Therefore we see that, to achieve a particular inverse temperature *λ*_0_ for the data distribution, we should rescale the learned score function by time-dependent factor

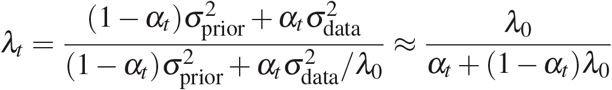

where in the last step we assumed 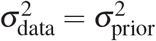. So one interpretation of the previously observed insufficienes of low temperature sampling based on score-rescaling [Dhariwal and Nichol, 2021] is that these were hampered by *uniform* rescaling the score function in time instead of in a way that accounts for the shift of influence between the prior and the data distribution.

#### Temperature-adjusted reverse time SDE

We can modify the Reverse-time SDE by simply rescaling the score function with the above time-dependent temperature rescaling as

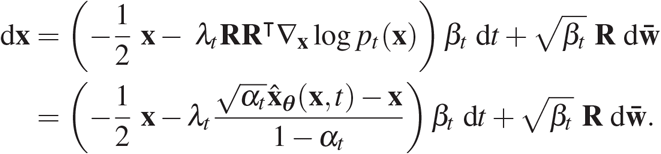

#### Temperature adjusted probability flow ODE

Similarly for the Probability Flow ODE we can rescale as

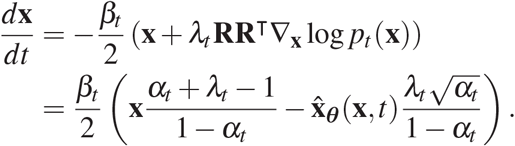

#### Rescaling does not reweight

We derived the above rescaling rationale by considering a uni-modal Gaussian, which has the simple property that the score of the perturbed diffusion can be expressed as a rescaling of the learned diffusion. This will not necessarily be true in general, and sure enough we find that the above dynamics *do drive towards local maxima* but *do not reweight populations based on their relative probability* (Supplementary Figure 1) as true low temperature sampling does. To address this, we next introduce an equilibration process that can be arbitrarily mixed in with the non-equilibrium reverse dynamics.

### B.2 Annealed Langevin Dynamics SDE

Instead of reversing the forwards time diffusion in a non-equilibrium manner, we can also use the learned time-dependent score function ∇_x_ log*p_t_* (**x**) (expressed in terms of the optimal denoiser 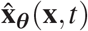) to do slow, approximately equilibrated sampling with annealed Langevin dynamics [Song and Ermon, 2019].

**Supplementary Figure 2:**
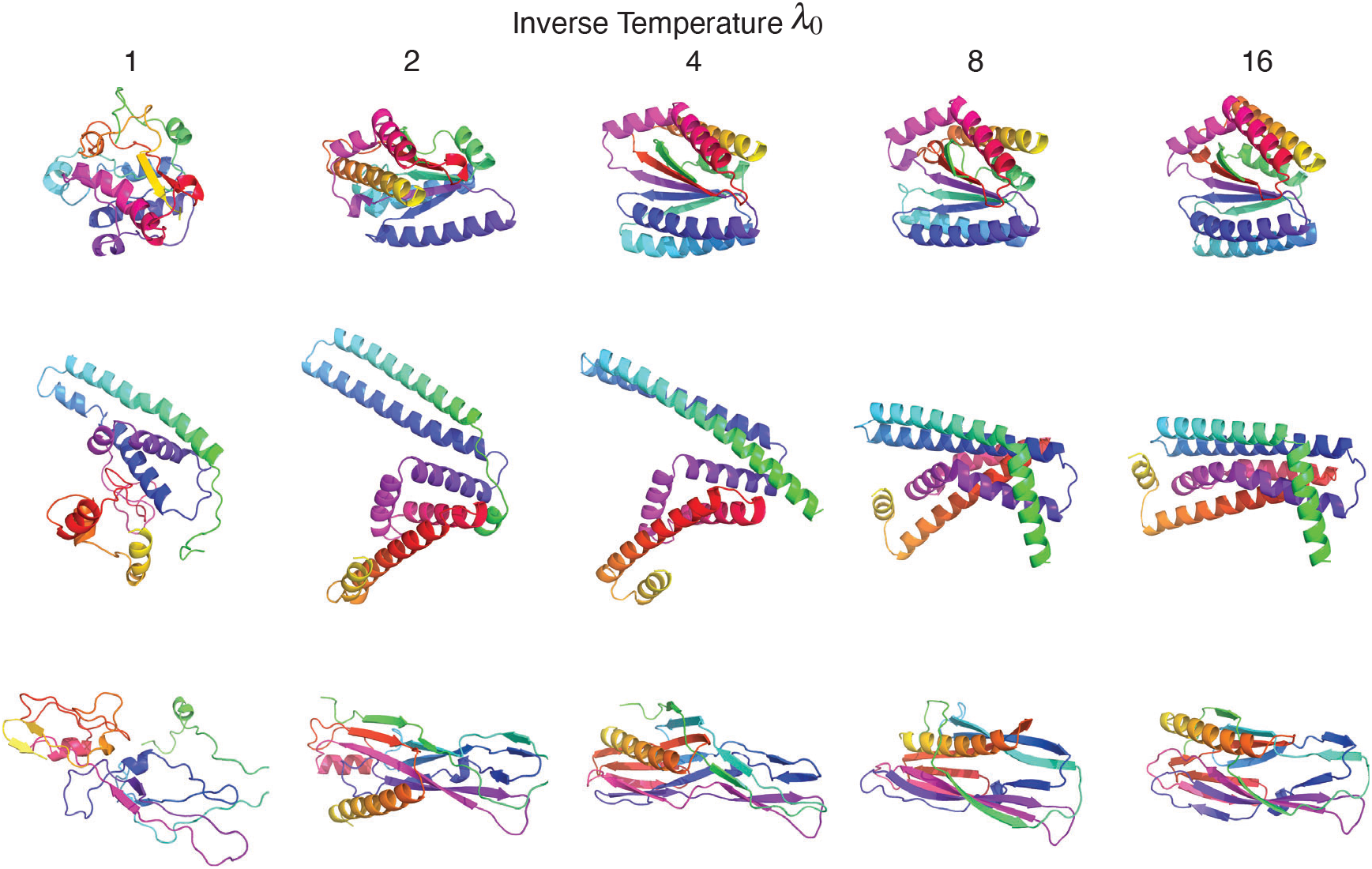
Low-temperature sampling trades reduced sample diversity (entropy) for increased sample quality (likelihood). Given a fixed random seed for generation (each row), structures sampled at higher inverse temperature *λ*_0_ leads have higher secondary structure content and tighter packing as compact, globular folds.

While the annealed Langevin dynamics of [Song and Ermon, 2019] was originally framed via discrete iteration, we can recast it in continuous time with the SDE

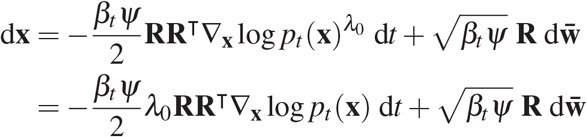

where *ψ* is an “equilibration rate” scaling the amount of Langevin dynamics per unit time. As *ψ* → ∞ the system will instantaneously equilibrate in time, constantly adjusting to the changing score function. In practice, we can think about how to set these parameters by considering a single Euler-Maruyama integration step in reverse time with step size 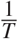 where *T* is the total number of steps

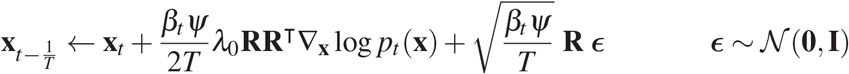

which is precisely preconditioned Langevin dynamics with step size 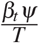. For a sufficiently small interval (*t* – d*t*, *t*) we can keep the target density approximately fixed while increasing *T* to do an arbitrarily large number of Langevin dynamics steps, which will asymptotically equilibrate to the current density log *p_t_* (**x**).

### B.3 Hybrid Langevin-Reverse Time SDE

We can combine the annealed Reverse-Time SDE and the Langevin Dynamics SDE into a hybrid SDE that infinitesimally combines both dynamics. Denoting the inverse temperature as *λ*_0_ and the ratio of the Langevin dynamics to conventional dynamics as *ψ*, we have

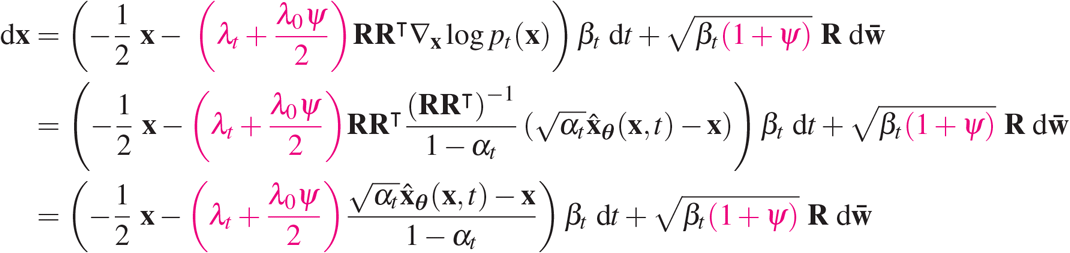

where we highlight in pink the terms that, when set to unity, recover the standard reverse time SDE.

Representative samples using this modified SDE are shown in Fig 7. Without the low temperature modification, this idea is very reminiscent of the Predictor Corrector sampler proposed by Song et al. [2021], except in that case the authors explicitly alternated between reverse-time diffusion and Langevin dynamics while we fuse them into a single SDE.

#### Equilibration is not free

Generally speaking, as we increase the amount of Langevin equilibration with *ψ*, we will need to simultaneously increase the resolution of our SDE solution to maintain the same level of accuracy. That said, we found that even a modest amount of equilibration was sufficient to significantly improve sample quality in practice with *ψ* ∈ [1,5].

#### Even more equilibration

Lastly, while the Hybrid Langevin-Reverse Time SDE can do an arbitrarily large amount of Langevin dynamics per time interval which would equilibrate asymptotically in principle, these dynamics will still inefficiently mix between basins of attraction in the energy landscape when 0 < *t* ≪ 1. We suspect that ideas from variable-temperature sampling methods, such as simulated tempering [Marinari and Parisi, 1992] or parallel tempering [Hansmann, 1997], would be useful in this context and would amount to deriving an augmented SDE system with auxiliary variables for the temperature and/or copies of the system at different time points in the diffusion.

## C Polymer-Structured Diffusions

Most prior applications of diffusion models to images and molecules have leveraged uncorrelated diffusion in which data are gradually transformed by isotropic Gaussian noise. We found this approach to be non-ideal for protein structure applications for two reasons. First, noised samples break simple chain and density (e.g., radius of gyration, *R_g_*) constraints that almost all structures satisfy [Hong and Lei, 2009, Tanner, 2016], forcing the model to allocate a significant amount of capacity towards re-implementing these basic constraints. And second, this “out-of-distribution” aspect of high-noise samples tends to limit the performance of efficient domain-specific neural architectures for molecular systems. To this end, we introduce multivariate Gaussian distributions for protein structures that (i) are *SO*(3) invariant, (ii) enforce protein chain and radius of gyration statistics, and (iii) can be computed in linear time. Throughout this section, we will introduce covariance models for protein polymers (which can be thought of as a de-whitening transform **R**, see Appendix A) with parameters that can be fit *offline* from training the diffusion model.

### C.1 Preliminaries: Diffusion models linearly interpolate between the average quadratic forms of the data and prior

Here we show how the diffusion processes described above will predictably affect molecular geometry as a function of the covariance structure of the noising process. We will use this result to reflect on how the covariance structure should be designed. Squared distance 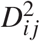 and the squared radius of gyration 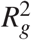 are both functions that can be expressed as quadratic forms in the coordinates. That means they can be expressed as a function 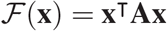 where **A** is a matrix weighting the different cross-terms as 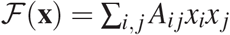. Suppose we want to understand the behavior of these quantities as they evolve under the forward process of a diffusion model. Recall that we can write samples from the forward diffusion process as

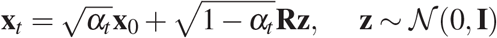

So we can write the time-expectation of any quadratic form as

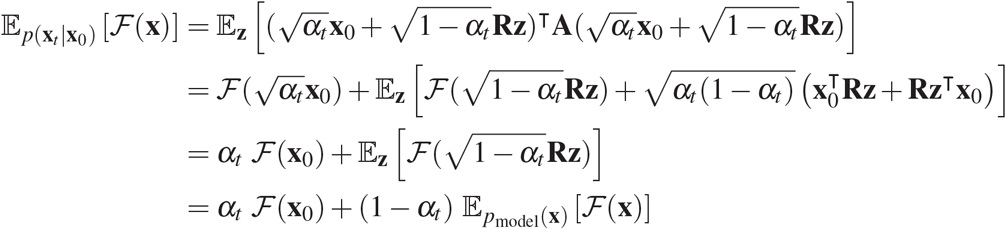

Squared distance is a quadratic form, so diffusion processes will simply linearly interpolate to the behavior of the prior as

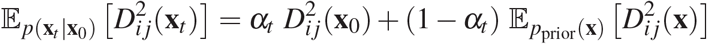

and squared radius of gyration will similarly evolve under the diffusion as

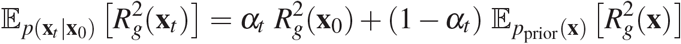

#### Punchline

Because diffusion models will do simple linear interpolations between the average squared distances and *R_g_* of the data distribution and of the prior, we should focus on covariance structures that empirically match these properties as closely as possible. Two primary ways will be in the chain constraint, i.e., that *D*_*i,i*+1_ (**x**_*t*_) should always be small and match the data distribution, and the size/density constraint of how 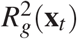 should behave as a function of protein length and typical packing statistics.

### C.2 Covariance model #1: Ideal chain

In this section, we introduce one of the simplest covariance models that enforces the chain constraint but ignores the *R_g_* scaling. It will interpolate between the data distribution and the ensemble of unfolded random coils.

#### Noise process

We index our amount of noise with a diffusion time *t* ∈ [0,1]. Given a denoised structure **x_0_**, a level of noise *t*, and a noise schedule *α_t_*, we sample perturbed structures from a Multivariate Gaussian distribution 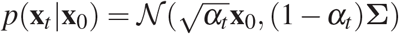 as

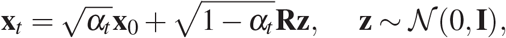

where the covariance matrix enforcing our chain constraint **Σ** = **RR**^⊤^ can be expressed in terms of its square root **R**, which is defined below.

Key to our framework is a matrix **R** whose various products, inverse-products, and transpose-products with vectors can be computed in *linear* time. We define the matrix **R** in terms of its product with a vector *f* (**z**) = **Rz**:

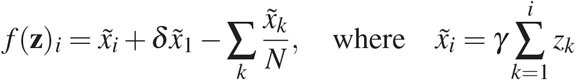

The inverse product *f*^-1^ (**x**) = **R**^-1^**x** is then

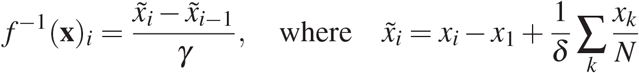

This definition of **R** induces the following inverse covariance matrix on the noise, which possesses a special structure:

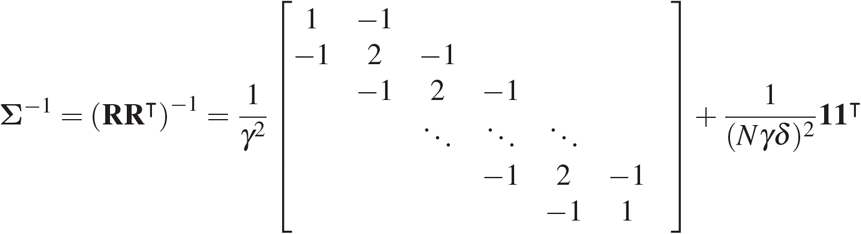

The parameter *γ* sets the length scale of the chain and the parameter *δ* sets the allowed amount of translational noise about the origin. This latter parameter is important for training on complexes where each chain may not have a center of mass at 0.

#### C.2.1 Covariance model #1 has ideal chain scaling *R_g_* ∝ *N*^(1/2)^

Our ideal-chain model is a simple Brownian motion and so the interatomic residual is Gaussian distributed with zero mean and *γ*^2^|*i* – *j*| variance, i.e.,

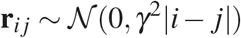

The expected squared norm for a Multivariate Normal Distribution (MVN) with spherical covariance is 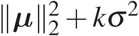 where *k* is the dimensionality, so we have:

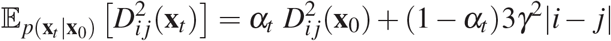

When *α_t_* = 0, the expected squared distances are those of the data distribution, while when *α_t_* = 1, they are those on an ideal Gaussian chain.

To compute the expected radius of Gyration, we can use the identity that it is simply half of the root mean square of inter-residue distances

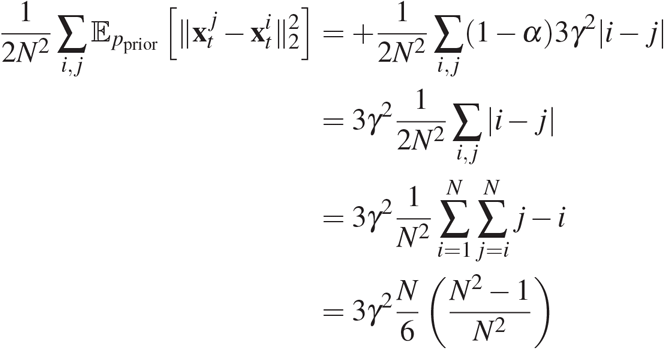

Therefore, we can also view the mean behavior of the diffusion as linearly interpolating the squared radius of gyration as

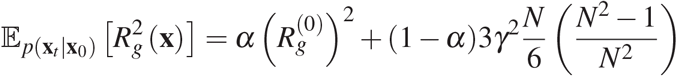

When *α* → 0 and *N* ≪ 0, the term 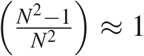 we recover the well-known scaling for an ideal chain with 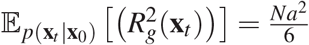 where the segment length is 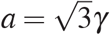.

#### C.3 Covariance model #2: *R_g_*-confined, linear-time Polymer MVNs

In this section we consider how to extend the previous model that also constrains the the scaling of the radius of gyration *R_g_*. We consider a family of two-parameter linear chain models that include the previous model as a special case. Specifically, consider the following linear recurrence

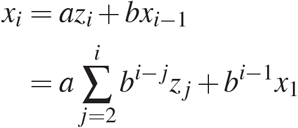

Here, the parameter *a* is a global scale parameter setting the “segment length” of the polymer and *b* is a “decay” parameter which sets the memory of the chain to fluctuations. We recover a spherical Gaussian when *b* = 0 and the ideal Gaussian chain when *b* = 1.

This system can also be written in matrix form as **x** = **Rz** with

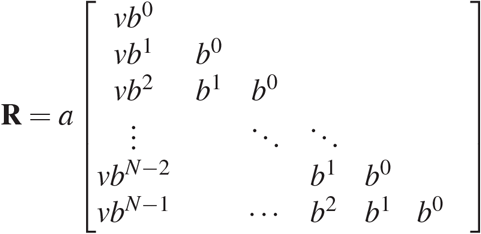

where 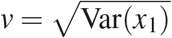.

We can solve for the equilibrium value of *v* via the condition Var(*x*_1_) = *a*^2^*v*^2^ = Var(*x_i_*) = Var(*x*_*i*-1_). The solution is

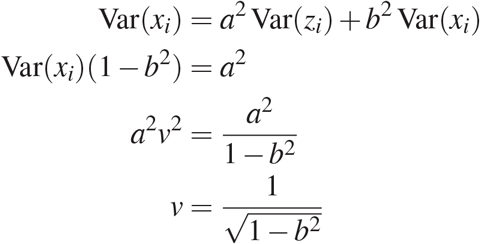

So our final recurrence is

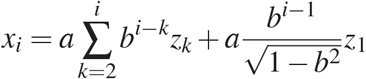

##### C.3.1 Expected 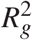 as a function of *b*

To compute the expected Radius of Gyration, we will use the identity 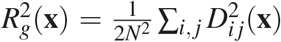, which we can compute via the variance of the residual between *x_i_* and *x_j_*. Assuming *j* > *i*, we have

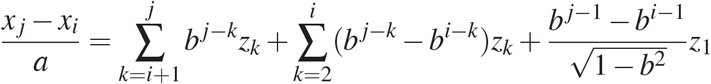

and the variance of which is

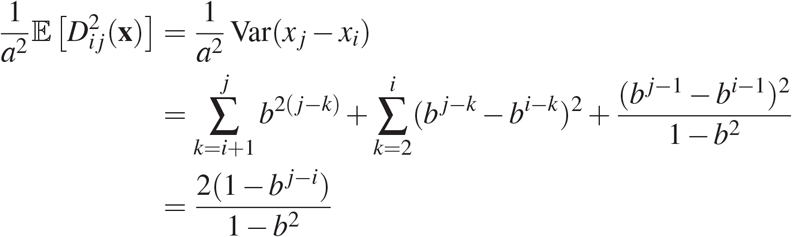

So the expected squared radius of gyration is

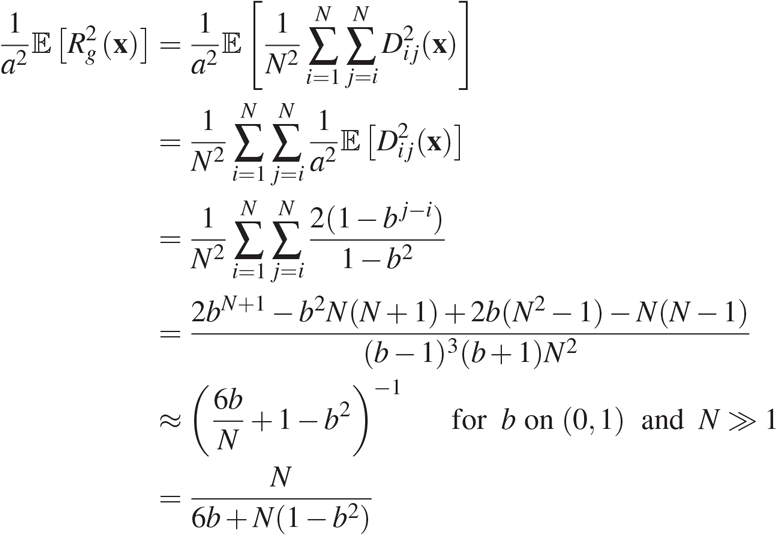

The approximation in the penultimate step works quite well in practice and becomes more accurate with growing *N*:

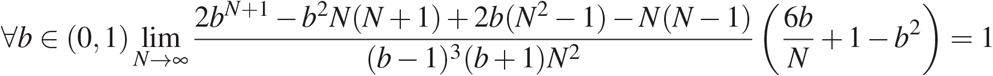

Additionally, we can verify that this result recovers the expected limiting behavior of an ideal unfolded chain when *b →* 1

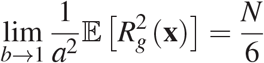

and of a standard normal distribution when *b* → 0

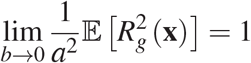

To finish up, we can add back in our global scaling factor *a* for our final result:

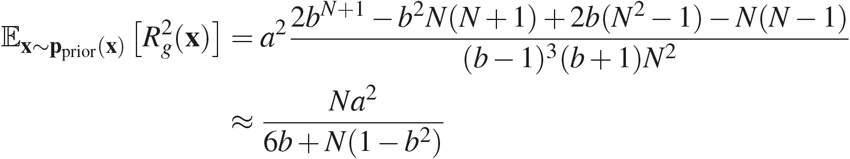

##### C.3.2 How to implement any 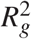 scaling

Equipped with a simple dependence of 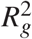 on *b*, we can solve for the correct spring strength by simply asking for what value of *b* we achieve our scaling law with Flory coefficient *v*

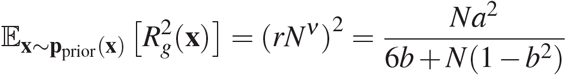

This gives a quadratic equation with the solution

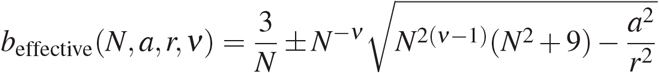

The positive branch is the relevant one to us (the negative branch corresponds to a pathological solutions for small *N*), giving us the result:

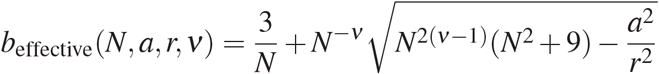

##### C.3.3 Standardizing the translational variance

Currently, the above procedure has diverging marginal variance as *b* → 1. We can arbitrarily retune the *translational variance* of each chain with the following mean-deflation operation which enforces 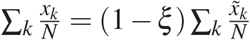:

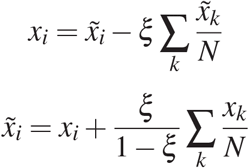

##### C.3.4 Setting the parameters

We have two procedures for setting the value of *ξ*, leading to two different named covariance models:

1. **Pile-of-globs covariance**. Set *a* and *b* to satisfy within-chain *R_g_* scaling of each chain in a complex based on its length. Set *ξ* so that the translational variance of each chain is unity. This will cause chains to have a realistic radius of gyration but pile up at the origin.
2. **Glob-of-globs covariance**. Set *a* and *b* to satisfy within-chain *R_g_* scaling of each chain in a complex based on its length. Set *ξ* per chain by solving for the translational variance that also implements the correct whole-complex *R_g_* scaling as a function of the number of residues. This will cause chains to preserve a realistic complex-level radius of gyration and also intra-chain radius of gyration that scales as that of individual globular proteins.

##### C.3.5 Factorization of the matrix

When also including a centering transform, we can factorize the matrix **R** as a product of three matrices, which can be useful for computing inverses and transposes:

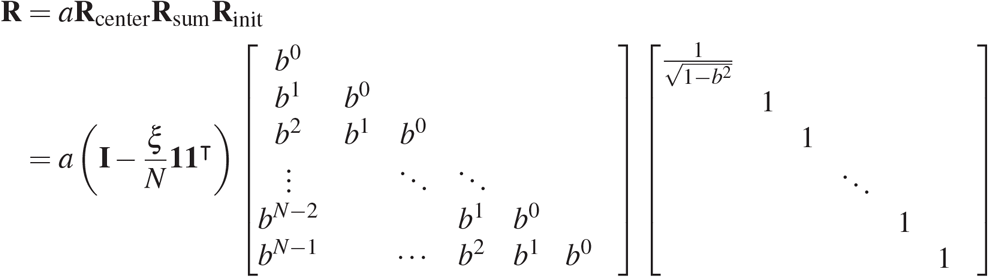

#### C.4 Inverse covariance and intuition

Ignoring the translational rescaling and by numerical investigation, it appears that the precision matrix Σ^−1^ = (**RR**^⊤^)^−1^ is

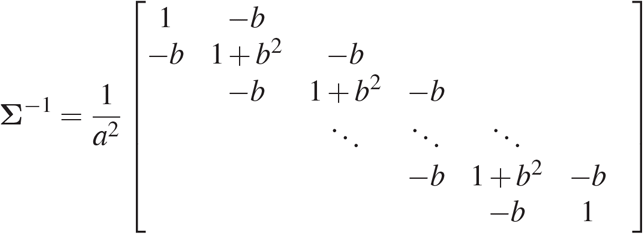

We can decompose this as the sum of terms for the precision of Brownian motion (Chain Laplacian), for a spherical Gaussian (Identity matrix), and some nuisance boundary conditions

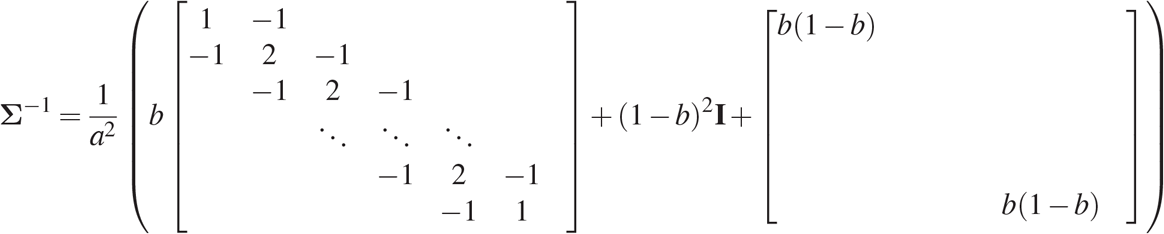

## D Random Graph Neural Networks

Prior approaches to predicting or generating protein structure have relied on neural network architectures with 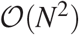 or 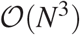 computational complexity [Jumper et al., 2021, Anand and Achim, 2022, Trippe et al., 2022], in part motivated by the need to process the structure at multiple length scales simultaneously and/or to reason over triples of particles as is done during distance geometry methods. Here we introduce an effective alternative to these approaches with sub-quadratic complexity by combining Message Passing Neural Network [Gilmer et al., 2017] layers with random graph generation processes. We design random graph sampling methods that reproduce the connectivity statistics of efficient *N*-body simulation methods, such as the Barnes-Hut algorithm [Barnes and Hut, 1986].

### D.1 Background: efficient *N*-body simulation

One of the principal lessons of computational physics is that *N*-body simulations involving 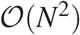 dense interactions (e.g. gravitational simulations and molecular physics) can often be effectively simulated with only 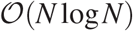-scaling computation. Methods such as Barnes-Hut [Barnes and Hut, 1986] and the Fast Multipole Method take advantage of a common particular property of (and inductive bias for) physical systems that *more distant interactions can be modeled more coarsely for the same level of accuracy*. For example, in cosmological simulations, you can approximate the gravitational forces acting on a star in a distant galaxy by approximating that galaxy as a point at its center of mass.

So far, most *relational* machine learning systems [Battaglia et al., 2018] for protein structure have tended to process information in a manner that is either based on local connectivity (e.g. a *k*-Nearest Neighbors or cutoff graphs) [Ingraham et al., 2019] or all-vs-all connectivity [Jumper et al., 2021, Anand and Achim, 2022, Trippe et al., 2022]. The former approach is natural for highly spatially localized tasks such as structure-conditioned sequence design and the characterization of residue environments, but it is less clear if local graph-based methods can effectively reason over global structure in a way that is possible with fully connected Graph Neural Networks, such as Transformers [Vaswani et al., 2017]. Here we ask if there might be reasonable ways to add in long-range reasoning while preserving sub-quadratic scaling simply by random graph construction.

#### Related work

Our method evokes similarity to approaches that have been used to scale Transformers to large documents by combining a mixture of local and deterministically [Child et al., 2019] or randomly sampled long-range context [Zaheer et al., 2020]. Distant-dependent density of context has also been explored in multiresolution attention for Vision transformers [Yang et al., 2021] and in dilated convolutional neural networks [van den Oord et al., 2016].

### D.2 Random graph generation

We propose to build scalable graph neural networks for molecular systems by sampling random graphs that mix short and long-range connections. We define the graph 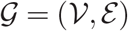 where 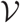 is the node set and 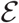 is the edge set. A protein can be represented as a point set **x** ∈ ℝ^*N*×3^. We define the process of constructing the geometric graph as 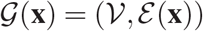 with 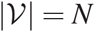. Different from the usual graph construction scheme, the edges are generated stochastically, and 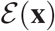 describes the process. We consider schemes in which edges for each node are sampled without replacement from the set of possible edges, weighted by an edge propensity function based on spatial distance (Fig. 3). In practice, we implement this weighted sampling without replacement using Gumbel Top-*k* sampling Kool et al. [2019] (Algorithm 1). Throughout this work, we use hybrid graphs which include the 20 nearest neighbors per node together with 40 randomly sampled edges under the inverse cubic edge propensity function so that both short-range and long-range interactions are sampled with appropriate rates.

**Algorithm 1.**
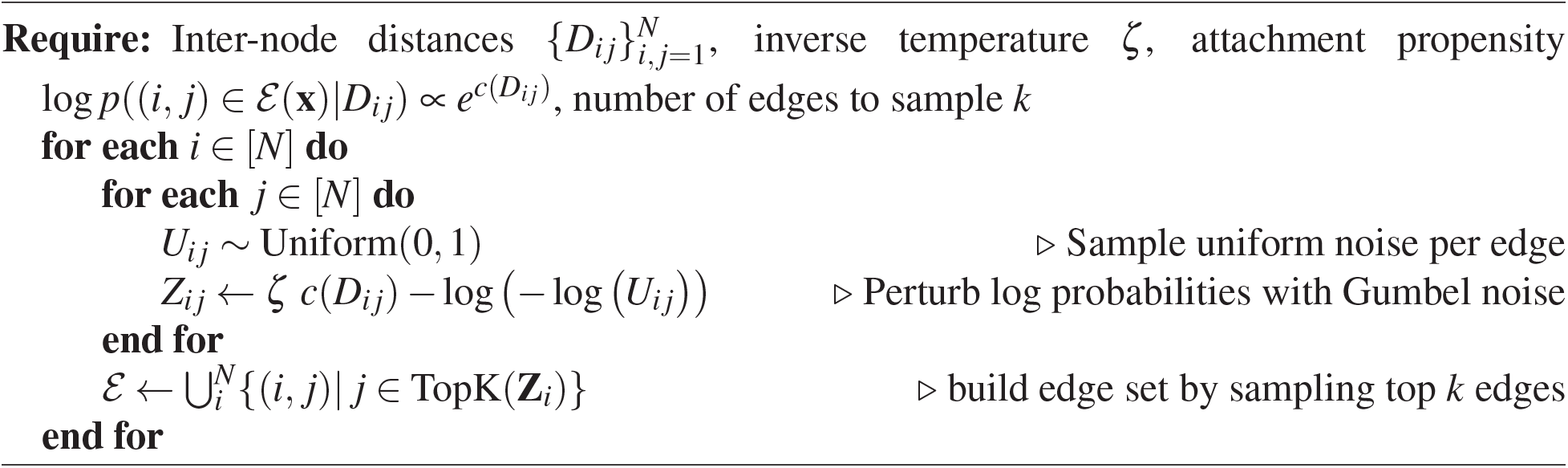
Random graph generation

### D.3 Computational complexity

Under the inverse cubic attachment model, the cumulative edge propensity as a function of distance will scale as 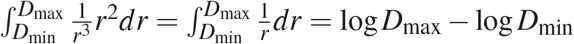. As we increase the total size (radius) of the system by *D*_max_, we only need to increase the total number of of edges per node by a factor of log *D*_max_ to keep up with the increase in total edge propensity (and to therefore ensure that increasingly distant parts of the system do not “steal” edge mass from closer parts of the system). This means that, even if we were to scale to extremely large systems such as large, solvated molecular dynamics systems with millions of atoms, the total amount of computation required for a system of *N* atoms will scale as 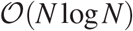. In practice, we found that for protein sizes considered in this work (complexes containing up to 4000 residues^2^) it was sufficient to simply set the number of edges per node to a constant *k* = 60, which means that the graph and associated computation will scale within this bounded size as 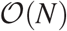. This is a considerable improvement on previous approaches for global learning on protein structure such as methods based on fully connected graph neural networks Trippe et al. [2022] 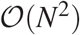 or Evoformer-based approaches [Jumper et al., 2021] which scale as 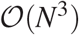. These sparse graphs also combine favorably with our method for synthesizing updated protein structures based on predicted inter-residue geometries (Section E).

**Supplementary Figure 3:**
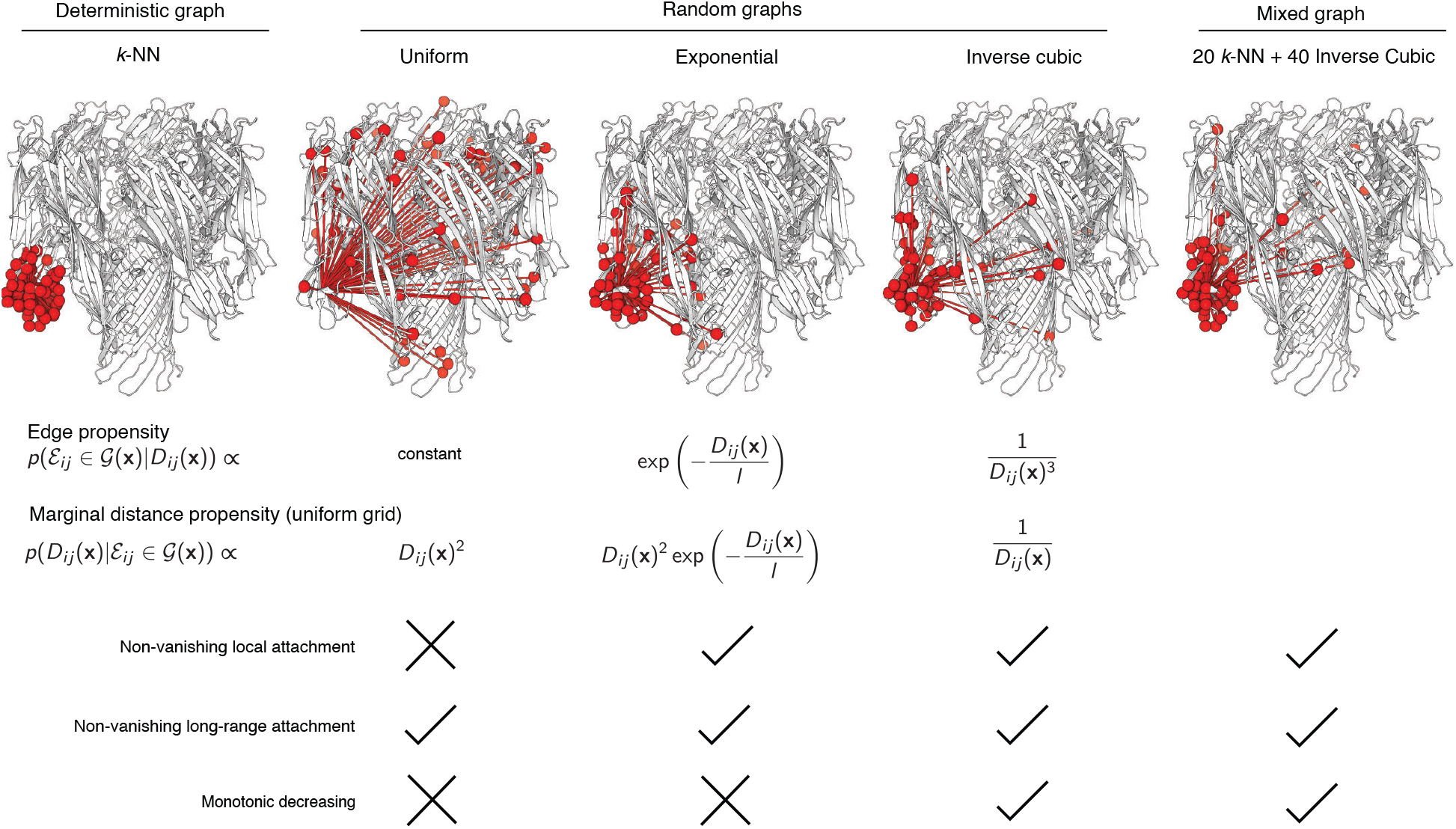
Random graphs with distance-weighted attachment efficiently capture long-range context. Contemporary graph neural networks for learning from molecular systems achieve efficiency via spatial locality, e.g. with a spatial *k*-Nearest Neighbors graphs or cutoff graph (top left, 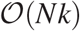). We propose methods that retain this efficiency while incorporating long-range context through random edge sampling weighted by spatial distance (middle columns). We consider three different graph sampling schemes: (i) Uniformly random sampling (middle left) introduces long-range context but at the expense of vanishing local attachment. (ii) Exponential distance weighting (middle center), which can be related to dilated convolutions [van den Oord et al., 2016], includes both short- and long-range attachment but introduces a *typical* length scale as it induces Gamma-distributed distances. (iii) Inverse cubic distance weighting (middle right), which is the effective connectivity scaling of fast *N*-body methods such as Barnes-Hut [Barnes and Hut, 1986], retains a balance of both short and long-term distances with a marginal distance propensity that gently and monotonically decays with *D*. In practice, we combine inverse cubic sampled random graphs with deterministic *k*-NN graphs to guarantee coverage of the *k* closest nodes while adding in long-range context (top right).

## E Equivariant Consensus Structure from Weighted Inter-residue Geometries

### E.1 Background and motivation

Prior neural network layers for generating molecular geometries in proteins have typically relied on either (i) direct prediction of backbone internal coordinates (i.e., dihedral angles) [AlQuraishi, 2019, Wu et al., 2022a], which incurs accumulating errors along the chain in the form of “lever effects” that hinder performance beyond small systems; (ii) prediction of inter-residue geometries followed by offline optimization [Anand and Huang, 2018, Senior et al., 2020], which builds on the successes of predicting protein structure from contacts [Marks et al., 2012] but is difficult to make end-to-end trainable; or (iii) iterative local coordinate updates based on the entire molecular system [Jumper et al., 2021].

In principle, protein structures arise from a balance of competing intra-molecular forces between atoms in the polymer. Indeed, protein structure can be thought of as the solution to a constraint satisfaction problem across multiple length scales and with many kinds of competing interactions. It is therefore natural to think about protein structure prediction as a so-called “Structured Prediction” problem [Belanger and McCallum, 2016] from machine learning, in which predictions are cast as the low-energy configurations of a learned potential function. Structured Prediction models often learn efficiently because it is usually simpler to express a system in terms of its constraints than to directly characterize the solutions to these constraints. This perspective can be leveraged literally for molecular geometries via differentiable optimization or differentiable molecular dynamics [Ingraham et al., 2018, Schoenholz and Cubuk, 2020, Wang et al., 2020], but these approaches are often unstable and can be cumbersome to integrate as part of a larger learning system.

### E.2 Equivariant structure updates via convex optimization

Here we introduce a novel framework which realizes the benefits of inter-residue geometry prediction and end-to-end differentiable optimization in an efficient form based on convex optimization. We show how predicting pairwise inter-residue geometries as pairwise roto-translation transformations with anisotropic uncertainty induces a convex optimization problem which can be either locally solved analytically admits a dast iteration scheme for a global consensus configuration. Throughout this section we will refer to the coordinate frames of residues with a notation that is similar to that used in AlphaFold2 [Jumper et al., 2021], but with rotations **R** replaced with **O** (for **O**rientation) as in Ingraham et al. [2019]. The functions for synthesizing the backbone structure from residue poses (i.e., StructureToTransforms and TransformsToStructure) are those described in the supplementary information of [Jumper et al., 2021].

The key idea of our update is that we ask the network to predict a set of inter-residue geometries **T**_*ij*_ together with confidences *w_ij_* (which will initially be simple but can be extended to anisotropic uncertainty) and we then attempt to either fully or approximately solve for the consensus structure that best satisfies this set of pairwise predictions.

#### Transform preliminaries

Let **T** = (**t**, **O**) ∈ SE(3) be a transformation consisting of a translation by vector **t** ∈ ℝ^3^ followed by a rotation by an orthogonal matrix **O** ∈ SO(3). These transformations form a group with identity, inverse, and composition given by

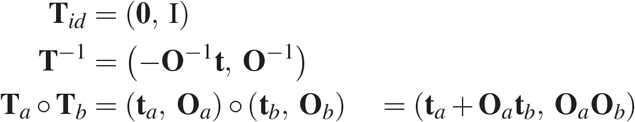

We denote the transformation to the frame of each residue *a* as **T**_*a*_, and denote the relative transformation from residue *a* to residue *b* as

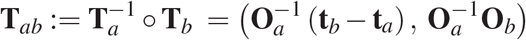

These relative transformations satisfy equations

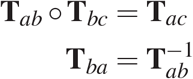

#### Convex problem

Given a collection of pairwise inter-residue geometry predictions **T**_*ij*_ and confidences *w_ij_* ∈ ℝ, we score a candidate structure 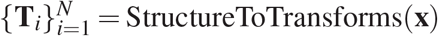 via a weighted loss *U* that measures the agreement between the current pose of each residue **T**_*i*_ and the predicted pose of the residue from the frame of each neighbor **T**_*j*_:

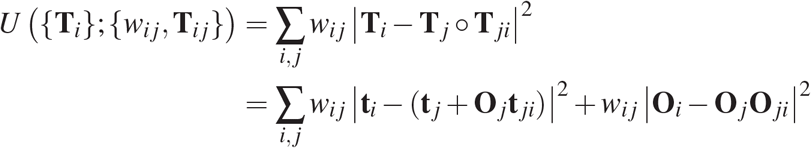

We wish to optimize each local pose **T**_*i*_ with neighbors fixed as

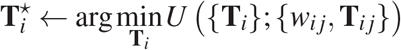

This problem of finding the local “consensus pose” 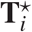 is a convex optimization problem, the solution to which can be analytically realized as a weighted average with projection,

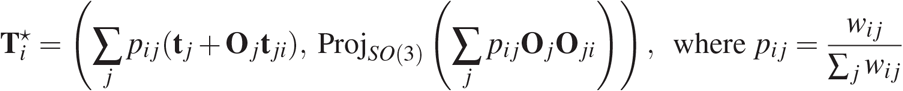

where the projection operator is accomplished via SVD as in the Kabsch algorithm [Kabsch, 1976] for optimal RMSD superposition. If we iterate this update multiple times to all positions in parallel, we obtain a parallel coordinate descent algorithm. Putting this together, we parameterize our denoising function as

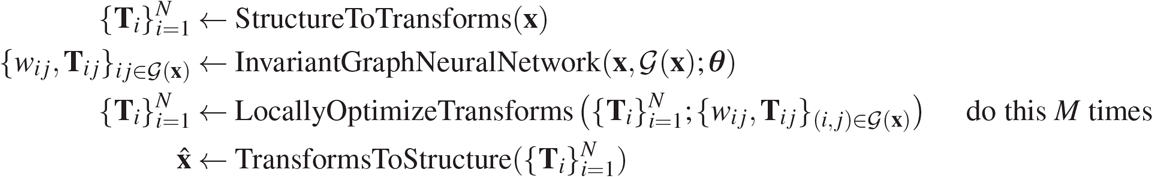

#### Extensions to anisotropic uncertainty models

The above iteration leverages an *isotropic* uncertainty model in which the error model for the translational component is spherically symmetric and coupled to the uncertainty in the rotational component of the transform. We consider two forms of anisotropic uncertainty: in the first, two-parameter version, we decouple the weight *w_ij_* into separate factors for the translational and rotational components of uncertainty, 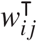 and 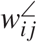, respectively. This makes intuitive sense when, for example, the network will possess high confidence about the relative position of another residue but not its orientation.

In a second and more sophisticated form of anisotropic uncertainty, we extend this framework to ellipsoidal error models bespoke to each *ij*, while retaining a closed form iteration update using approaches from sensor fusion. We parameterized this anisotropic error model by separating this precision term *w* into three components: 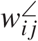 for rotational precision and two components for position: 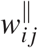 for radial distance precision, and 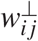 for lateral precision. The radial and lateral precision terms are each eigenvalues of the full 3×3 precision matrix **P**_*ij*_ for translation errors (i.e., inverse covariance matrix under a multivariate normal error model):

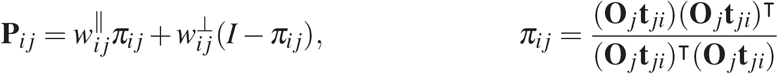

where *π_ij_* is the projection matrix onto the radial direction from **t**_*j*_ to the predicted position **t**_*j*_ + **O**_*j*_ **t**_*ji*_ of **t**_*i*_, and I–*π_ij_* is the projection matrix onto lateral translations (spanned by the remaining two eigenvectors). These anisotropic terms finally combine as

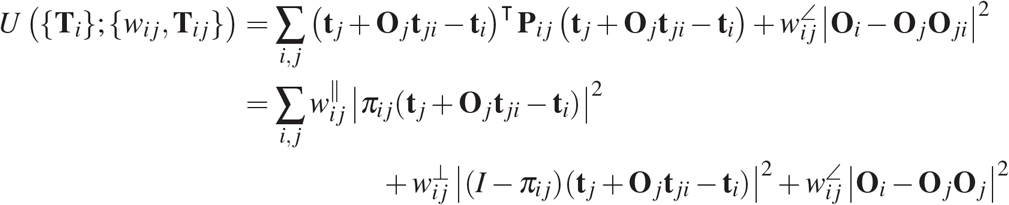

As we expect the radial precision to always exceed the lateral precision, our neural predictor outputs three positive parameters 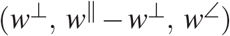. Whereas the isotropic objective above is solved by weighted averaging, the anisotropic translation part of this objective is solved by a standard Gaussian product operation from sensor fusion [Murphy, 2007]:

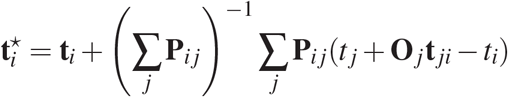

Supplementary Figure 4 illustrates this anisotropic Gaussian fusion operation.

#### Extension as a generalization of the AlphaFold Structure Module

If an additional “dummy” edge is added that is connected to the current state of **T**_*i*_, then this transform will serve an identical role to the predicted frame updates in AlphaFold2. Thus, our framework can be cast as a generalization of this family of backbone updates that opens up opportunities for complex fusion of predicted interrresidue geometries.

**Supplementary Figure 4:**
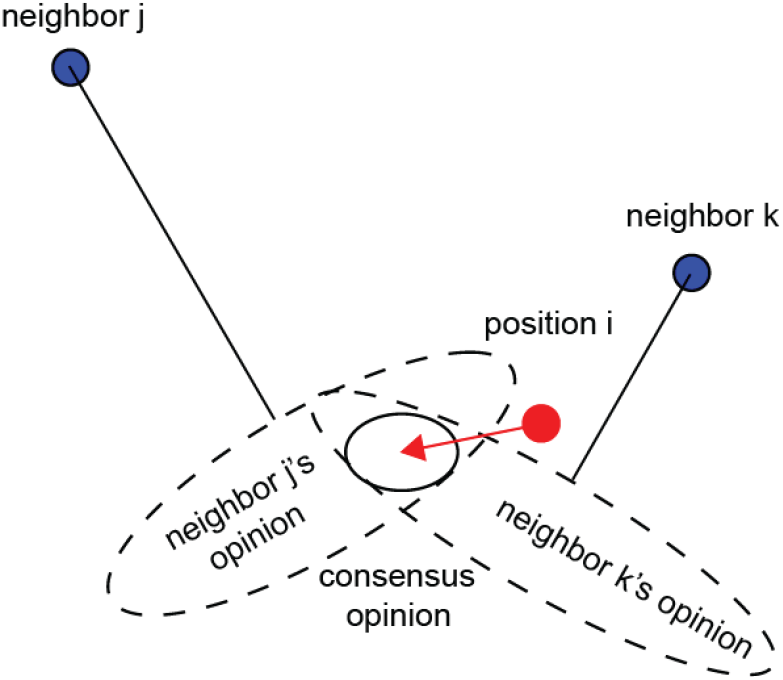
Anisotropic consensus update. Position *i* is forced towards its consensus position which is the mean of a fusion of anisotropic Gaussians. Here we visualize the covariance ellipsese of component the Gaussians, i.e. the inverses of the precision matrices predted by our network.

### E.3 Extension to equivariant prediction of all backbone atoms

The above updates predict rigid residue poses {**T**_*i*_} but our diffusion model (Appendix C) requires independent prediction of all backbone heavy atoms. We can straightforwardly augment the above predictions in an equivariant manner by predicting from every node embedding *local coordiates* for each atom position relative to the parent residue pose. To ease learning, we cast these as residual updates from the ideal geometry positions of each backbone heavy atom. To build the final atomic structure, we simply compose these right-compose these local coordinate predictions with the parent poses. These predictions will be equivariant because they are right-composed with the parent residue poses, which are equivariant because they are built from relative, equivariant geometric transformations off of the initial geometry.

## F Chroma architecture

Supplementary Figure 5 provides a diagram of the Chroma architecture, which includes the *backbone network*, the *design network*, and the underlying graph neural network components. We list important hyperparameters for the backbone network in Supplementary Table 2 and for the design network in Supplementary Table 3. We design sequences by extending the framework of [Ingraham et al., 2019] and factorizing joint rotamer states autoregressively in space, and then locally autoregressively per side-chain *χ* angle within a residue as done in [Anand et al., 2022]. For the sequence decoder, we explore both autoregressive decoders of sequence (pictured in fig. 5) and conditional-random field decoding of sequence, which was also explored in concurrent work [Li et al., 2022].

### Graph Neural Network

All of our neural network models are based on graph neural networks that reason over 3D structures of proteins by transforming them into attributed graphs built from rigid transformations (*SE*(3)) invariant features. This approach has been pursued in several prior works [Ingraham et al., 2019], and our primary architectural innovations on those models are two-fold:

- We propose *random graph neural networks* that add in long-range connections and reasoning while preserving subquadratic / quasi-linear computational complexity (Appendix D)
- We introduce a method for efficiently and differentiably generating protein structures from predicted inter-residue geometries based on parallel coordinate descent (Appendix E)

**Supplementary Figure 5:**
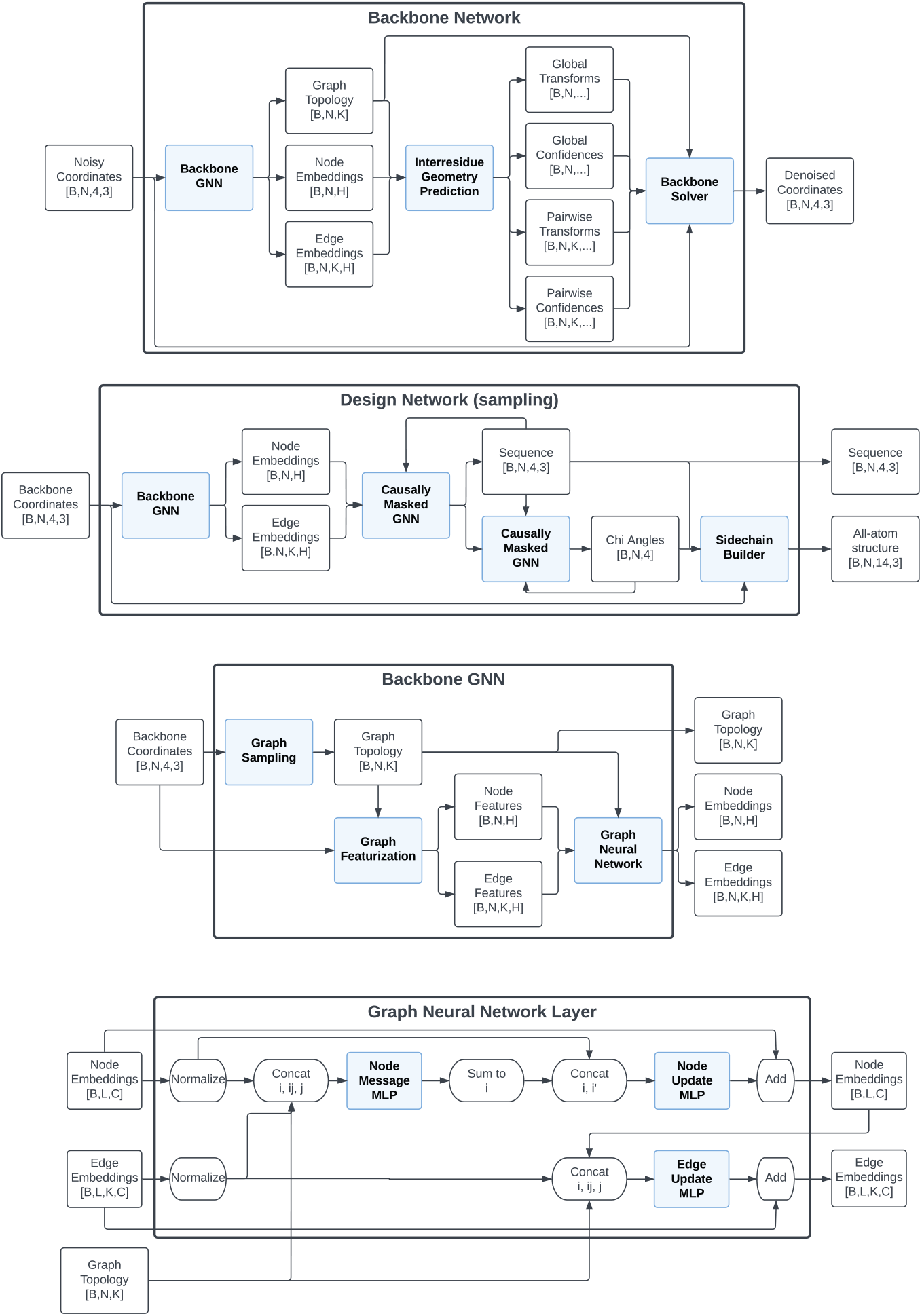
Chroma is composed of graph neural networks for backbone denoising and sidechain design.

**Table 2:** Hyperparameters of backbone network configuration.

| Category | Hyperparameter | Value in Backbone Network A | Value in Backbone Network B |
| --- | --- | --- | --- |
| Diffusion model | Covariance model | Pile-of-Globs | Glob-of-Globs |
|  | Noise schedule | Log-linear SNR (-7,13.5) | Log-linear SNR (-7,13.5) [Kingma et al., 2021] |
| Graph Featurization | Node Features | Internal Coordinates | Internal Coordinates |
| | Edge Features | Distances, Interresidue transforms, $\vert i - j\vert $, same_chain | Distances, Interresidue transforms, $\vert i - j\vert $, same_chain |
|  | Number of edges per node, k | 60 | 60 |
|  | Number of kNN edges | 20 | 20 |
|  | Number of Inverse Cubic edges | 40 | 40 |
| Graph Neural Network | Number of GNN layers | 12 | 12 |
|  | Node Embedding Dimension | 512 | 512 |
|  | Edge Embedding Dimension | 256 | 256 |
|  | Node MLP hidden dimension | 512 | 2048 |
|  | Edge MLP hidden dimension | 128 | 128 |
|  | Dropout p | 0.1 | 0.1 |
| Backbone Solver | Uncertainty model | Isotropic (1-parameter) | Decoupled (2-parameter) |
|  | Number of iterations | 3 | 10 |
| Loss Function | Whitened ELBO weight | 1 | 1 |
|  | X-space pseudo-ELBO weight | 1 | 1 |
|  | X-space units | Nanometers | Nanometers |
| Total Number of Parameters |  | 18.6M | 94.1M |

**Table 3:** Hyperparameters of design network configuration.

| Category | Hyperparameter | Value in Design Network A | Value in Design Network B |
| --- | --- | --- | --- |
| Graph Featurization | Node Features | Internal Coordinates | Internal Coordinates |
| | Edge Features | Distances, $\vert i - j\vert $, same_chain | Distances, $\vert i - j\vert $, same_chain |
|  | Number of edges per node, k | 30 | 30 |
|  | Number of kNN edges | 30 | 30 |
|  | Number of inverse cubic edges | 0 | 0 |
| Graph Neural Network | Number of GNN layers | 6 | 3 |
|  | Node embedding dimension | 128 | 128 |
|  | Edge embedding dimension | 128 | 128 |
|  | Node MLP hidden dimension | 512 | 512 |
|  | Edge MLP hidden dimension | 128 | 128 |
|  | Dropout p | 0.1 | 0.1 |
| Sequence decoder | Type | Potts model | Autoregressive |
| Chi decoder | Number of chi bins | N/A | 36 |
| Total Number of Parameters |  | 3.7M | 9.2M |

## G Training

### G.1 Dataset

The PDB was queried (on 2022-03-20) for non-membrane X-ray protein structures with a resolution of 2.6 *A* or better. Structures with homologous sequences were removed by assigning each chain sequence to a cluster ID using USEARCH [Edgar, 2010] at a 50% sequence identity threshold and removing entries with chain cluster ID completely found in another entry. An additional set of 1726 non-redundant antibody structure cluster using 90% sequence identity was added to the reduced set. All 28819 remaining structures were transformed to their biological assembly by favouring assembly ID where the authors and software agreed, followed by authors and finally by software only. Missing side-chain atoms were added with pyRosetta [Chaudhury et al., 2010]. An 80/20/20 train, validation and test splits were generated by minimizing the sequence similarity overlap using entries of PFAM family ID, PFAM clan ID [Mistry et al., 2021], UniProt ID [Bateman et al., 2020] and MMSEQ2 cluster ID at a 30% threshold [Steinegger and Söding, 2017]. A graph pruning method was used to minimize shared label overlap between all splits. Briefly, a graph is built where each PDB entry is represented by a node connected to other entries that share at least one identical annotation. Connected sub-graphs are identified and broken apart by iteratively deleting the most central annotations until there are 50 or fewer connected nodes. Using this procedure, the generated test set had 9%, 59%, 82% and 89% of its entries that did not share any PFAM clan, PFAM family, MMSEQ30 cluster, or Uniprot IDs with the training set, respectively. Whereas a random split would have given 0.1%, 10%, 50% and 70% for the same label types, respectively.

### G.2 Optimization

We train the backbone model by optimizing the regularized ELBO loss (Appendix A) with the Adam optimizer [Kingma and Ba, 2014] and leverage data parallelism across 8 GPUs. We train the design networks by optimizing the sequence (pseudo)likelihoods and chi angle likelihoods with Adam on a single GPU.

## H Evaluation

### H.1 Unconditional samples

Two sets of unconditional protein samples were generated for display and analysis with Chroma. Both sets used the same parameters: 200 steps, *λ*_0_ = 10, and *ψ* = 2. Of these two sets, one was comprised of single chain proteins, and the other, of multi-chain proteins. The single chain set contained 50 thousand samples and the lengths were drawn from a ”1/Length” distribution where the probability of a protein chain’s length was proportional to one over its length. The multichain set contained 10 thousand samples and length distribution of each chain was determined empirically from the chain length statistics from complexes in the PDB. Specifically, a random protein complex was drawn from the PDB and the number of chains and length of each chain for the random sample was determined from that random PDB complex. In order to show typical non-cherry picked random samples from the model we provide supplementary Figure 6 for single chain and Figure 7 for multi-chain examples.

### H.2 Backbone geometry statistics

To evaluate the structural validity of Chroma generated single chain structures, they were characterized based on secondary structures and residue interactions alongside a non-redundant subset of PDB database (**Table 4**). The distribution of secondary structures (*α*-helix, *β*-strands, and coil) was evaluated using Stride [Frishman and Argos, 1995]. Residue interaction was determined by any pairwise residue (C-*α* to C-*α*) with an Euclidean distance less than 8 *Å*. Mean and long-range residue contact were computed. Contact order [Ivankov et al., 2003] and radius of gyration [Tanner, 2016] were computed and length normalized according to their corresponding empirical power laws. All metrics except for secondary structures are normalized for Fig. 2b.

### H.3 Novelty and structural homology

The novelty of Chroma-generated samples was assessed by comparing their structures to natural protein folds. Using TMalign [Zhang and Skolnick, 2005], they were each aligned to the 32k structurally conserved domains from CATHdb S40 set [Sillitoe et al., 2021] and filtered for TMscore greater than 0.5 when normalized by the shortest sequence. The number of domains needed to cover at least 80% of the query was greedily determined by identifying the hits with the highest number of residues within 5 *Å* of the query that wasn’t already covered. The number of domains required increases with query size given that CATH domains typically have a length ranging between 50 and 200 amino acids. As a baseline, we ran the test set with the same algorithm.

**Supplementary Figure 6:**
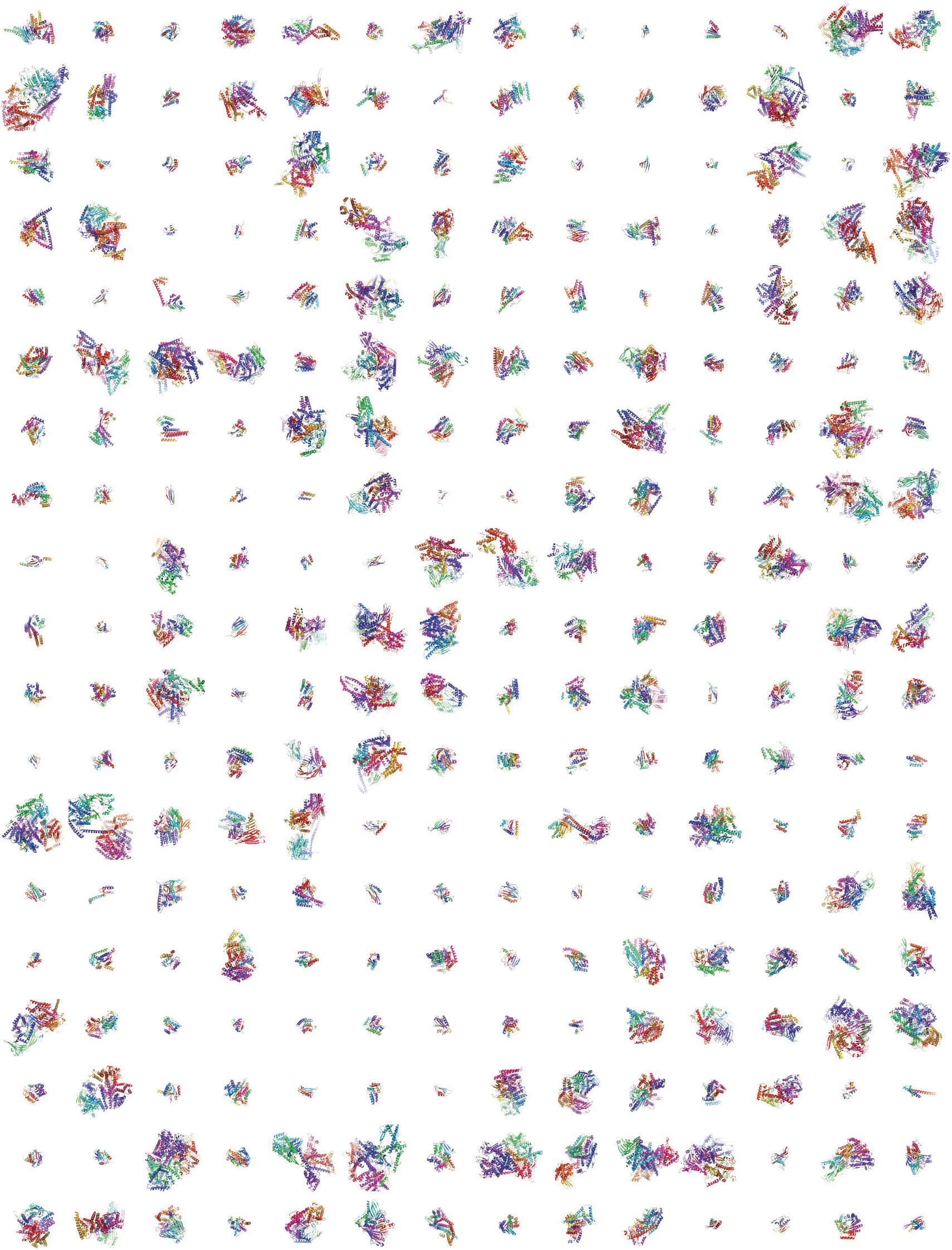
Random single chain samples from backbone model A.

Single chain structures from Chroma and the test set were embedded in 31 Gauss Integral dimensions using the pdb2git program from the Phaistos suite [Harder et al., 2012] [Borg et al., 2009]. It failed to embed structures with chain breaks or with protein lengths greater than 876. The final set of 6492 generated structures and 561 natural folds were projected to a 2 dimensions space using UMAP [McInnes et al., 2018] with default parameters of 25 neighbours and a minimal distance of 0.5.

**Supplementary Figure 7:**
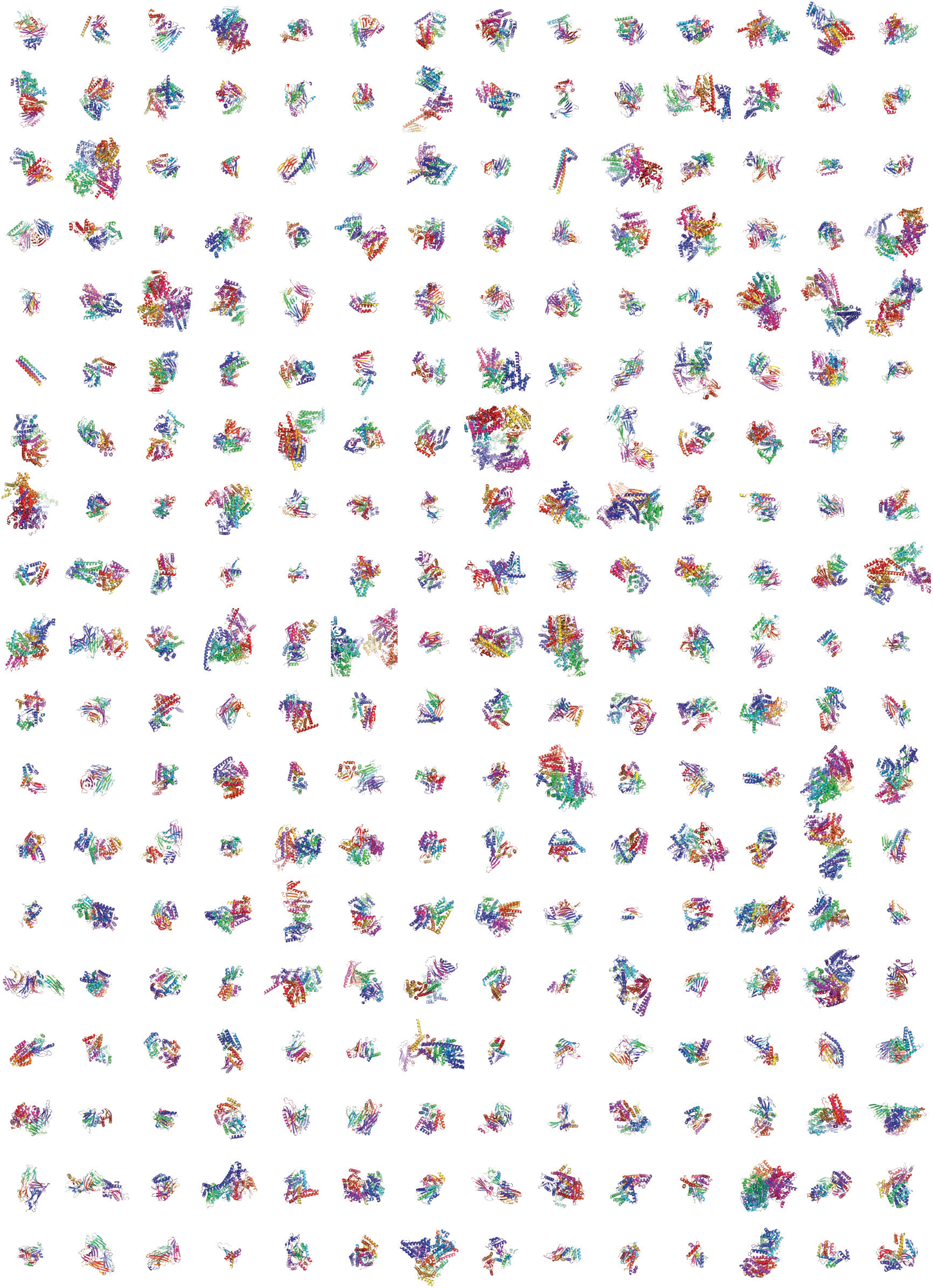
Random complex samples from backbone model A.

**Table 4:**
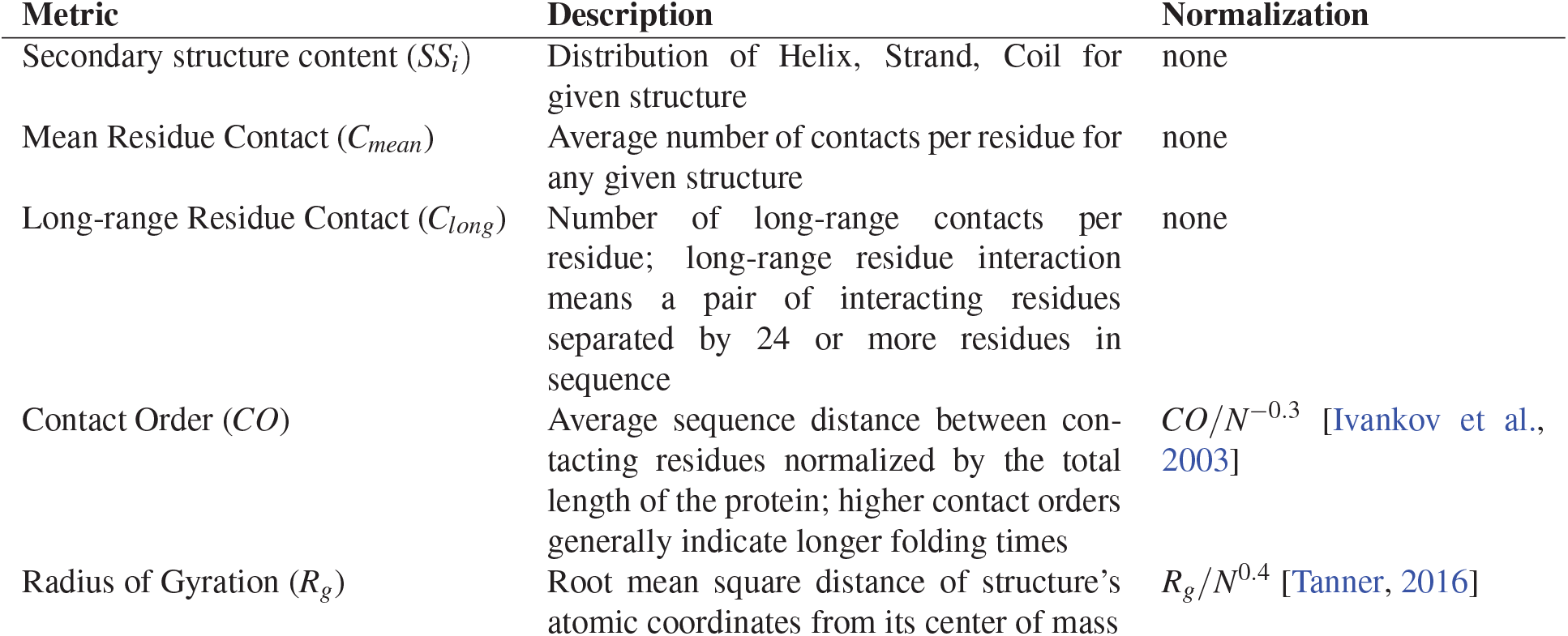
Metrics to describe backbone geometry of structures.

**Table 5:**
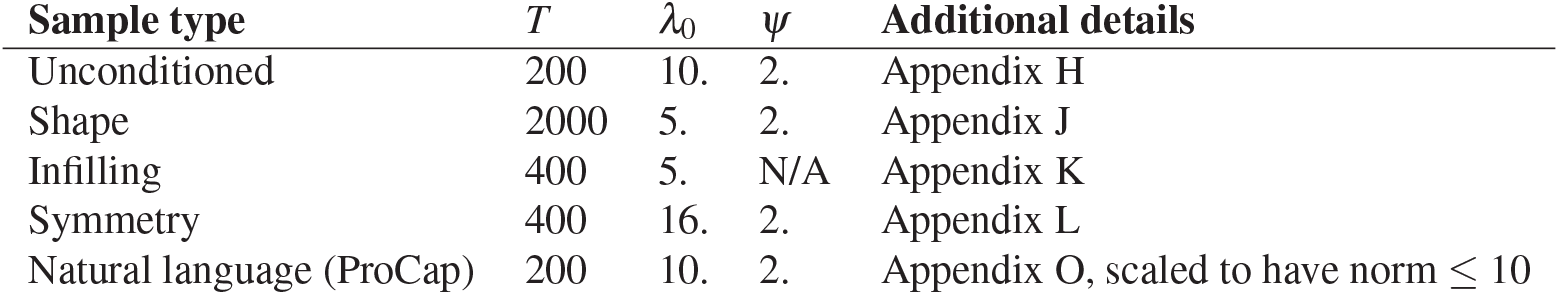

| Sample type | $T$ | $\lambda_0$ | $\psi$ | Additional details |
| --- | --- | --- | --- | --- |
| Unconditioned | 200 | 10. | 2. | Appendix H |
| Shape | 2000 | 5. | 2. | Appendix J |
| Infilling | 400 | 5. | N/A | Appendix K |
| Symmetry | 400 | 16. | 2. | Appendix L |
| Natural language (ProCap) | 200 | 10. | 2. | Appendix O, scaled to have norm $\leq 10$ |

### H.4 Structure prediction-based designability

Structures were generated using Chroma with *λ*_0_ = 10. Sequences conditioned on generated structures were designed by using our sequence design module with a Potts decoder to create a pairwise sequence-level energy table representing the sequence landscape compatible with the fold. Sequences were sampled from this Potts model using 10 independent cycles of simulated annealing Monte Carlo (MC), each with 200 · *n* steps with *n* being the length of the protein. The score used in the annealing was the Potts energy plus a flat-bottom restraint energy around the sequence complexity calculated as equation 4 in [Wootton and Federhen, 1993]. The restraining potential linearly penalized sequence complexity dropping below one standard deviation under the mean for a native sequence of equivalent length, and was otherwise zero. For each generated structure, the above MC procedure was run 100 times to produce 100 sequences, each of which were used as input into OmegaFold [Wu et al., 2022b] for structure prediction. For each backbone, the highest obtained TM score was used as the evidence for whether the underlying backbone was designable in our analysis.

### H.5 TERM-based designability

RMSD-based search was performed using an in-house implementation of the method FASST available as part of the open-source software package Mosaist (https://github.com/Grigoryanlab/Mosaist). The method is a close relative of the previously published approach MASTER [Zhou and Grigoryan, 2014, 2020a]. The training and test sets for Chroma were used as the search database and the set of native proteins in this analysis, respectively. Although the test and training sets have been split by chain-level sequence homology, we took further care to exclude any apparent homologues of native TERMs from consideration as matches. To this end, we compared the local 31-amino acid sequence windows around each TERM segment and its corresponding match, with any pairings reaching 60% or more sequence identities not being allowed to participate in a match.

## I Programmability: Overview

### Overview

In principle, the set of proteins satisfying a given set of functional constraints can be described using Bayes’ Theorem,

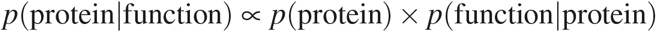

where the posterior distribution of proteins *p* (protein | function) is proportional to the *likelihood* of satisfying the set of functional constraints *p*(function|protein) times the *prior* probability of the protein molecule being functional *p*(protein). This characterization has been appreciated for several decades [Simons et al., 1997], but leveraging it is challenging in practice for two reasons. First, developing tractable and accurate priors over the space of possible proteins has proven extremely difficult owing to the tremendous complexity in a single protein system (a complex can easily have > 10^4^ atoms) and the intractabilities of marginalizing out low level details. Secondly, even with an accurate prior, sampling from the space of polypeptide conformations is extremely difficult as it will generally be a rugged landscape under which global optimization is infeasible.

One potential way to make the difficult inverse problem posed by protein design more tractable is given by contemporary methods from machine learning. In particular, diffusion models make conventionally intractable inference and inverse problems tractable by learning to gradually transform a complex data distribution into a simple and tractable distribution [Sohl-Dickstein et al., 2015, Song and Ermon, 2019]. This has enabled transformative applications in text-to-image modeling [Ramesh et al., 2022, Saharia et al., 2022].

The manner in which diffusion models enable Bayesian inversion can be made especially clear in the continuous-time formulation of Diffusion models, where we can take advantage of the fact that the score functions are independent of normalizing constants and we can therefore express the time-dependent *posterior score* ∇_**x**_ log *p_t_* (**x**|**y**) as the sum of the *prior score* ∇_**x**_ log *p_t_* (**x**) and the *likelihood score* ∇_x_ log*p_t_* (**y**|**x**) because

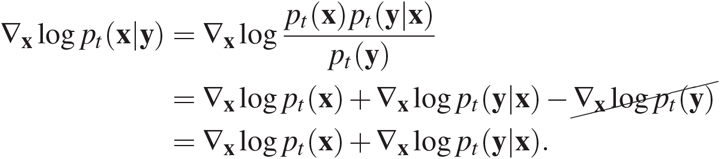

We describe how to take advantage of this via the posterior SDE and ODE in Appendix A.

We note that while we primarily focus on classifier conditioning of backbone conditioning throughout this work, it is also very feasible to extend this to tractable *sequence* classifier conditioning with new discrete sampling methods based gradient-based locally-adjusted MCMC proposals [Grathwohl et al., 2021, Rhodes and Gutmann, 2022].

**Table 6:** Conditioners available to Chroma.

| Conditioner | Dim. | Granularity | Type | Examples and applications |
| --- | --- | --- | --- | --- |
| Sequence | 1D | Residue | Learned | Sequence design & sequence conditioning |
| Domain classifier | 1D | Chain | Learned | Pfam, CATH, Taxonomy |
| Secondary Structure | 1D | Residue | Learned | Topological constraints |
| Distances (contacts) | 2D | Atoms | Analytic | Fold constraints, binder design |
| Sub-structure RMSD | 1D | Atoms | Analytic | Scaffolding-based constraint |
| Sub-structure | 1D | Atoms | Analytic | Structural “in filling” |
| Shape constraint | 1D | Atoms | Analytic | Molecular shape control |
| Symmetry constraint | 1D | Atoms | Analytic | Self-assembling oligomers, e.g. capsids |
| Text caption | 1D | Chain, Complex | Learned | Natural language prompting |

### I.1 Example applications of constraint composition

We list a table of composable constraint models in Table 6. Some practical protein design problems that could be realized through composite constraints under this framework are

***De-novo* binders** Combine (i) substructure conditioning on antigen, (ii) optional scaffold constraint on binder, and (iii) contact constraints on epitope/paratope
**Enzyme miniaturization** Use substructure RMSD to graft an active site into a novel scaffold or known scaffold (via combining with substructure constraints
**Nanostructure control** Use the shape constraint to sample novel designable folds or complexes satisfying arbtitrary shape constraints
**Nanomaterial design** Combine nanostructure control with interfacial binding constraints on periodic boundary conditions

## J Programmability: Distance-based constraints

### J.1 Motivation and problem statement

In some instances, it may be useful to generate diverse protein chain and/or complex structures under the constraint that one or more specific residue pairs be in spatial proximity (i.e., form a “contact”). Such a conditioner could be used, for example, in designing binders, to ensure that the desired binding site is being engaged. Or it could be used to insure some desired topological properties–i.e., the proximity of N- and C-termini (e.g., for ease of circular permutation). Assuming that we are interested in conditioning on a contact between atoms *i* and *j*, we are seeking the probability that the distance between these two atoms in the fully denoised structure is below some desired cutoff *c*, 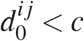, given a noised sample at time *t* and the corresponding distance 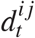.

### J.2 Approach

One approach would be to train a time-dependent classifier *p_t_* (**y**|**x**(*t*)) to classify noisy inputs. For the case of a contact classifier, however, we can directly compute the desired probability analytically. By definition of our forward noise process, the *i*-th coordinate of our protein at time 0 and *t* are related to each other by

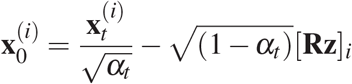

Below we sketch the derivations of the distribution 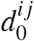 cases of Brownian and globular noise schedules.

#### J.2.1 Brownian noise

Here we have that

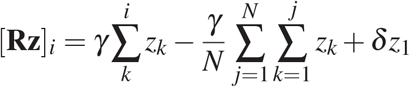

and therefore

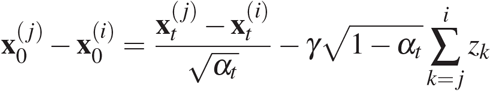

But as 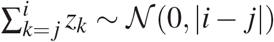 by independence of {*z_i_*}, we have 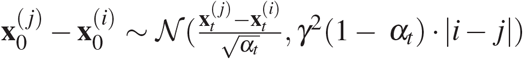, so that:

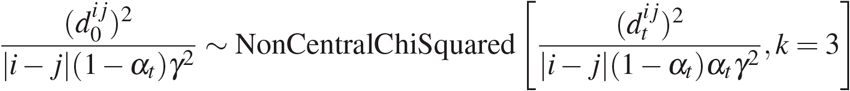

For a contact threshold *c* > 1 we have:

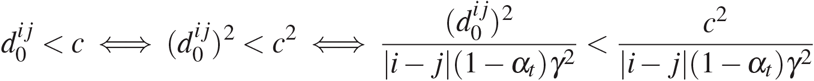

and so we can conclude that 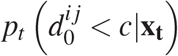 is given exactly by the CDF of the noncentral chi-squared distribution above, evaluated at *c*^2^ [|*i* – *j* | (1 — *α_t_*) *γ*^2^]^-1^.

#### J.2.2 Globular noise

For the globular chain noise process we instead have that

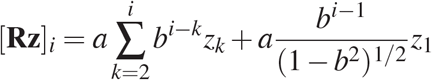

By substituting we see

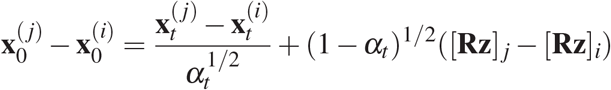

So that 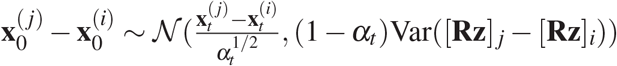. But, assuming *j* > *i*:

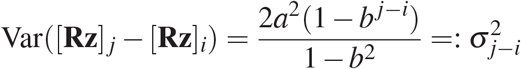

It then follows that

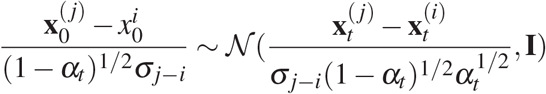

and finally

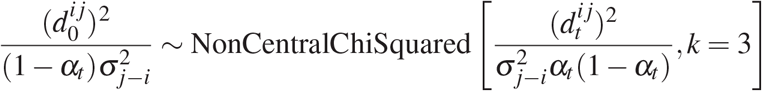

## K Programmability: sub-structure RMSD

### K.1 Motivation and problem statement

It would be very useful for a variety of protein engineering applications to condition structure generation on the presence of a particular structural “motif.” By this we mean an arbitrary substructure, composed of any number of disjoint backbone segments, that we would like to exist within our final generated structure. In practice, such a motif could represent a functional (e.g., catalytic) constellation of residues or a metal/small-molecule binding site—this could be useful for designing enzymes or other functional proteins, by exploring ideas around a core functional mechanism. In another example, the motif could correspond to a “scaffolding” part of the molecule that we would want to preserve—e.g., the binding scaffold that can admit different loop conformations. Or the motif could represent a desired epitope that we would like to faithfully present on the surface of a generated protein in the context of vaccine design. Fig. 8 shows an example motif and two unrelated native protein structures in which this motif is found with low RMSD.

**Supplementary Figure 8:**
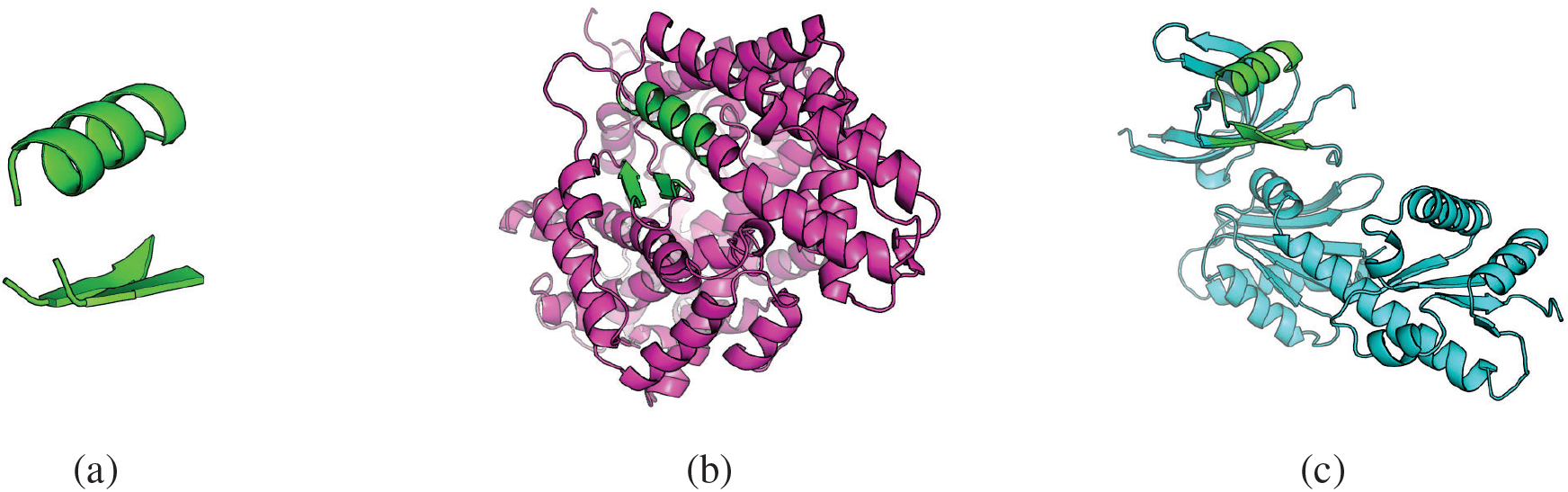
Motifs can occur in entirely unrelated structural contexts. **a,** An example motif composed of three disjoint segments. **b,** PDB entry 3NXQ harbors the motif with a backbone RMSD of 0.45 *Å*. **c,** PDB entry 3OBW harbors the motif with a backbone RMSD of 0.64 *Å.*

The task of determining whether the pre-specified motif is present in a given structure *S* is simple– we can, for example, find the substructure of *S* with the lowest optimal superposition root-mean-squared-deviation (RMSD) to the motif in question and ask whether this RMSD value is below a desired cutoff (this can be done using previously published algorithms [Zhou and Grigoryan, 2015, 2020b]). But what we need for conditional generation is the ability to estimate the probability that the final de-noised structure will harbor the desired motif, given a noisy structure at the current time point in the diffusion.

### K.2 An empirical approach

Specifically, if **x**_*t*_ ∈ ℝ^*N*×3^ is our coordinate array and the forward diffusion process is represented by:

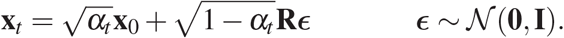

we need to express *p*(*y*|**x**_*t*_)–the probability that **x**_0_ contains the motif given **x**_*t*_, where *y* stands for the condition of motif presence (e.g., as defined by RMSD to a template motif below a desired cutoff). If we define the presence of a motif in terms of optimal-alignment best-fit RMSD being below a cutoff, we need to understand how this RMSD behaves (in a probabilistic sense) as a function of noise. Further, as we will generally not be given where within **x**_*t*_ the motif may be (i.e., we would not know *a priori* the matching between motif atoms and a sub-structure of the target structure), our *p*(*y*|**x**_*t*_) needs to integrate information for the full structure **x**_*t*_ to determine possible motif location(s). Achieving this analytically seems non-trivial. For this reason, here we consider an empirical approach to expressing *p*(*y*|**x**_*t*_).

The goal is to observe the behavior of optimal-alignment best-fit RMSD in practice, as a function of *α_t_*, using a set of reasonable structures and diverse motifs, and find an analytical approximation for its probability distribution. Specifically, given a motif *m* and a structure represented by **x**_*t*_, let *r_t_* represent the RMSD of optimal alignment of *m* onto **x**_*t*_ (i.e., the lowest RMSD between atoms of *m* and any sub-structure of **x**_*t*_), and *r*_0_ represents the RMSD induced by the same matching in the context of structure **x**_0_. We seek to approximate the cumulative distribution function *F*(*r*_0_ – *r_t_*|**x**_*t*_, *α_t_*). With this, we would calculate *p*(*y*|**x**_*t*_) as *p*(*y*|**x**_*t*_) = *p*(*r*_0_ < *σ*|**x**_*t*_) = *p*(*r*_0_ – *r_t_* < *σ* – *r_t_* |**x**_*t*_) = *F*(*σ* – *r_t_* |**x**_*t*_, *α_t_*), where *σ* is the desired RMSD cutoff for classifying the existence of the motif. The procedure in algorithm 2 was run to generate an empirical data set of 10^6^ points to describe the behavior of *r*_0_ – *r_t_*.

**Algorithm 2.**
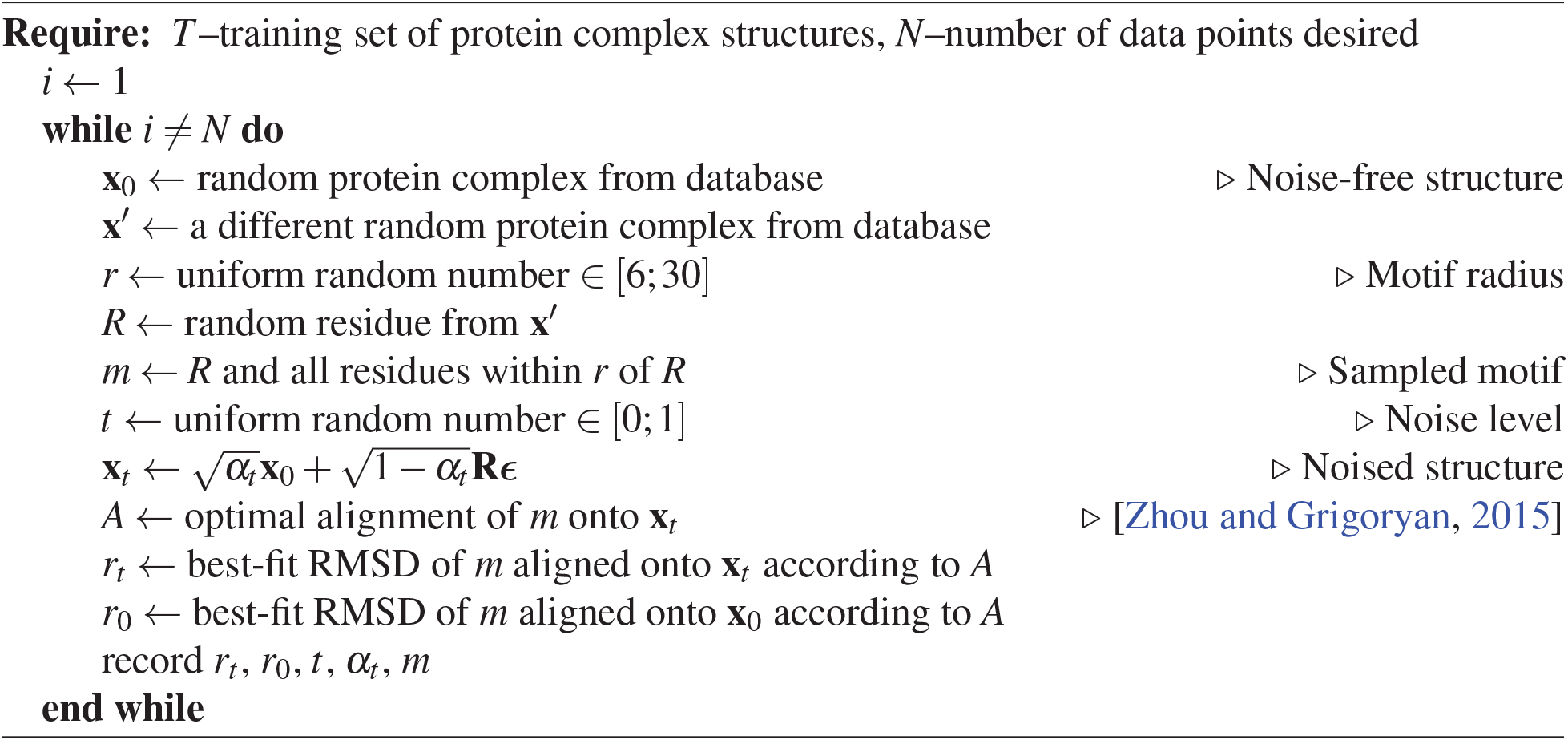
Data generation procedure for RMSD classifier fitting

#### K.2.1 Motif size and complexity dependence

Clearly, the distribution of *r_t_* (and Δ*r_t_* = *r*_0_ – *r_t_*) should depend on *α_t_*. But these distributions should also depend on the size and complexity of the motif. For example, in the extreme case when the motif consists of a single atom, *r_t_* will always be zero. On the other hand, for large and complex motifs, we may expect *r_t_* to increase rapidly with added noise.

The simplest surrogate for motif complexity is its size—i.e., the number of residues it involves. However, under our noise model, the atoms closer to each other in the protein chain will move in a more correlated manner than those that are farther apart. So it should matter whether the motif consists of multiple short disjoint segments matching to far-away (in sequence) portions of the target structure versus a motif consisting of one long contiguous segment. As a purely empirical measure to capture this notion, we propose the following effective length definition:

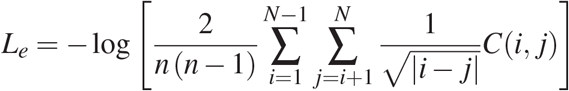

where ***C***(*i, j*) is an indicator function that is 1 if atoms *i* and *j* are part of the same chain and 0 otherwise. The motivation for the inverse square root of the index distance is from Brownian motion (displacement distance growing as the square root of time, here the number of atom hops). And the motivation for ignoring atom pairs from different chains is that these move independently under our noise model. In practice, *L_e_* appears to better explain variation of *r_t_* – *r*_0_ than just pure number of motif residues *L*, despite the fact that overall *L_e_* correlates somewhat closely with log(*L*).

**Supplementary Figure 9:**
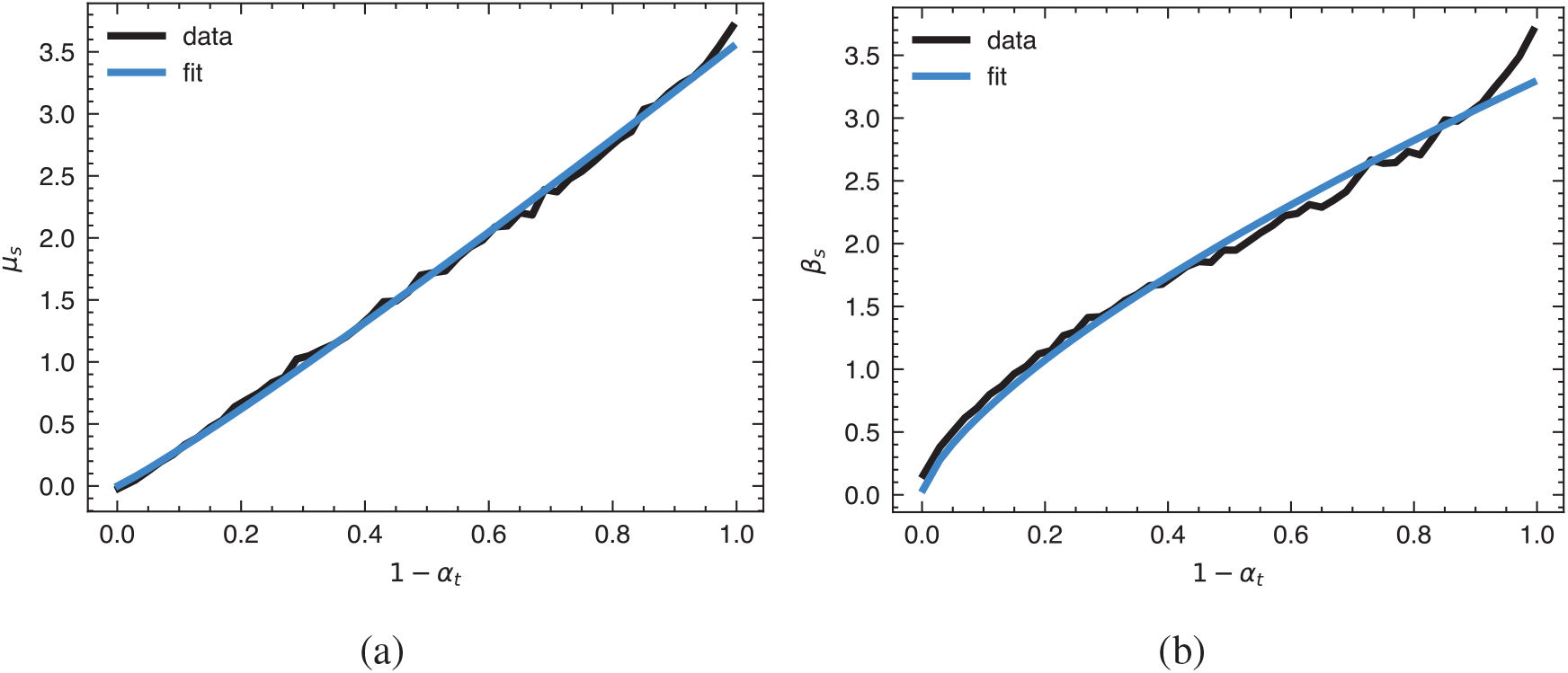
Fitting empirical Gumbel-distribution parameters *μ_s_* and *β_s_* from local *α_t_*-window fits (which capture *L_e_*-dependence of location and scale, respectively) as analytical functions of *α_t_*. Both the location **(a)** and scale **(b)** parameters vary monotonically with *α_t_*, closely following the functional form *k* · (1 – *α_t_*)^*n*^.

#### K.2.2 Distribution of *r*_0_ – *r_t_*

We expect the distribution of *r*_0_ – *r_t_* to depend on *α_t_* and *L_e_.* To get a sense of the general shape of this distribution and its dependence on *αt*, one can take slices of the training data with *α_t_* in different narrow ranges. Inspection and fitting of these *α*_t_-window histograms of *r*_0_ – *r_t_* suggested that the Gumbel family of distribution should work reasonably well for describing the observed variations.

The dependence on *L_e_* can be captured defining the parameters of the Gumbel distribution as functions of *L_e_*. Towards defining a reasonable functional form, we consider extremes. The Gumbel distribution has two parameters–location *μ* and scale *β*. The latter is solely responsible for the variance (i.e., 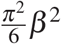) and the mean is contributed to by both (*μ* + *βγ*, where *γ* is the Euler–Mascheroni constant, or approximately 0.577). Clearly, for a motif that only has one atom, we expect Δ*r_t_* to be a delta function at 0, meaning that both *μ* and *β* would be zero. And in general, for small (and simple) motifs we would expect *μ* and *β* to be low, while for large (and complex) motifs we would expect it to be high. Thus, both *μ* and *β* should be monotonically increasing functions of *L_e_* that pass through the origin. Experimentation with different curve families under these criteria, using the overall data likelihood as the objective metric (see below), we arrived at the simple linear parameterization option as being best–i.e., where *μ* = *μ_s_L_e_* and *β* = *β_s_L_e_* with *μ_s_* and *β_s_* being fitting parameters.

### K.3 Fitting procedure

With the parameterization choices above, the fitting approach took the following steps.

#### K.3.1 Fitting individual *α_t_* windows

For 50 equally-spaced *α_t_* windows, fit the observed Δ*r_t_* = *r*_0_ – *r_t_* to Gumbel distributions, whose location and scale parameters linearly depend on *L_e_* of each motif, using likelihood maximization. Specifically, the likelihood function being maximized was:

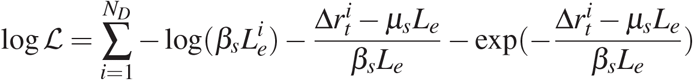

were 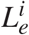 and 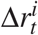 are the effective motif length and Δ*r_t_* is the *i*-th data point, respectively, and *N_D_* is the number of data points. The result of this procedure then estimates *μ_s_* and *β_s_* parameters specific for the current *α_t_* window.

#### K.3.2 Fitting parameters as functions of *α_t_*

We next fit *μ_s_* and *β_s_* as functions of *α_t_* analytically. The functional form chosen for both parameters was *k* · (1 – *α_t_*)^*n*^, ensuring that at *α_t_* = 1 both parameters necessarily become zero (i.e., as the noise level reaches zero, the Δ*r* distribution should approach a delta function). Both the raw locally-fit values of *μ_s_* and *β_s_* and the corresponding analytical fits are shown in Fig. 9.

#### K.3.3 Assessment of fit quality

Given the now fully parameterized *p*(Δ*r_t_* |*α_t_*, *L_e_*), we integrate over *L_e_* in each *α_t_* window to produce the expected distribution of Δ*rt* and compare with the corresponding observed distribution to evaluate the overall goodness of fit. The results, shown in Fig. 10, demonstrate an excellent overall fit. This is especially encouraging given the wide range of motif sizes (anywhere from one to over 350 residues) and numbers of disjoint segments present in the training set (one to five).

### K.4 Conditioning with pre-registration: structural infilling

In some cases, the residue indices of the desired sub-structure in the context of the larger structure are given a priori, for instance, in the case of imputing missing structural information. let 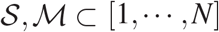 denote the atoms comprising the unknown scaffold and known motif respectively.

#### K.4.1 Related work

Song et al. [2021] presents a replacement method for drawing approximate conditional samples from 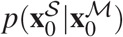 in which one samples a sequence of noised motifs 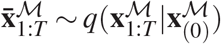, then running diffusion backwards in time but at each time step replacing 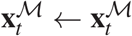 before sampling **x**_*t*-1_ ~ *p*(**x**_*t*-1_ |**x**_*t*_). Trippe et al. [2022] demonstrated that this method introduces irreducible error that is exacerbated by the correlation introduced by *q* and propose a particle-filtering based approach which furnishes arbitrarily accurate conditional samples given sufficient computation. Informally, the error introduced by the replacement method arises from imputing noised motifs that are highly unlikely given the corresponding noised scaffold.

**Supplementary Figure 10:**
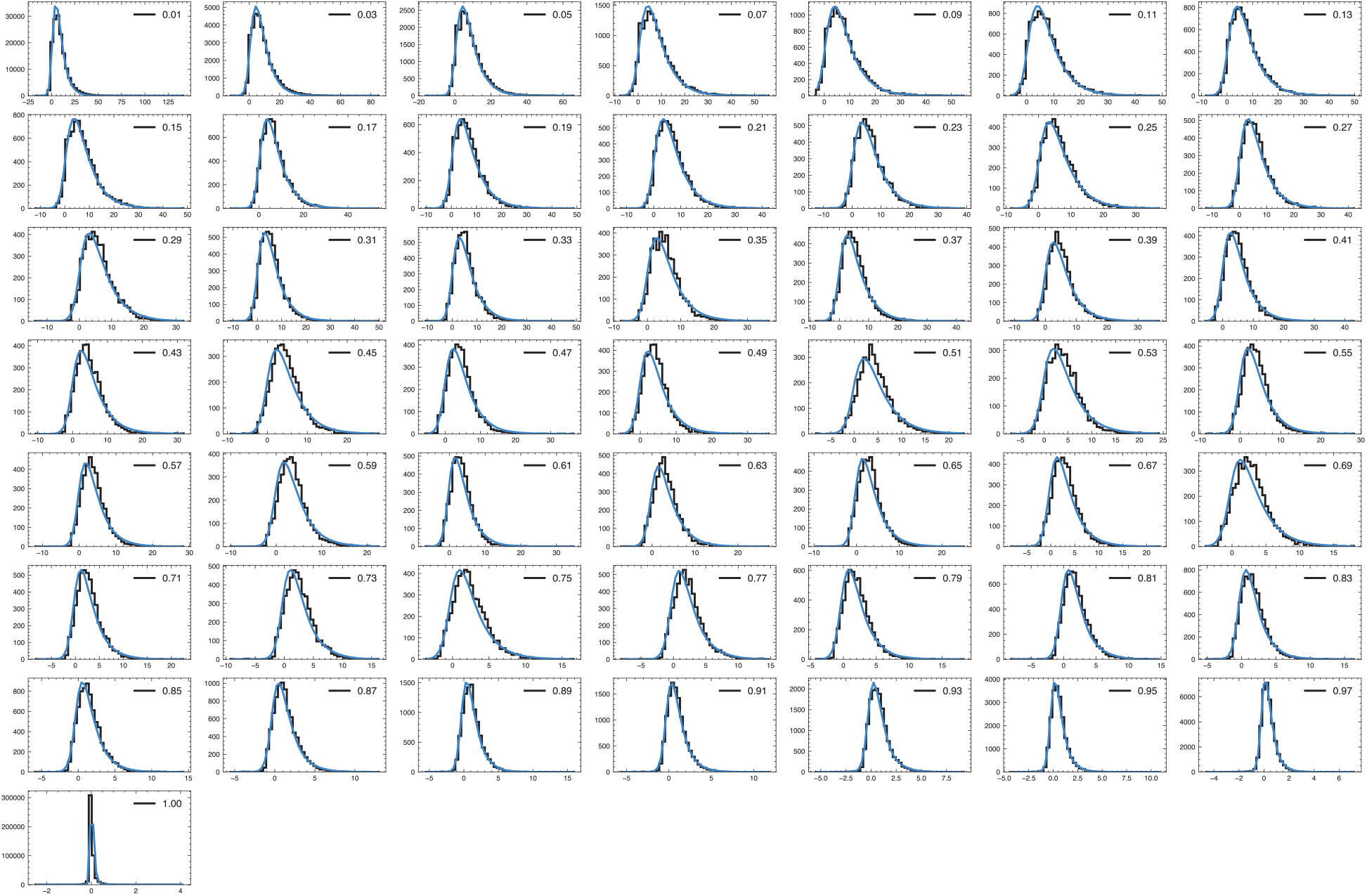
Comparison between expected (based on inferred model parameters) and observed distributions of Δ*r_t_* in different windows of *α_t_*. Observed histograms are show in black and the analytical prediction in blue. Legend indicates the mean *α_t_* value for each window.

#### K.4.2 Method

Given 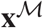, we impute for every contiguous missing fragment 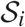 we sample 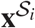 from a brownian bridge with endpoints fixed at 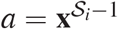 and 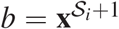 for internal fragments as follows. Let 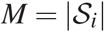, we sample 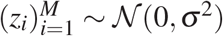, where *σ*^2^ was tuned to 4.0.

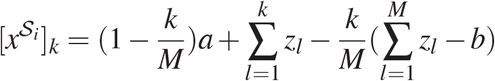

If a fragment terminates on the right (e.g there is no right endpoint), instead:

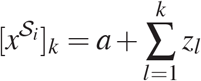

with left-terminating fragments handled similarly.

Once initialized, we integrate backwards in time the conditional probability flow ODE where the conditional 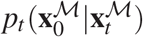 is taken be 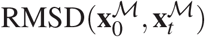 To address issues with clashes and discontinuities, we include terms: *p*_clash_(**x**_0_|**x**_*t*_) and *p*_violation_(**x**_0_|**x**_*t*_) where *p*_clash_ is a *L*1 penalty on all-atom distance matrix of **x**_*t*_ restricted to non-adjacent residues penalizing distances less than 1.5 *Å*., and *p*_violation_ is given by the violation loss defined in [Jumper et al., 2021].

## L Programmability: Symmetry

### L.1 Motivation

Built from identical subunit proteins, many protein complexes are assembled symmetrically. Many symmetric complexes such as tube-shaped channel proteins and icosahedral viral capsids are biologically important [Goodsell and Olson, 2000]. Incorporating symmetry in computational protein generation holds promise in designing large functionalized protein complexes [Hsia et al., 2016]. To fully explore the sampling of protein complexes subject to symmetry constraints, we propose a method to symmetrize the underlying ODE/SDE sampling to satisfy any prescribed Euclidean symmetries.

Incorporating group equivariance in machine learning has been an important topic in the machine learning community. [Cohen and Welling, 2016] Incorporating space group symmetries is critical in molecular simulations [Cox and White, 2022, Zabrodsky et al., 1992]. In this work, we proposed a method to incorporate symmetry for diffusion probabilistic models with applications in generating large-scale protein complexes with arbitrary symmetry groups.

### L.2 Symmetry breaking in sampling

Let 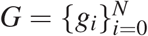 be a collection of symmetry operations that form a group such as point groups and space groups. For point sets in R^3^, these symmetry operations can be represented as a set of orthogonal transformations (rotation/reflection) and translations. The sampling SDE proposed in our work can be generally cast in the following form:

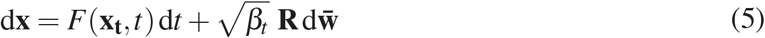

For synthesizing symmetric protein complexes, we want to sample complexes ***X***_*t*=0_ ∈ ℝ^*N*×*n*×3^ which are built from *N* = |*G*| identical single-chain proteins **X**^(i)^ ∈ ℝ^*n*×3^ where *n* is the number of residues for each subunit. Under the time-dependent noise 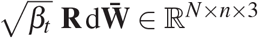, the samples are generated from:

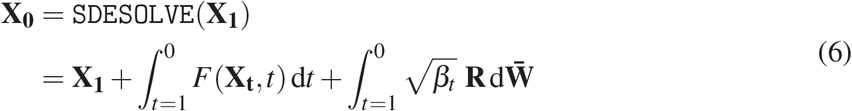

To constrain the sample generation to respect symmetries for an arbitrary group *G*, the SDE/ODE dynamics need to be *G*-invariant up to a permutation of subunits. Let · represent the symmetric operations (rotation, reflection, and translation) performed on point sets in R^3^, we define the sampling procedure `SDESOLVE`: ℝ^|*G*|×*n*×3^ → ℝ^|*G*|×*n*×3^ with *X*_0_ = `SDESOLVE`(*X*_1_) being the desired samples. The sampling procedure needs to follow the following invariance condition:

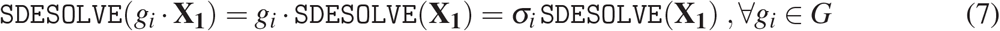

where *g_i_* indicates the *i*-th group element in *G* and we impose an arbitrary order on *G* and our method is equivariant to the permutation of subunits. *σ_i_* is the induced permutation operation satisfying the relation: *g_i_G* = *σ_i_G*, as computed from the group multiplication table (also called the Caley table).

The first equality in eq. (7) is trivially satisfied if *F*(·) is *E*(3) equivariant, as *G* consists of only orthogonal transformations and translations. However, the second equality is generally not satisfied. For molecular simulations where the Hamiltonian dynamics is used, the second equality can be satisfied if (i) the energy function is E(3) invariant, and (ii) the initial **X**_1_ and 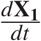 are symmetric, i.e 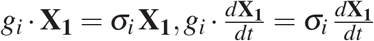. At each successive time step, *X_t_* automatically satisfies the prescribed *G*-symmetry. This approach confines both the position and momentum update to ensure the sampled configurations remain symmetric.

However, this is not the case with SDE/ODE sampling in our framework. We list three origins of symmetry-breaking error if eq. (6) is used: (i) *F*(**X**_t_, *t*) uses distances as features and is automatically *E*(3) equivariant. However, because the protein feature graphs are generated probabilistically, *F*(*g_i_* · **X**_*t*_, *t*) ≠ *g_i_* · *F*(**X**_t_, *t*) with each subunit protein *x^i^* having different geometric graphs, albeit being symmetric. (ii) Our polymer structured noise is randomly sampled from 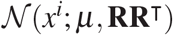, so each subunit protein has different chain noises, i.e. 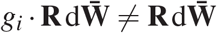. (iii) The sampling procedure requires solving an ODE/SDE which is vulnerable to accumulated integration error. Integration error can induce unwanted geometric drifts such as rotation and translation[Harvey et al., 1998], and be a substantial symmetry-breaking force.

### L.3 Symmetric sampling

By leveraging the desired symmetry, we averaged the SDE/ODE update from symmetric subunit proteins and broadcast with symmetry operation *G*. By performing the symmetric broadcasting, we remove the symmetry-breaking error for each symmetric subunit. The goal is to construct sampling protocols that satisfy eq. (7) which also capture interactions between subunits.

#### L.3.1 Symmetric initialization

For samples to remain *G*-invariant throughout the sampling process, it is necessary to have the initial noised structure **X**_1_ symmetrized. We define the following symmetrized “copying” operation:

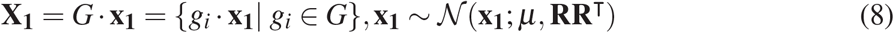

By construction, the generated structures are symmetric under *G*, i.e.

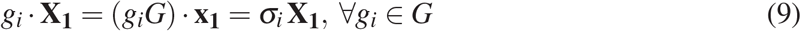

#### L.3.2 Symmetrized SDE

##### Trivial symmetrization

Given symmetrically initialized **X**_1_, a trivial construction is to set

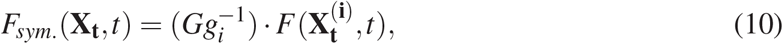

where *i* is the index to any subunit that is generated with *g_i_* and we dropped *t*-dependence in *F* for a more compact notation. Note that one does not need to make a special choice of 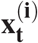 and *g_i_* as all subunits are symmetric. However, it is more convenient to select *i* where *g_i_* corresponds to the identity transformation. The method only requires performing an update on a single subunit, followed by symmetrized broadcasting. However, it satisfies eq. (7), because it fails to capture any subunit-subunit interactions which are important in capturing protein complexes.

##### Symmetric broadcasting

To incorporate subunit interactions, we use the entire symmetric complex as input for *F*(·,·) so that the subunit interactions are captured, and our backbone GNN is also designed to capture large protein complexes. Similar to the trivial construction described above, we select a particular *g_i_* and compute:

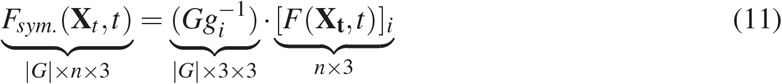

For noise, we simply sample polymer structured noise and broadcast:

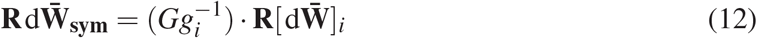

where [*F*(**X**_t_)]_*i*_ retrieves the gradient update for subunit *i*. Intuitively, the method computes the gradient update globally and broadcasts the update vector to all symmetric positions.

For simulating large complexes where certain long-range interactions are not as critical, we can instead focus on updating a subset 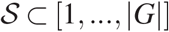 of subunits in **X**_t_ to save memory and time. Given a chosen subunit *i*, we deduce 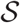 by choosing the k-nearest neighbor ((k-NN)) subunits based on the distances between the subunits’ geometric centers, so that short-range subunit interactions are included. We select *K* subunits in this way, and *K* is a hyperparameter.

##### Symmetric averaging

An alternative symmetrization method is to average the *F*(**X**_*t*_, *t*) at symmetric coordinates over all possible symmetry operations in *G*.

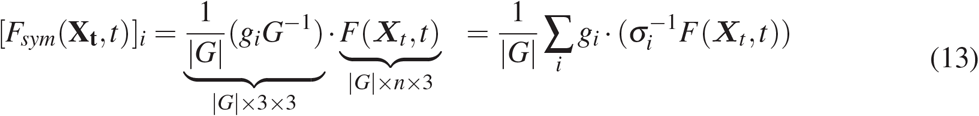

where (*g_i_G*^−1^) aligns the [*F*(**X**, *t*)]_*j*_ onto the symmetric subunit *i* and 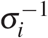 equivalently permute symmetric gradient update to argree with order used to generate the seed geometry *x*_1_. Then we average the symmetric *F(x, t*) contribution together to obtain [*F*(**X**, *t*)]_*i*_. In practice, one can just pick *i* to compute [*F*(**X**, *t*)]_*i*_ and broadcast to symmetric subunits via *G*. Similarly, the noise can also be symmetrized by averaging:

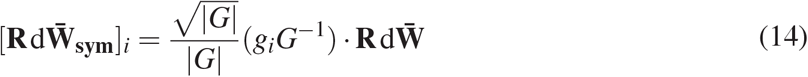

where 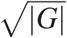 is used to correct the shrinking in the noise covariance because chain noise is obtained by averaging |*G*| symmetric subunits.

In summary, we proposed two methods of constructing *F_sym_*. (·) and **R***dW* and replace the sampling operation described in eq. (6) with the following modified SDE:

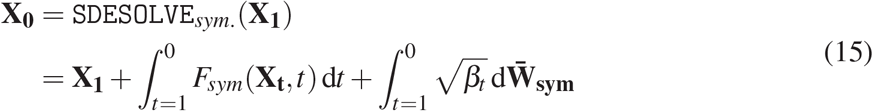

The high-level algorithm is described in Figure 11 with an example that illustrates the *C*_4_ symmetric sampling.

### L.4 Additional symmetric samples

We include more generated samples for selected point groups including *C_n_* (cyclic symmetry), *D_n_* (dihedral symmetry), *T* (tetrahedral symmetry), *O* (octahedral symmetry), *I* (icosahedral symmetry). For each all the samples we use *λ*_0_ = 16 and *ϕ* = 2 with the Heun SDE solver that integrates from 1 to 0 for 400 steps. We use subunit k-NN sampling with *K* = 6. When *K* > |*G*|, we set *K* = |*G*|. We provide additional samples categorized by the imposed symmetry group in Figure 12 with a range of sequence lengths per subunit. Our method strictly imposes symmetries. However, the sampled geometries can sometimes show poor contact while still being symmetric. We provide such samples in Figure 13.

**Supplementary Figure 11:**
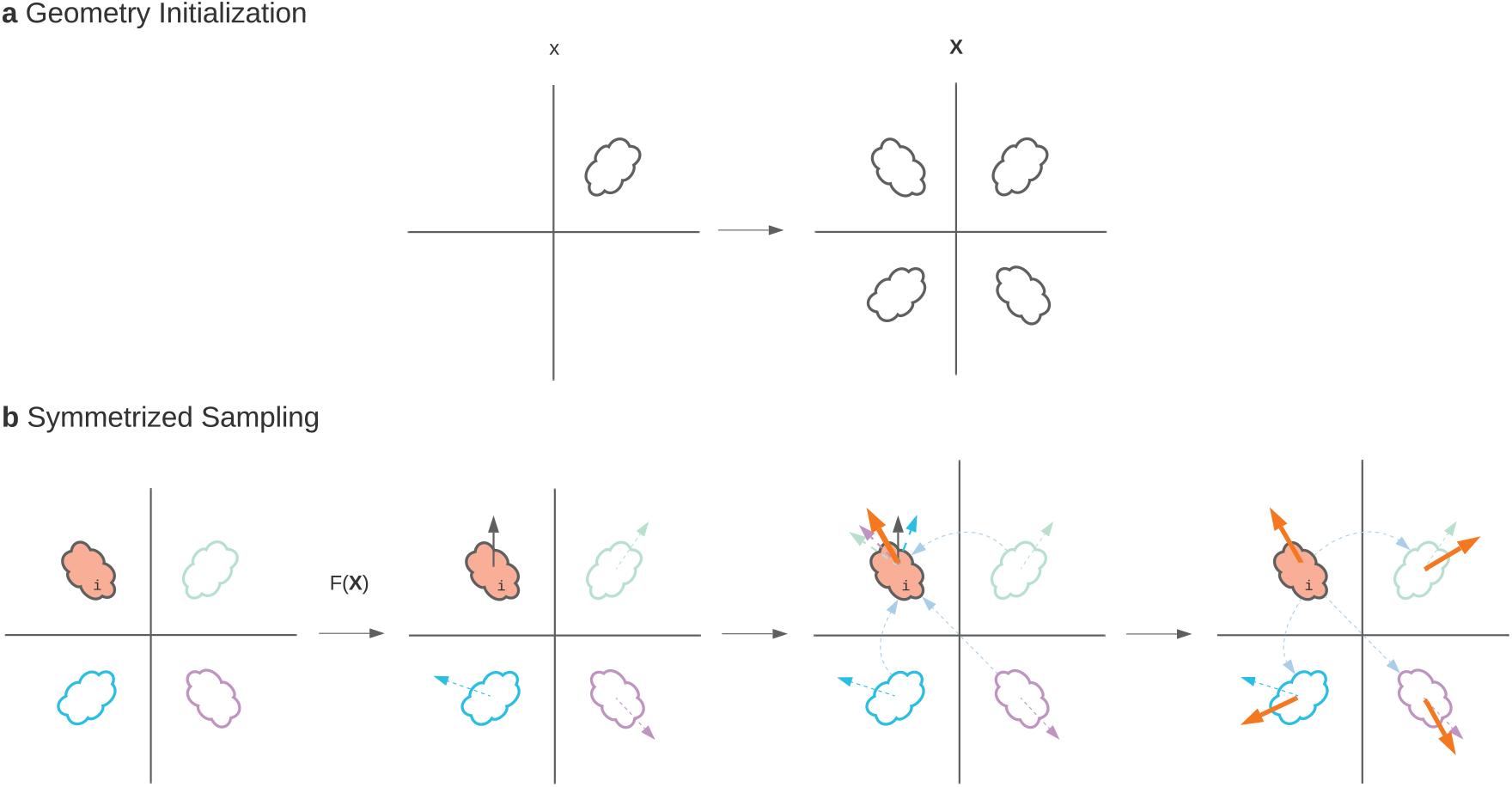
(**a**) The protein complexes are initialized by performing symmetry operations on an initial protein. (**b**) *F* (**X**) and 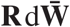 are symmetrized by averaging [*F*(**X**)]_*j*_ at symmetric positions.

## M Programmability: Shape

### M.1 Motivation

Proteins often realize particular functions through particular shapes, and consequently being able to sample proteins subject to generic shape constraints would seem to be an important tool for fully realizing the potential of protein design. Pores allow molecules to pass through biological membranes via a doughnut shape, scaffolding proteins spatially organize molecular events across the cell with precise spacing and interlocking assemblies, and receptors on the surfaces of cells interact with the surrounding world through precise geometries. Here we aim to explore and test generalized tools for conditioning on volumetric shape specifications with Chroma.

### M.2 Approach

Our shape conditioning approach is based on Optimal Transport [Peyré et al., 2019], which provides tools for identifying correspondences and geometric distances between objects, such as the atoms in a protein backbone and a point cloud sampled from a target shape. We leverage two metrics from the optimal transport theory: (i) the Wasserstein distance [Peyré et al., 2019], which can measure the correspondence between point clouds in absolute 3D space and (ii) the Gromov-Wasserstein distance, which can measure the correspondences between objects in different domains by comparing their intra-domain distances or dissimilarities. Because it leverages *relational* comparisons, Gromov-Wasserstein can measure correspondences between unaligned objects with different sturctures and dimensionalities such as a skeleton graph and a 3D surface [Solomon et al., 2016] or even between unsupervised word embeddings in two different languages [Alvarez-Melis and Jaakkola, 2018].

**Supplementary Figure 12:**
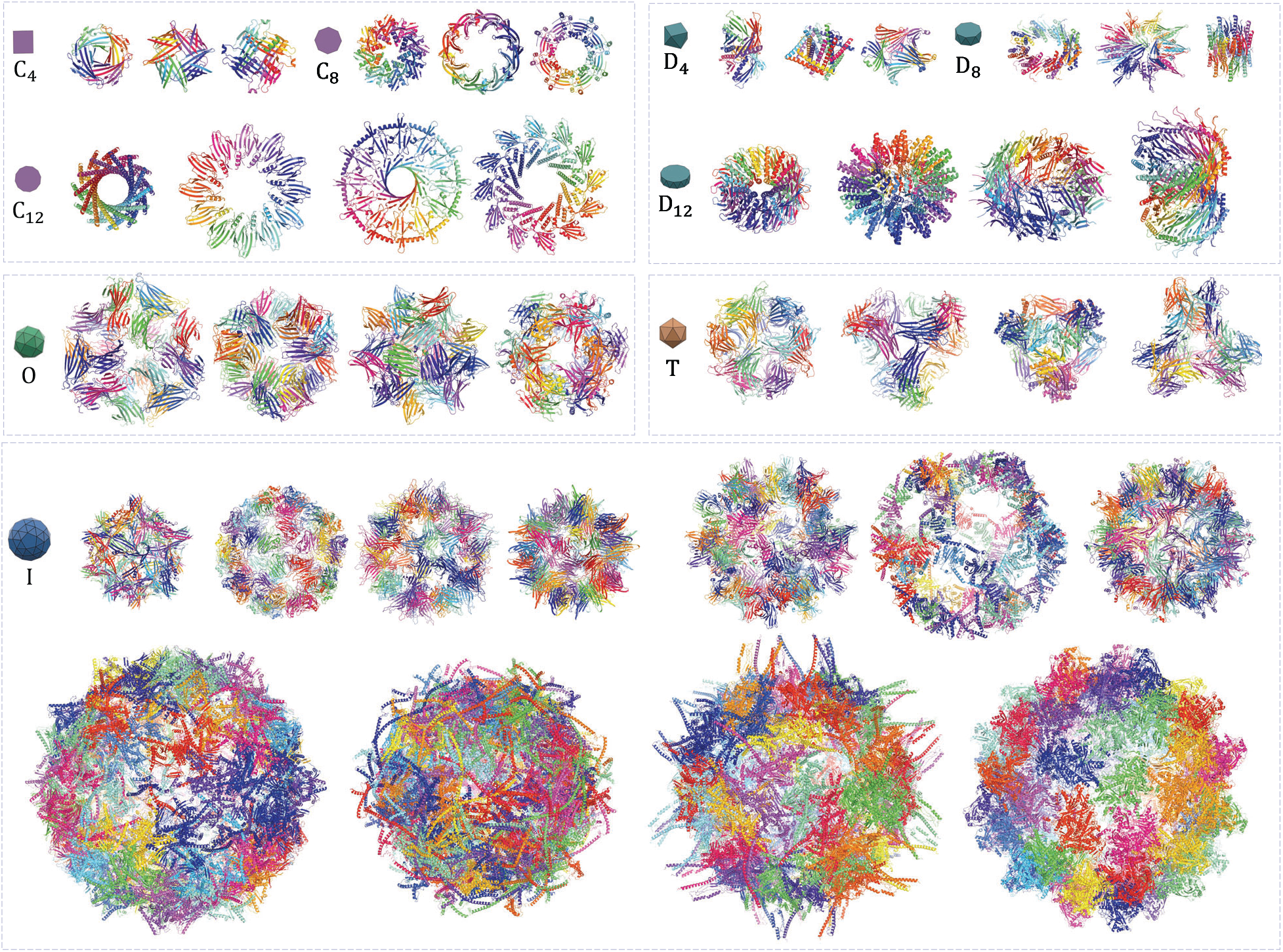
Additional generated complexes grouped based on imposed symmetry groups.

**Supplementary Figure 13:**
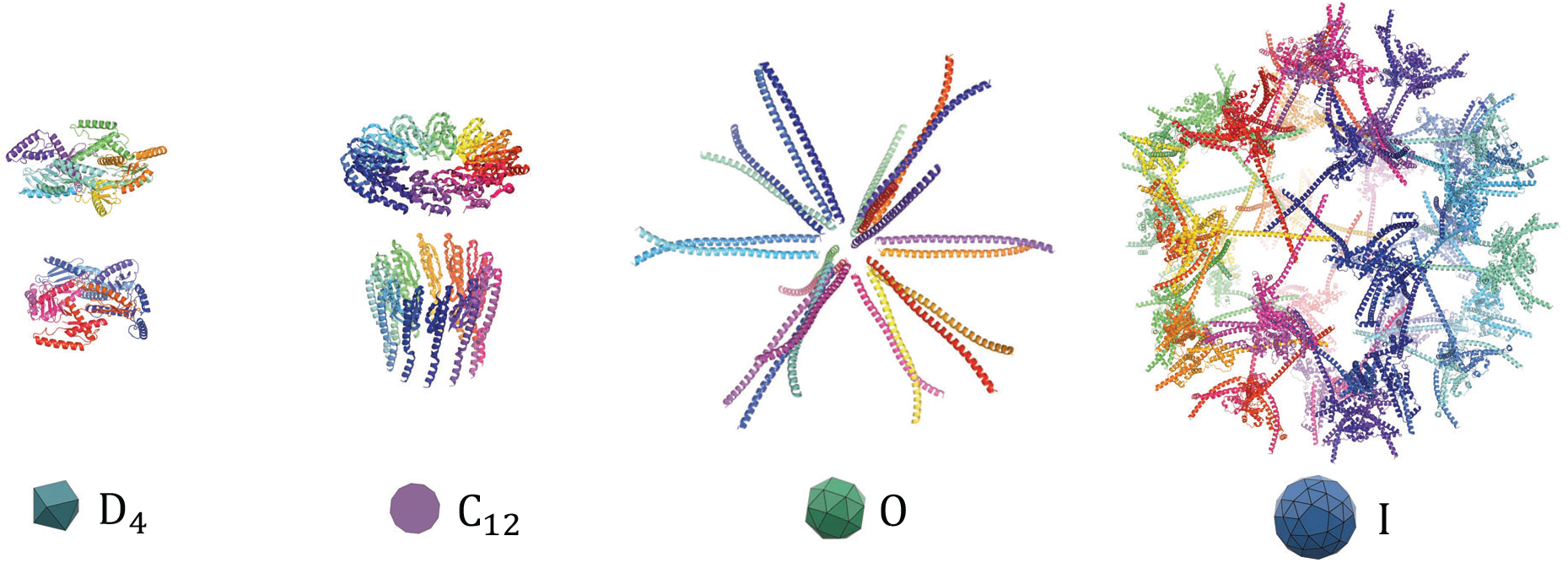
The generated complexes can form a poor protein-protein interface while still respecting imposed symmetry.

We initially experimented with adding heuristic gradients to the diffusion based on just the Wasserstein distance (estimated with the Sinkhorn algorithm [Peyré et al., 2019]), but found that the huge degeneracy in potential volume-filling conformations would often lead to jammed or high-contact-order solutions. While long-run Langevin sampling might help to allow gentle annealing into a satisfactory configuration in principle, we sought to accelerate convergence by breaking this degeneracy with a very coarse ”space-filling plan” for how the fold should map into the target point cloud, which the prior can then realize with a specific protein backbone.

#### Mapping 1D to 3D

We can leverage Gromov-Wasserstein (GW) optimal transport to answer the question “How would a protein with *ideal* distance scaling and a given length fill space in a target 3D volume?”. To do so, we (i) built an idealized distance matrix for a protein based on the scaling law^3^ *D_ij_* = 7.21 ×|*i* – *j*|^0.32^, (ii) compute the distance matrix for our target shape, and (iii) solve for the Gromov-Wasserstein optimal transport given these two distance matrices [Peyré et al., 2019] yielding a coupling matrix *K*_GromovWasserstein_ with dimensionality *N*_atoms_ × *N*_points_. This coupling map sums to unity and captures the correspondence between each atom in the abstract protein chain and each point in the target point cloud. We use a small amount of entropy regularization to solve the optimal transport problem.

#### Optimal Transport loss

In the inner loop of sampling, we can combine the GromovWasserstein coupling with simple Wasserstein couplings as a form of regularization towards our fold “plan”. Our final loss is then

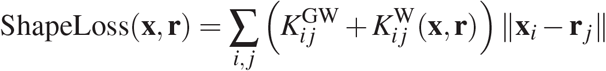

where we compute the Wasserstein optimal couplings 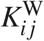 with the Sinkhorn algorithm [Peyré et al., 2019]. This yields a fast, differentiable loss that can be used directly for sampling.

#### Generating 3D shapes

We rendered letters and numbers from the English alphabet in the Liberation Sans font, extruded these 2D images into 3D volumes, and then sampled isotropic point clouds from these volumes.

## N Programmability: Residue, Domain, and Complex-level Classification

### N.1 Model Inputs

Noised backbone coordinates obtained from the PDB are passed as input to the model, along with a scalar 0 < *t* < 1 denoting the time during diffusion (indexed between zero and one) that the noise was sampled at. The model optionally can consume sequence information if available.

**Supplementary Figure 14:**
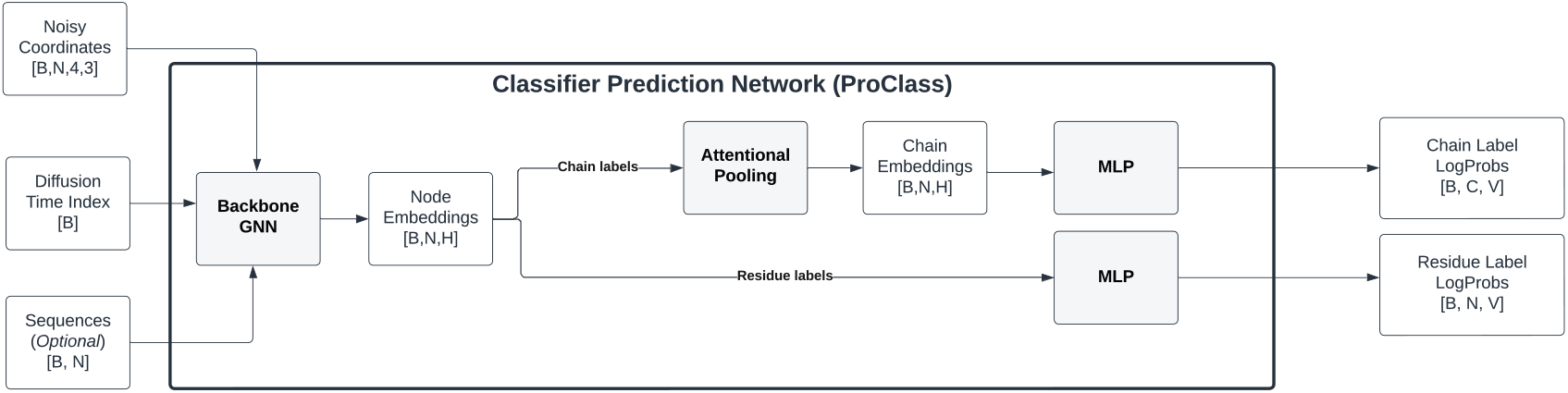
ProClass model architecture.

### N.2 Featurization

The time component is encoded with a random fourier featurization (e.g., see Tancik et al. [2020]). Provided sequence is encoded with a learnable embedding layer of amino acid identity. Backbone coordinates are passed to our `ProteinFeatureGraph` that extracts 2-mer and chain-based distances and orientations. These components are summed and passed to the neural network.

### N.3 Architecture

The encoder is a message passing neural network. The graph is formed by taking K=20 nearest neighbors and sampling additional neighbors from a distribution according to a random exponential method.

Node and edge embeddings are passed to each layer, with each node being updated by a scaled sum of messages passed from neighbors. The message passed from node *i* to node *j* is obtained by stacking the embeddings at node *i*, those at node *j*, and 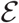, and passing these to a multi-layer perceptron (1 hidden layer). Edges are updated similarly. Each layer also applies layer normalization (along the channel dimension) and dropout (dropout probability=0.1).

After processing by the MPNN, node embeddings are passed to a different classification head for each label. If a head corresponds to a chain-level label, residues from each chain are pooled using an attentional pooling layer. The resulting embeddings are then passed to an MLP with 1 hidden layer to output logits for each label.

### N.4 Labels and loss functions

The model is trained to predict the following labels: CATH, PFAM, Funfam, Organism, Secondary Structure, Interfacial Residue. The loss for predicting each label is quantified using cross entropy loss, and all components are summed and weighted equally.

### N.5 Training

The model is trained for 50 epochs with an Adam optimizer [Kingma and Ba, 2014] with default momentum settings (betas=(0.9,0.999)), the learning rate is linearly annealed from 0 up to 0.0001 over the first 10,000 steps then kept constant. During training, first a time stamp 0 < *t* < 1 is sampled uniformly, then noise is sampled from the globular covariance distribution, injected into the backbone coordinates, and fed to the model. Next, label predictions are made, loss are computed, and parameters are updated with the Adam optimizer.

### N.6 Hyperparameters

The classification model has 4 layers, the size of node feature dimension is 512 and the edge feature dimension is 192, node update MLP has hidden dimension 256 with 2 hidden layers, and edge update MLP has hidden dimension 128 with 2 hidden layers.

## O Programmability: Natural Language Annotations

### O.1 Motivation

Recent advances in text-to-image diffusion models such as DALL-E 2 [Ramesh et al., 2022] and Imagen [Saharia et al., 2022] have produced qualitatively impressive results using a natural language interface. Given the open availability of pre-trained language models and a corpus of protein captions form large scientific databases such as the PDB [Berman, 2000] and UniProt [Consortium, 2020], we explore the possibility of creating a natural language interface to protein backbone generation. To do this, we build a protein captioning model (ProCap), which predicts *p*(*y*|**x**_*t*_), where *y* is a text description of a protein and **x**_*t*_ is a noised protein backbone. This conditional model, when used in conjunction with the structural diffusion model presented in the main text, can be used as a text to protein backbone generative model.

### O.2 Dataset curation

To build a caption model, we begin by curating a paired dataset of protein structures and captions from both the PDB and UniProt databases. Caption information is collected for the structures used for the backbone diffusion model training, as well as the individual chains within these structures. For each structure, we use the PDB descriptive text as an overall caption. For each chain in a structure, we obtain a caption by concatenating all available functional comments from UniProt. Structures containing more than 1000 residues are not used, corresponding to a minority (10%) of all structures. The final set used to train and validate the caption model contains approximately 45 thousand captions, including those from both PDB and UniProt. Unlike for the backbone model, the splits used for training are completely random. The small size of the dataset constrained architecture choices to those with relatively few free parameters.

### O.3 Model architecture

#### O.3.1 Architecture overview

To predict captions given noised structures, we construct ProCap using a pretrained language model and a pretrained protein encoder. The pretrained language model is the GPT-Neo 125 million parameter model [Black et al., 2021]. GPT-Neo was trained on the Pile [Gao et al., 2021] which contains articles from arXiv and Pubfed. Its choice is motivated to maximize the chance that the model would begin training with some understanding of protein-related text. We also use the pretrained graph neural network encoder from ProClass, the protein structure classification model introduced above, to encode protein backbones. Analogously to the choice of the language model, the purpose of the structure encoder is to start ProCap with semantic knowledge of protein structure. To condition the autoregressive language model, GPT-Neo, pseudotokens are formed from structures using the ProClass encoder and prepended to the caption as context, similar to [Lester et al., 2021].

**Supplementary Figure 15:**
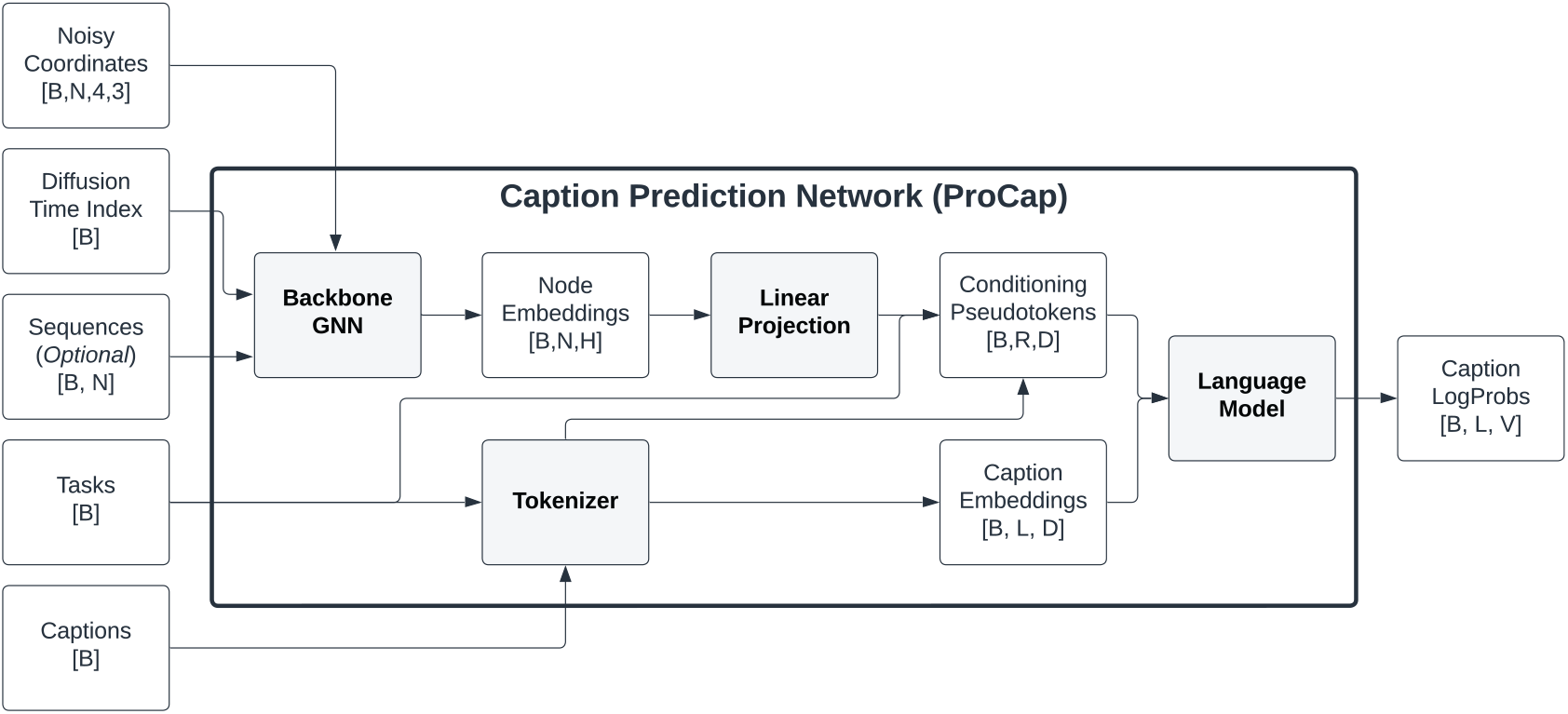
ProCap model architecture. ProCap connects a pretrained graph neural network encoder to an autoregressive language model trained on a large data corpus including scientific documents. We use the 125M parameter GPT-Neo as the language model, with internal dimension *D* = 768. Conditioning is achieved with pseudotokens generated from encodings of protein complex 3D backbone coordinates (batch size *B*, number of residues *N*, embedding dimension *H*) and a task token indicating whether a caption describes the whole complex or a single chain. The *R* relevant pseudotokens for each caption, consisting of the chain/structure residue tokens and the task token, are passed to the language model along with the caption. When used in the forward mode, ProCap can describe the protein backbone by outputting the probabilities of each word in the language model’s vocabulary of size *V* for each of the *L* tokens of a caption. When used in conjunction with the prior model, it can be used for text to protein backbone synthesis. In training, ProCap uses a masked cross entropy loss applied only to the caption logits.

#### O.3.2 Data embedding

Here, we describe the embedding of task, caption, and structure data into a shared tensor representation for input to the language model. Captions and task tokens are encoded using a modified version of the GPT-Neo tokenizer, whose vocabulary we augment with a special token to distinguish between prediction tasks involving single chains and those relating to entire structures. Structure inputs are converted into pseudotokens with the same shape as text embeddings through the graph neural network encoder of the pre-trained ProClass model. The task, structure, and caption embeddings are concatenated into a representation that is passed to the language model to obtain logits representing the probabilities of caption tokens. The model is trained on a standard masked cross entropy loss of the caption. The overall architectural flow is detailed in Fig. 15. We proceed to discuss the details of the embedding procedure.

Structure encoding in ProCap relies on a pretrained ProClass model. This classifier model consists of a GNN with multiple heads to extract different class information, as described previously. The GNN portion of the classifier network is used to obtain embeddings of each residue in the latent space of the classifier, with the intent that the pre-trained classifier weights should help ProCap learn the relationship between structures and captions. Besides the 3D information of the atoms in each structure, the diffusion timestep (noise level) is input to the GNN via a Fourier featurization layer which converts the diffusion time to a vector with the same dimension as the GNN node embedding space using randomly chosen frequencies between 0 and 16. To allow for ProCap to learn the optional use of sequence information, in 25% of the training data sequences are randomly passed along with structures. In these cases, the amino acid information for each residue is converted through a single embedding layer with output size equal to that of the GNN node embedding space dimension, then added to the time step vector.

Task tokens are added to the model to allow for captions of both single chain and full complex captions. For the prediction of UniProt captions describing single chains within structures, only the embeddings of the residues in the relevant chain are passed to the language model. For the prediction of the PDB captions related to entire structures, all residue embeddings are passed. In addition, a linear layer is added after the ProClass embeddings to go between the ProClass latent space and the embedding space of the language model, which are of different dimensionality. Finally, in order to help the model distinguish between PDB and UniProt prediction tasks, the encodings of the entire structures are each prepended with an embedding vector of a newly defined PDB marker token. We normalize the components of all structure vectors such that each one has zero mean and unit variance.

In summary, the ProCap architecture consists of a pre-trained GNN model for structure embedding and a pre-trained language model for caption embedding, with a learnable linear layer to interface between the two and a learnable language model head to convert the raw language model outputs to token probabilities.

### O.4 Model training

We train ProCap to be compatible with conditional generation using the structural diffusion prior model. Like the other conditional models in this paper, each structure is noised according to the schedule of the structural diffusion model. During ProCap training, the graph neural network encoder weights from the pre-trained ProClass model are frozen. In addition, the internal weights of the GPT-Neo language model are also frozen, except for the head whose parameters are allowed to train. We choose to freeze these model parameters because of the relatively small training data size compared to that which was used to pre-train the language model. The language model head is allowed to learn, both to improve the embeddings of its usual tokens, as well as to optimize the encoding of new tokens. In training, we add a <|`PDB`|> task token to the GPT-Neo vocabulary to cue the model to predict whole complex captions from the PDB.

Training is conducted on a single V100 with a constant learning rate of 5 × 10^−5^ and the Adam optimizer with hyperparameters *β*_1_ = 0.9, *β*_2_ = 0.999. We evaluate loss on a validation set after every 2000 training examples. During training, we set an early stopping patience of 40 iterations, which was triggered after approximately 20 epochs at a cross entropy loss of 3.29.

### O.5 Performance

In order to test ProCap as a generative model, we draw high-quality conditional and corresponding unconditional low-temperature samples from the model. To that end, we employ a structural denoising approach in a similar fashion to the method described in [Song et al., 2021]. Specifically, the hybrid Langevin-reverse time SDE of Appendix B is used to evolve noisy random sample structures drawn from the diffusion model prior, with gradients of the ProCap loss with respect to structure added to the gradients of the structure diffusion model. When the size of the ProCap gradients is too small relative to those from the prior model, there is little appreciable difference between a caption-conditioned sample and an unconditional sample drawn from the same seed. We thus scale the ProCap gradients up by a factor of 100 and find that the resulting samples are better conditioned, analogously to previous work on classifier guidance [Dhariwal and Nichol, 2021]. Simultaneously, we observe that the sample quality decreases as ProCap gradients are scaled up further, resulting in the loss of secondary structure and even breakdown of backbone bond length constraints. To mitigate this effect, we limit the size of the gradient,

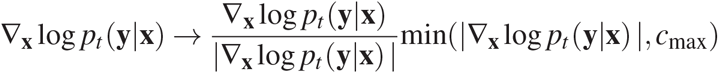

with the choice *c*_max_ = 10.

Examples of our generated samples are presented in the right two columns of Fig. 5. To evaluate ProCap model performance, we measure the improvement in caption loss during the SDE evolution between the unconditioned and conditioned samples. As an independent check, we also examine the gain in the TM-score between our sample (conditioned over unconditioned) and a target PDB structure which exemplifies the caption being used for conditioning. Finally, we analyze the generated structures visually for structural coherence. Qualitatively, starting from the same noisy random structure, the diffusion model yields denoised structures which demonstrate desirable characteristics including secondary structure elements, both with and without guidance from the caption model.

The caption loss and TM-score metrics for the sampling trajectories leading to the structures in Fig. 5 are shown in Fig. 16. Both are initially quite noisy, and the conditioned and unconditioned samples are equally likely at high *t* to have lower ProCap loss and/or better alignment with the target structure. However, over the course of the reverse diffusion, the effect of the conditioning is demonstrated in both panels. It is particularly notable that the TM-score is relatively stable at low *t*, indicating a regime where the SDE evolution is fine-tuning structural details rather than making significant changes.

**Supplementary Figure 16:**
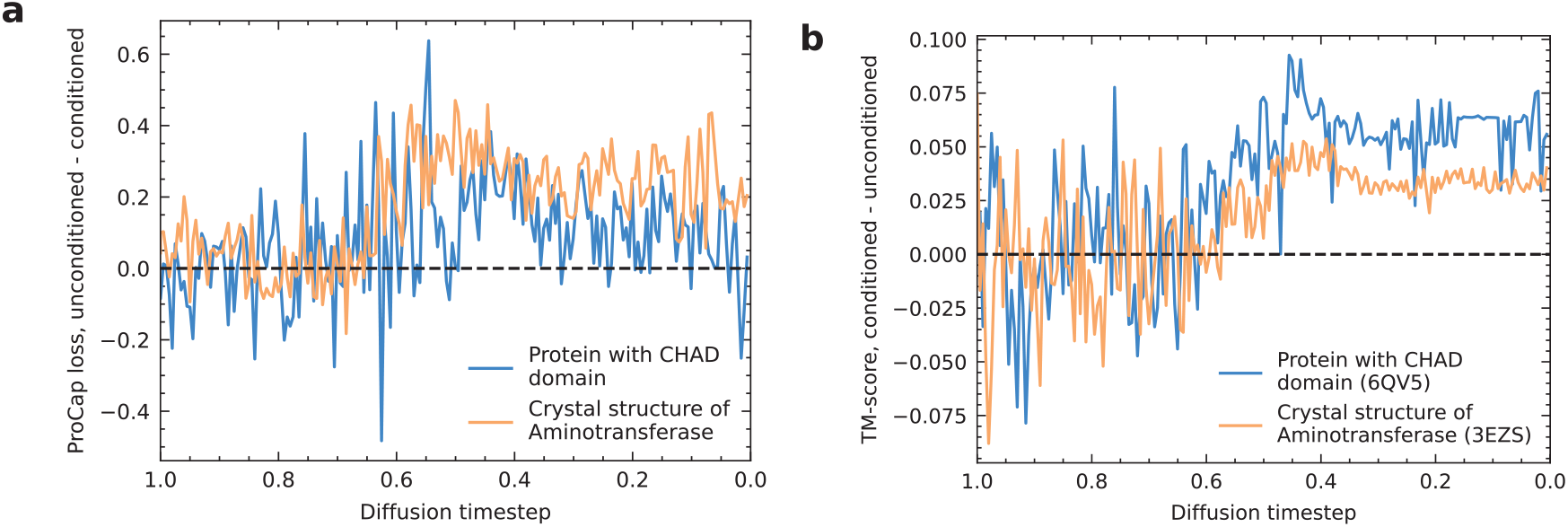
ProCap evaluation metrics show effect of natural language conditioning compared to unconditioned samples from the same noised seed structure. Panel **a** shows the caption model cross-entropy loss as a function of diffusion timestep, for two sample trajectories with and without the use of caption gradients. Panel **b** shows the TM-score between sampled structures and example structures from the PDB corresponding to the captions used for conditioning.

## Footnotes

1 We will assume the data are centered (have zero mean) for ease of notation.

2 In some of our symmetry examples we find that models still generalize well to systems larger than they were trained on.

3 This scaling law was fit on a large single-domain protein 6HYP.

## Notes

### Competing Interest Statement

The authors have declared no competing interest.

